# A taxon-restricted duplicate of *Iroquois3* is required for patterning the spider waist

**DOI:** 10.1101/2023.08.08.552527

**Authors:** Emily V. W. Setton, Jesús A. Ballesteros, Pola O. Blaszczyk, Benjamin C. Klementz, Prashant P. Sharma

## Abstract

The chelicerate body plan is distinguished from other arthropod groups by its division of segments into two tagmata: the anterior prosoma (“cephalothorax”) and the posterior opisthosoma (“abdomen”). Little is understood about the genetic mechanisms that establish the prosomal-opisthosomal (PO) boundary. To discover these mechanisms, we created high-quality genomic resources for the large-bodied spider *Aphonopelma hentzi*. We sequenced specific territories along the antero-posterior axis of developing embryos and applied differential gene expression analyses to identify putative regulators of regional identity. After bioinformatic screening for candidate genes that were consistently highly expressed in the posterior segments, we validated the function of highly ranked candidates in the tractable spider model *Parasteatoda tepidariorum*. Here, we show that an arthropod homolog of the Iroquois complex of homeobox genes is required for proper formation of the boundary between arachnid tagmata. The function of this homolog had not been previously characterized, because it was lost in the common ancestor of Pancrustacea, precluding its investigation in well-studied insect model organisms. Knockdown of the spider copy of this gene, which we designate as *waist-less*, in *P. tepidariorum* resulted in embryos with defects in the PO boundary, incurring discontinuous spider germ bands. We show that *waist-less* is required for proper specification of dorso-ventral identity in the segments that span the prosoma-opisthosoma boundary, which in adult spiders corresponds to the narrowed pedicel. Our results suggest the requirement of an ancient, taxon-restricted paralog for the establishment of the tagmatic boundary that defines Chelicerata.

## Introduction

Functional understanding of the evolution of animal body plans is frequently constrained by two bottlenecks. First, developmental genetic datasets and functional toolkits are often asymmetrically weighted in favor of lineages that harbor model organisms, to the detriment of phylogenetically significant non-model groups. Second, models of ontogenetic processes that are grounded in model systems vary in their explanatory power across diverse taxa, both as a function of phylogenetic distance, as well as the evolutionary lability of different gene regulatory networks (GRNs) [1–4]. In Arthropoda, understanding of morphogenesis, as well as the evolutionary dynamics of underlying GRNs, is largely grounded in hexapod models, and particularly holometabolous insects. Candidate gene approaches derived from studies of insect developmental genetics have thus played an outsized role in understanding of the mechanisms of arthropod evolution, with emphasis on processes like segmentation, limb axis patterning, and neurogenesis [5–10]. However, the candidate gene framework has its limits in investigations of taxon-specific structures (e.g., spider spinnerets; sea spider ovigers) [11–13], or when homologous genes or processes do not occur in non-model taxa (e.g., *bicoid* in head segmentation) [7,14].

These limits are accentuated in Chelicerata (e.g., spiders; scorpions; mites; horseshoe crabs), the sister group to the remaining arthropods. The bauplan of most chelicerates consists of two tagmata, the anterior prosoma (which bears the eyes, mouthparts, and walking legs) and the posterior opisthosoma (the analog of the insect abdomen). Even at this basic level of body plan organization, differences in architectures are markedly evident between chelicerates and the better-studied hexapods. The chelicerate prosoma typically has seven segments and includes all mouthparts and walking legs, whereas the insect head has six segments and bears only the sensory (antenna) and gnathal appendages (mandibles, maxillae, labium); locomotory appendages of insects occur on a separate tagma, the thorax [15].

Comparatively little is known about how these functional groups of segments are established in chelicerates, by comparison to their insect counterparts. Due to the phylogenetic distance between hexapods and chelicerates, homologs of insect candidate genes that play a role in tagmosis can exhibit dissimilar expression patterns or incomparable phenotypic spectra in gene silencing experiments in spiders, a group that includes the leading models for study of chelicerate development [13,14,16–18]. A further complication is the incidence of waves of whole genome duplications (WGDs) in certain subsets of chelicerate orders, such as Arachnopulmonata, a group of six chelicerate orders that includes spiders [19–22]. The retention of numerous paralogous copies that diverged prior to the Silurian represents fertile ground for understanding evolution after gene duplication, but also presents the potential barrier of functional redundancy or replacement between gene copies. Accordingly, there are few functional datasets supporting a role for lineage-specific gene duplicates in the patterning of arachnid body plans [23,24].

To advance the understanding of chelicerate body plan patterning and address possible roles for retained paralogs in chelicerate tagmosis, we generated transcriptional profiles of prosomal and opisthosomal tissues of a large-bodied spider (a tarantula), across developmental stages pertinent to posterior patterning. We applied differential gene expression (DGE) analyses to triangulate taxon-specific gene duplicates that were differentially expressed across the prosomal-opisthosomal (PO) boundary and screened candidates using an RNA interference (RNAi) gene silencing approach. Through this approach, we were able to identify one of the five spider paralogs of *araucan*/*caupolican* (*ara*/*caup*; *Iroquois4 sensu* [25]; *Iroquois3-2*, *sensu* [26]) as playing a role in dorso-ventral (D-V) patterning of the segments spanning the PO boundary. Our results provide a functional link between an unexplored gene copy restricted to non-pancrustacean arthropods and the boundary between the tagmata of chelicerates.

## Results

### Differential gene expression, RNAi screen, and identification of waist-less

To understand the genetic basis of posterior patterning in spiders, we aimed to generate tissue-specific transcriptomes of spider embryos. The leading model system for spider development, *Parasteatoda tepidariorum*, proved challenging in this regard, due to the small size of its embryos (500 μm) and the high internal pressure of the egg. We therefore generated differential gene expression datasets for the tarantula *Aphonopelma hentzi*, which features large and synchronous broods, and embryos with large diameter (2.4 mm) and low internal pressure [27]. We dissected clutches of synchronously developing tarantula embryos and generated RNA-seq libraries for segments bearing the labrum, chelicera, pedipalp, walking leg, book lung, anterior spinneret, and posterior spinneret. This protocol was performed for three developmental stages, encompassing establishment and differentiation of posterior appendages (e.g., book lungs and spinnerets) [27]. Differential gene expression (DGE) analysis identified genes 5,429-14,094 (stage 9: 7,609; stage 10: 5,429; stage 11: 14,094) as consistently differentially expressed across segments in an all-versus-all comparison (p ≤ 0.05; LFC≥ 1) (Fig. 1A). To triangulate genes that may play an important role in posterior patterning, we assessed the top 100 most differentially expressed genes for each developmental stage, as well as examined comparisons of specific tissue pairs, and screened candidates that were (1) consistently highly expressed in opisthosomal segments in at least two stages, and (2) consistently lowly expressed in prosomal segments in at least two stages (stage 9: 67; stage 10: 53; stage 11: 92) (Fig. S1). We prioritized 16 genes for functional screening (SI Appendix, Table S1). Colored circles correspond to different orthologs, following F. Boldface text indicates spider *waist-less* orthologs. Inset: Full unrooted gene tree of Iroquois homologs. F. Inferred evolutionary history of *Iroquois* gene duplications in Chelicerata. Scale bar: 100 μm.

**Figure 1.**
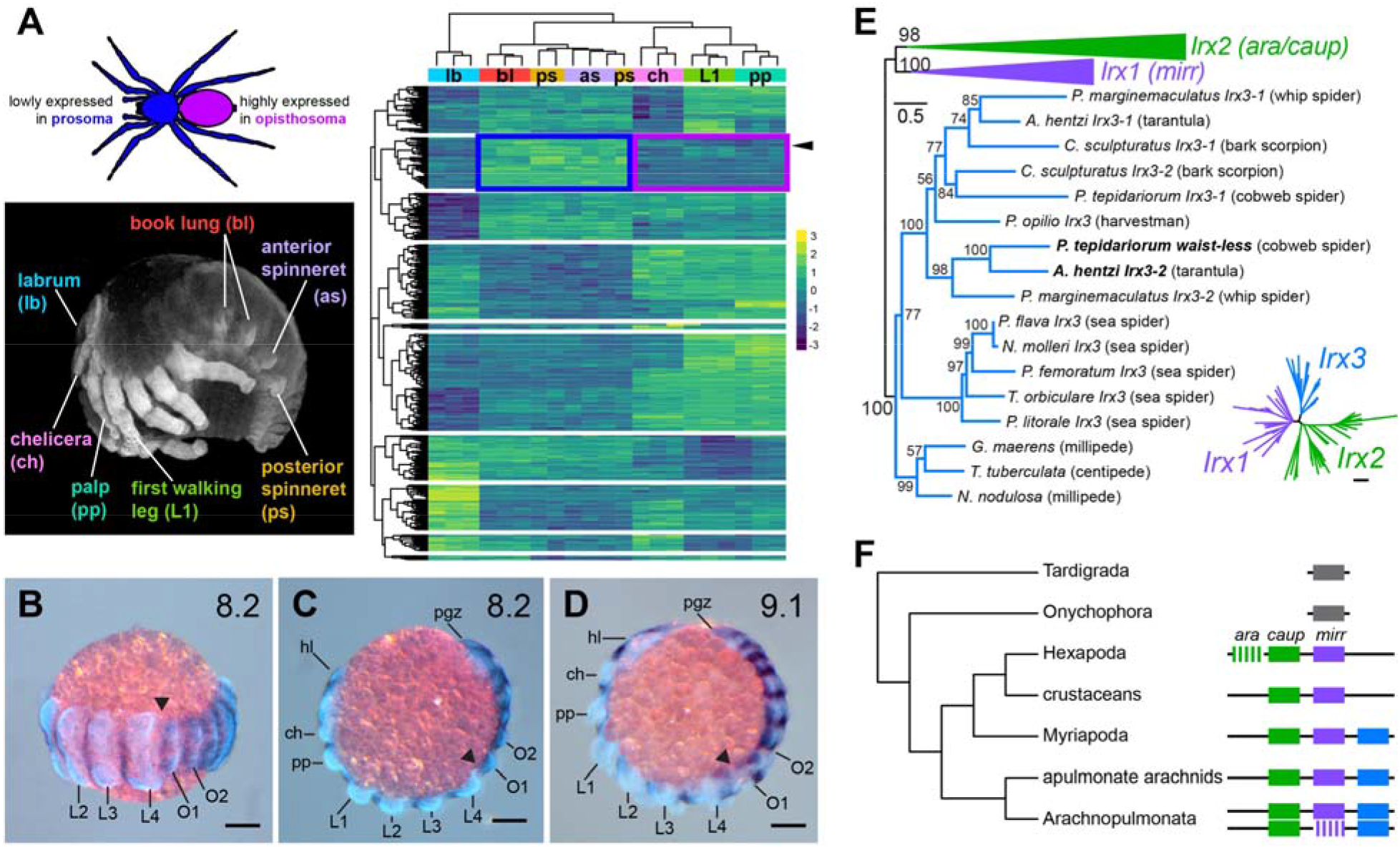
Overview of RNAseq design, candidate gene identification, and the ortholog identification within the *Iroquois* gene family. A. Tissues from regions representing major morphological characters along the anterior-posterior (A-P) axis were dissected from developing *Aphonopelma hentzi* embryos for mRNA sequencing. Differential gene expression (DGE) analysis of RNAseq libraries generated region-specific profiles to enable the identification of genes both lowly expressed in the prosoma (blue box) and highly expressed in the opisthosoma (purple box). Arrowhead indicates the ortholog of spider *waist-less*. B-D. Expression of *waist-less* in limb bud stage embryos of *Parasteatoda tepidariorum*, counterstained for Hoechst. Note the higher expression level in the opisthosoma compared to the prosoma. E. Maximum likelihood gene tree of *Iroquois2/3* homologs of Panarthropoda, rooted on Onychophora.

Due to the lack of gene silencing tools in the tarantula, we performed functional screening of candidate genes in the house spider *Parasteatoda tepidariorum*, following established protocols [17,28–30]. Of the 16 candidates, 14 yielded no discernable phenotype, paralleling outcomes of recent RNAi screens in this system [13]. One candidate that yielded a consistent phenotype was annotated as a member of the Iroquois complex of homeobox genes (Fig. 1B-D). Previously identified as “*Iroquois4*” in a recent survey of homeobox family duplications [25], this transcription factor is not orthologous to the identically named vertebrate homolog Iroquois4, nor is its homology to its two insect homologs (*mirror* and *arucan/caupolican*) understood [31]. To forfend redundancy of nomenclature within the chelicerate Iroquois complex, we rename the differentially expressed spider copy (previously, “*Iroquois4*”) *waist-less* (*wsls*), reflecting the phenotypic spectrum described below.

### Evolutionary history of panarthropod Iroquois homologs

To better understand the evolutionary history of this gene in arthropods, we inferred a gene tree of the Iroquois family, surveying genomes of four arachnopulmonates (arachnids that share a whole genome duplication; two spiders, a whip spider, and a scorpion), six non-arachnopulmonate chelicerates (chelicerates with an unduplicated genomes; five sea spiders and a harvestman), four myriapods (sister group to chelicerates with unduplicated genomes; two centipedes, two millipedes), three crustaceans, and 12 hexapods. The gene tree topology (Fig. 1E; Fig. S2) recovered *Iroquois1*, *Iroquois2*, and *Iroquois3* homologs as three separate clusters, with maximal nodal support for *Iroquois3*. Whereas exemplars from all major arthropod lineages bore *Iroquois1* and *Iroquois2* homologs, the cluster corresponding to *Iroquois*3 was comprised only of myriapod and chelicerate exemplars (Fig. 1E).

To polarize the evolutionary history of the Iroquois complex, we examined the organization of Iroquois homologs in well-annotated genomes of Panarthropoda (Fig. 1F; SI Appendix, Table S2). Whereas a single Iroquois homolog occurs in high-quality genomes of Tardigrada and Onychophora, chromosomal-level genomes of Myriapoda and apulmonate Chelicerata exhibited three Iroquois homologs arranged contiguously on single scaffolds, consistent with an origin of the arthropod Iroquois genes via two tandem duplications. Chromosomal-level genomes of spiders recovered five to six Iroquois copies, with homologs of *Iroquois1*, *Iroquois2*, and *Iroquois3* occurring on two separate scaffolds, consistent with whole genome duplication in the arachnopulmonate common ancestor. The ancestral arrangement of the three Iroquois homologs was observed to be reordered in one cluster in the spider *Dysdera sylvatica* (see also [26]. In support of this result, non-arachnopulmonate chelicerates (e.g., the harvestman; sea spiders) bore three Iroquois homologs in the gene tree (one homolog of *mirror*, one of *araucan/caupolican*, and one of *Iroquois3*), whereas spiders and scorpions bore up to six Iroquois homologs due to an arachnopulmonate-specific whole genome duplication. *P. tepidariorum* bore only five Iroquois homologs due to the loss of one *mirror* copy (Fig. S2).

The absence of *Iroquois3* in all sampled exemplars of hexapods and crustaceans is consistent with a loss of this gene in the branch subtending Pancrustacea. Additionally, the duplication and subdivision of *Iroquois2* into *araucan* and *caupolican* is limited to a subset of flies (e.g., *D. melanogaster*), not all Diptera (e.g., *Calliphora vicina*; *Anopheles gambiae*; *Culex pipiens quinquefasciatus*) (Fig. 1F).

### Expression of the waist-less ortholog in Parasteatoda tepidariorum

Expression of spider *Irx4* was previously reported for selected stages of development and a segmentation function had been suggested due to the segmentally reiterated stripes of expression [25,26,32]. We first surveyed *waist-less* expression across the embryogenesis of *P. tepidariorum*. Expression was initially detected at stage 6 as stripes corresponding to the segments of the germ band (Fig S3A). Segmentally repeated bands of expression persist throughout development (Fig S3). At stages 8 and 9, expression is notably stronger in opisthosomal segments compared to prosomal segments, due to the incidence of weaker domains bridging the segmentally iterated stripes of *waist-less* in the opisthosoma (Fig. 1B-1D), and corroborating the stronger posterior expression predicted by DGE. Additional expression domains include the a “V” shape in the anterior head beginning at stage 8.2 (Fig S3D-F). At stage 9.2 expression appears in the lateral body wall, together with a distinct, distal point of expression in the prosomal appendages. At this stage, the “V” of expression in the anterior head becomes a pair of arcs, curved inward toward each other approaching the ventral midline and comprising the medial head region that lies anterior to the cheliceral limb buds (Fig S3G-I). At stage 10.1 the crescents of expression on each side of the developing head become more concentrated. The segmentally repeated stripes of expression are still maintained ventrally, but with the stripes no longer of uniform strength across the germ band. At this stage, increased expression is seen in the opisthosomal appendages (Fig S3J-L). Gene expression at stage 10.2 expression is similar to 10.1, but with increased localization to the lateral margins and appendage primordia of the opisthosoma (Fig S3M-O). Stage 11 embryos exhibit increased division of the stripes across the width of the germ band and continued strong expression in the lateral part of the opisthosoma (Fig S3P-R).

### Knockdown of *waist-less* disrupts the prosoma-opisthosoma boundary in a spider

To assess the function of *waist-less*, RNA interference (RNAi) was performed using established protocols [17,28–30]. Parental RNAi against *Ptep-waist-less* via maternal injections of dsRNA resulted in a phenotypic spectrum affecting the PO boundary. Validation of knockdown was assessed using colorimetric *in situ* hybridization (Fig. S4). Phenotypes were scored in embryos stage 8.1 or later, when morphological landmarks are present, and designated into two classes. Class I phenotypes (17.2%; n = 41) exhibited a PO boundary defect, consisting of reduction of the first opisthosomal and the fourth walking leg segments (Fig. 2E). In later stages, the embryo developed as a discontinuous germ band, with no embryonic tissue in the region corresponding to the posterior prosoma and the anterior opisthosoma (Fig. 2F). Class II phenotypes (31.9%; n = 76) exhibited reduction of embryonic tissue spanning the anterior opisthosoma up to the middle of the prosoma (walking leg II) (Fig. 2G). Class II phenotypes also exhibited a discontinuity at the boundary between tagmata (Fig. 2H), but additional defects observed in the prosoma included fusion of adjacent limb buds, bifurcated pedipalps, and smaller chelicerae (Fig. 2G, 2H). Few embryos exhibiting *Ptep-waist-less* phenotypes completed development; in later stages of embryogenesis, a small number of embryos was observed with discontinuous prosoma and opisthosoma (n = 3/55) (Fig. S5D, S5E). In a separate RNAi trial, we observed three *Ptep-waist-less* RNAi postembryos, which exhibited the mildest phenotypic defect (truncated L4 segment on one side of the body; n = 3/59) (Fig. S5F).

**Figure 2.**
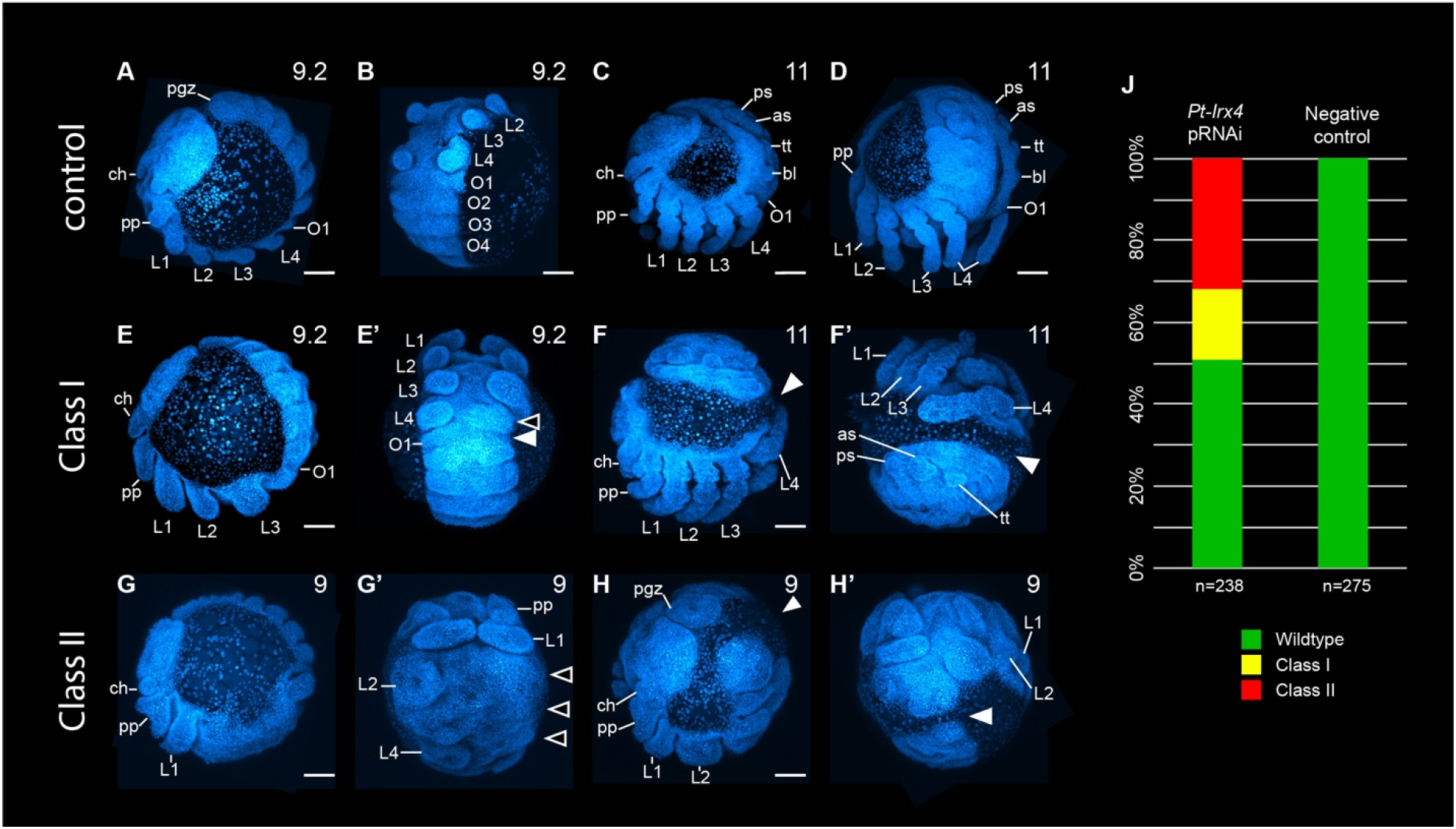
Phenotypic spectrum of *Ptep-waist-less* maternal RNAi. **A-D.** Wild type development of *P. tepidariorum* in negative control experiments. E-F’. Class I *Ptep-waist-less* RNAi embryos exhibit reduction or loss of L4 segment (E’ open arrowhead) or disruption of both L4 and anterior opisthosomal segments (E’ solid arrowhead). Some Class I embryos also exhibit discontinuous germ bands (F-F’ solid arrowhead). G-H’. Class II *Ptep-waist-less* RNAi embryos exhibit defects spanning the L2 or L3 segment to anterior opisthosomal segments, as well as bifurcating pedipalps and reduced chelicerae (G-G’). In the same manner as Class I phenotypes, some Class II phenotypes exhibit discontinuous germ bands (H-H’ solid arrowhead). J. Phenotypic distribution of *Ptep-waist-less* RNAi and negative control embryos. Abbreviations: as, anterior spinneret; bl, book lung; ch, chelicera; L1-L4, walking legs 1-4; O1-O4, opisthosomal segments 1-4; pgz, posterior growth zone; pp, pedipalp; ps, posterior spinneret; tt, tubular trachea. Scale bars: 100 μm.

To assess the identity of the territories impacted by *Ptep-waist-less* RNAi, we assayed a segmental boundary marker (*engrailed-1*; *en1*) and a distal appendage marker (*Distal-less*; *Dll*) (Fig. 3) [33,34]. In *Ptep-waist-less* RNAi embryos, expression of *Ptep-en* was lost at the PO boundary (L4 walking leg segment and O1 opisthosomal segment in Class I embryo), with concurrent loss or diminution of *Ptep-Dll* expression (Fig. 3C, 3D). Additional posterior walking leg primordia and their corresponding *engrailed* stripes were lost in Class II embryos (Fig. 3E, 3F). Separately, we assayed *Ptep-en1* and the Hox gene *Sex combs reduced-1* (*Scr1*), which is most strongly expressed in the distal territories of the L3 and L3 limb buds (Fig. 4) [21]. *Ptep-waist-less* RNAi embryos exhibited specific and consistent reduction in *Ptep-Scr1*, concomitant with disruption of *Ptep-en* stripes in this territory (Fig. 4C, 4D). These results support the interpretation that the most pronounced effects of *Ptep-waist-less* RNAi target the segments spanning the PO boundary.

**Figure 3.**
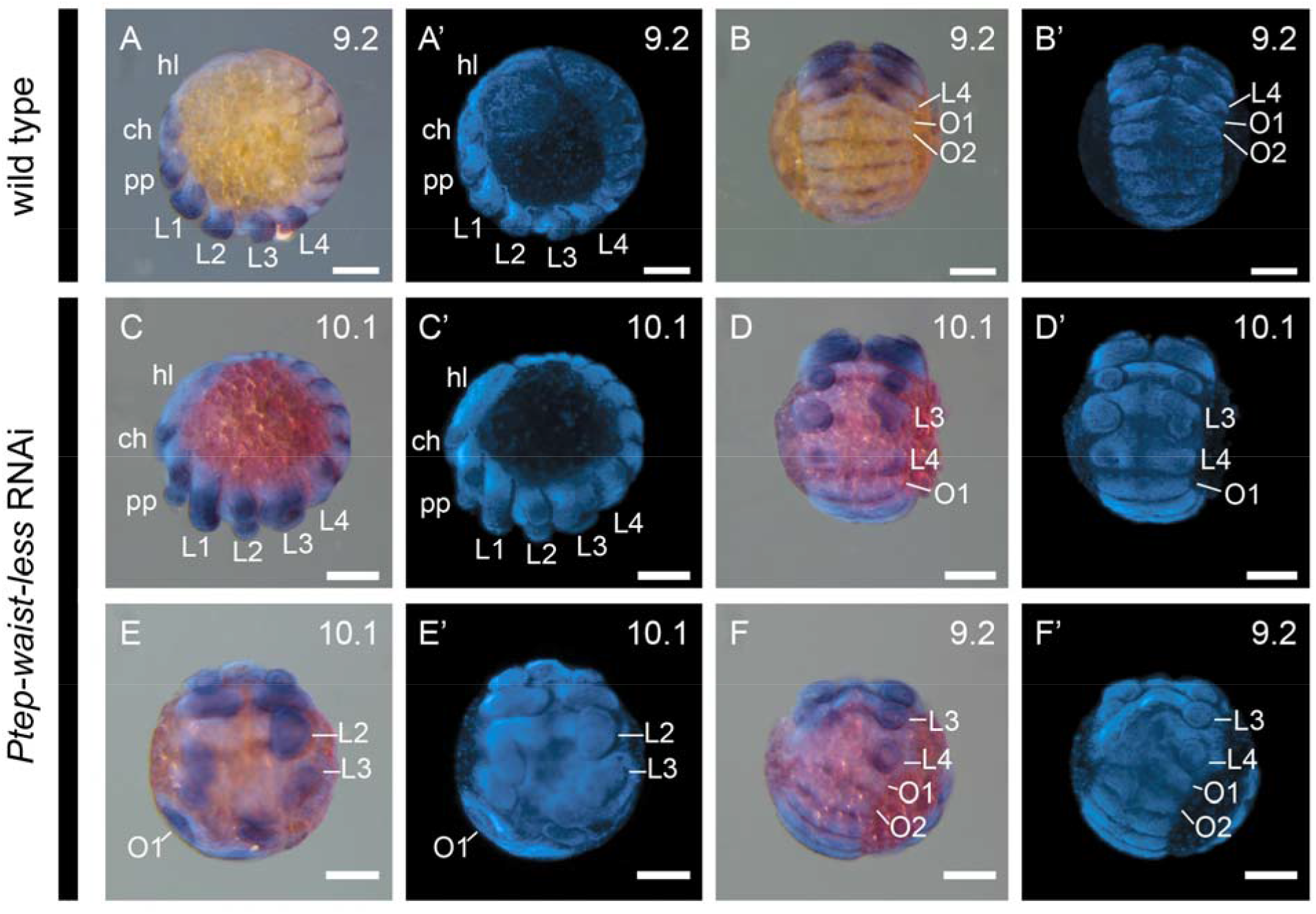
Effects of *Ptep-waist-less* RNAi affect segments spanning the prosoma-opisthosoma boundary. A-B’. Wild type embryos express the segmental marker *engrailed-1* (*en1*) in the posterior boundary of each segment; the limb-patterning gene *Distal-less* (*Dll*) is expressed in the distal part of each appendage. C-F’ *Ptep-waist-less* RNAi embryos show disruption of segments at the prosoma-opisthosoma boundary (*en1* expression lost or disrupted in L2-O1) and loss or reduction of L2-L4 appendages (*Dll* missing or disrupted). A’-F’. Hoechst counterstains of embryos in A-F. RNAi embryos have been overstained to ensure detection of riboprobes. Abbreviations: hl, head lobe. Other abbreviations as in Fig. 2. Scale bars: 100 μm.

**Figure 4.**
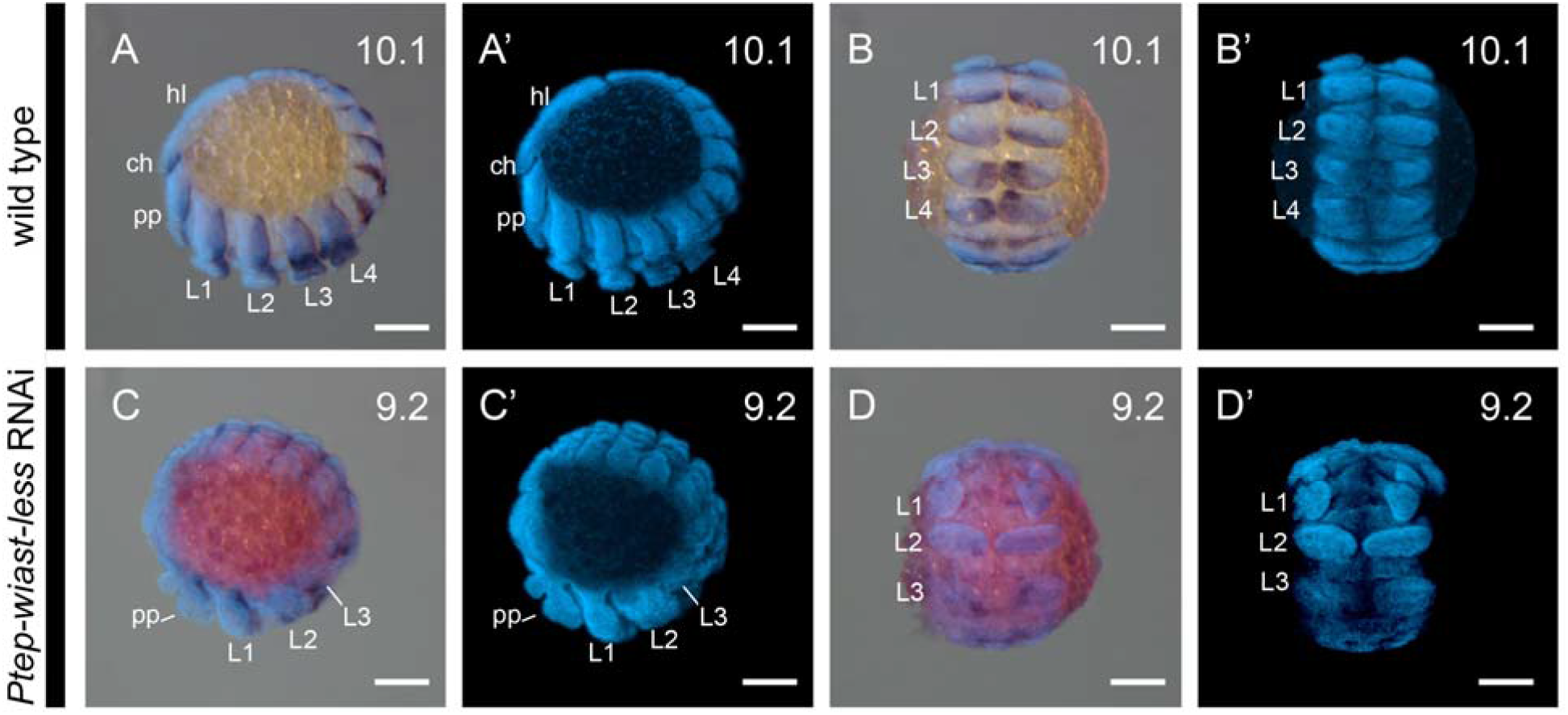
RNAi against *Ptep-waist-less* affects the posterior prosomal segments and is not associated with homeosis. A-B’. Wild type embryos express *engrailed-1* (*en1*) at the posterior boundary of each segment. The Hox gene *Sex combs reduced-1* (*Scr1*) is strongly expressed in the distal territories of L3 and L4 limbs. C-D’. *Ptep-waist-less* phenotypes show disrupted segmentation (*en1* expression lost or disrupted) and concomitant loss of the third and fourth walking legs (Class II phenotype, partial loss of L3 and L3 segmental boundary). A’-D’ Hoechst counterstains of embryos in A-D. RNAi embryos have been overstained to ensure detection of riboprobes. Abbreviations as in Fig. 2 and 3. Scale bars: 100 μm.

### *waist-less* acts through dorso-ventral patterning of the spider’s pedicel region

Disruption of the segments spanning the PO boundary in the *Ptep-waist-less* RNAi phenotype could alternatively reflect (a) a gap segmentation function localized to the boundary of the tagmata, or (b) a defect in proper dorso-ventral patterning. To distinguish between these two possibilities, we surveyed the expression of the ventral midline marker *short gastrulation* (*sog*), whose expression has been well characterized in *P. tepidariorum*, as well as other arthropods [28,35]. We reasoned that the *sog* expression domain would become discontinuous if *waist-less* bore a dorso-ventral patterning function, whereas *sog* would be unaffected by truncation of segments in a gap segmentation phenotype [7].

The lateral-most edges of the spider germ band correspond to the presumptive dorsal midline, as these two margins will fold to enclose the yolk via dorsal closure [36]. However, the canonical arthropod dorsal morphogen *decapentaplegic* is not applicable as a dorsal marker for *P. tepidariorum*, as it is not comparably expressed in the dorsal territory of developing arachnids [28,37]. In the fruit fly *Drosophila melanogaster*, the GATA family gene *pannier* is necessary for proper dorsal closure of the germ band and is also expressed in the amnioserosa (extraembryonic membrane; absent in chelicerates) [38–43]. Previous work in the fruit fly has identified antagonistic regulatory interactions between *araucan/caupolican* and *pannier* [40,44,45]. While *waist-less* is not orthologous to *Irx2/ara/caup*, our DGE analysis recovered the over-expression of one *pannier* copy in opisthosomal segments and in same gene expression cluster as *Ptep-waist-less*, across developmental stages (stages 9 and 11 of all-by-all comparisons for top 100 genes; Fig. S1). As with *Ptep-waist-less*, gene orthology was inferred using a gene tree of GATA sequences. Three spider GATA genes were identified as members of the *pannier* clade, with the highly expressed copy provisionally identified as *Ptep-pnr2* (Fig. S6).

We optimized a protocol for hybridization chain reaction for *P. tepidariorum* and assayed the expression of *Ptep-sog*, *Ptep-waist-less*, and *Ptep-pnr2* to better understand their spatial relationships. In wild type embryos, *Ptep-pnr2* is expressed in the lateral-most territory of the opisthosoma, which corresponds to the dorsal midline upon dorsal closure, as well as in a separate domain corresponding to the dorso-lateral margin of the head lobe (Fig. S7A’’’-C’’’). In the opisthosoma, *Ptep-waist-less* is expressed in a field of cells overlapping the *Ptep-pnr2*-positive territory in the dorsal margin, as well as more lateral cells (in addition to the stripes of expression in the ventral ectoderm described previously) (Fig. S7A’-C’). There is no overlap between the expression of *Ptep-sog* and the expression of either *Ptep-pnr2* or *Ptep-waist-less*, except for the ventral stripes of *Ptep-waist-less* expression (Fig. S7A’’-C’’).

We used HCR to characterize the *waist-less* loss-of-function phenotype. In *Ptep-waist-less* Class I RNAi embryos with discontinuous germ bands, the ventral midline expression of *Ptep-sog* was rendered discontinuous at the prosoma-opisthosoma boundary (Fig. 5E). In the same RNAi embryos, *Ptep-pnr2* was ectopically expressed at the narrowest point of the constricted germ band, in the territory that corresponded to the deleted ventral midline (Fig. 5F, 5G). Class I embryos of *Ptep-waist-less* RNAi without discontinuity of the AP axis (interpreted to mean a mild loss-of-function phenotype) retained *Ptep-pnr2* expression in the opisthosoma, but the expression domain of *Ptep-pnr2* was rendered irregular (Fig. S8). These results are consistent with the interpretation that *Ptep-waist-less* plays a role in dorso-ventral patterning of the segments spanning the two tagmata, rather than as a gap segmentation gene.

**Figure 5.**
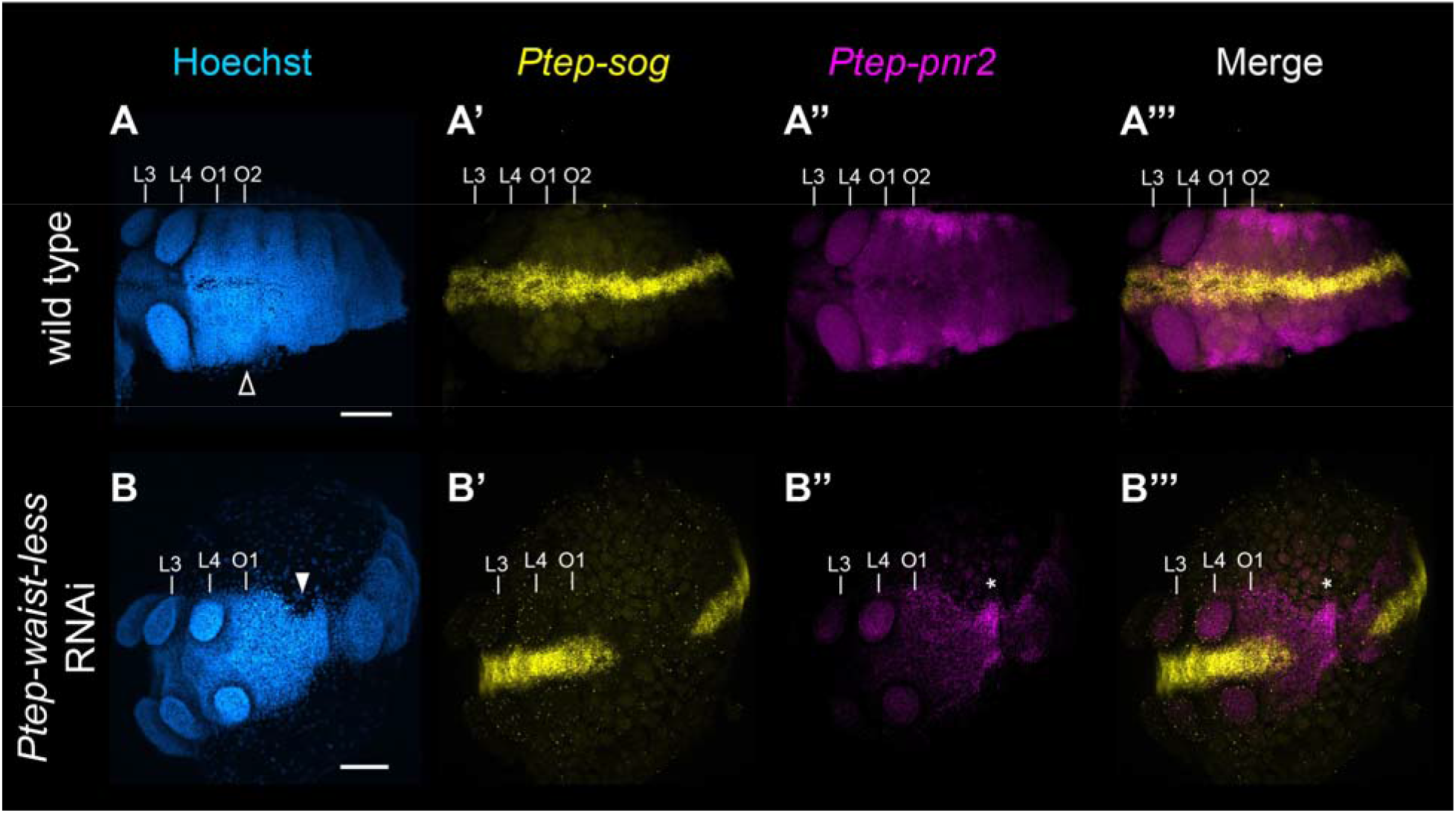
Knockdown of *Ptep-waist-less* incurs a dorso-ventral phenotype. **A.** Wild type stage 9 *P. tepidariorum* embryo with a continuous germ band (**A**, open arrowhead); continuous expression of *Ptep-sog* along the ventral midline of the antero-posterior axis (**A’**); and expression of *Ptep-pnr2* in the lateral margins of the opisthosoma (n=10/10) (**A’’**). **B.** *Ptep-waist-less* RNAi stage 9 embryo exhibit interrupted expression of the ventral marker *Ptep-sog* (**B’**) in regions affected by *Ptep-waist-less* knockdown (**B**, solid arrowhead), and concomitant expansion of *Ptep-pnr2* expression into the ventral territory (n=5/5) (**B’’**; asterisk). Abbreviations: L3-L4-walking legs 3-4; O1-O2-opisthosomal segments 1-2. Scale bars: 100 μm.

### Knockdown of *pannier2* results in ectopic dorso-lateral opisthosomal tissue in a spider

A single copy of each of *Iroquois* and *pannier* are known to occur in Onychophora (the sister group of Arthropoda) [46]. Despite the absence of *Iroquois3* homologs in pancrustaceans, we reasoned that *Iroquois3* could have retained regulatory interactions with a *pannier* homolog prior to its duplication in the common ancestor of Arthropoda. To test for gene regulatory interactions between spider *waist-less* and *pnr2*, we investigated the function of *Ptep-pnr2* using maternal RNAi. *Ptep-pnr2* RNAi embryos displayed ectopic opisthosomal tissue, resulting in a smaller proportion of extraembryonic territory in affected embryos at developmental stages associated with the beginnings of inversion and dorsal closure (n = 58/646) (Fig. 6B), as well as abnormal pouches resembling ectopic neuromeres (Fig. 6C). The opisthosoma of *Ptep-pnr2* RNAi embryos exhibited aberrant patterns of *Ptep-waist-less* expression, with loss or diminution of *Ptep-waist-less* in the ventral domains (stripes of the ventral ectoderm and ring domains in the legs), as well as expression in the dorso-lateral opisthosomal tissue (n = 4/7) (Fig. 6B’, 6C’). Intriguingly, *Ptep-pnr2* RNAi embryos also exhibited ectopic *Ptep-sog* expression in the dorsal margin of the opisthosoma (Fig. 6D’’), suggesting that *Ptep-pnr2* represses *Ptep-waist-less*. These data are consistent with the interpretation that *Ptep-pnr2* RNAi embryos also exhibit a dorso-ventral defect, wherein the dorsal midline takes on ventral identity in the absence of *Ptep-pnr2*.

**Figure 6.**
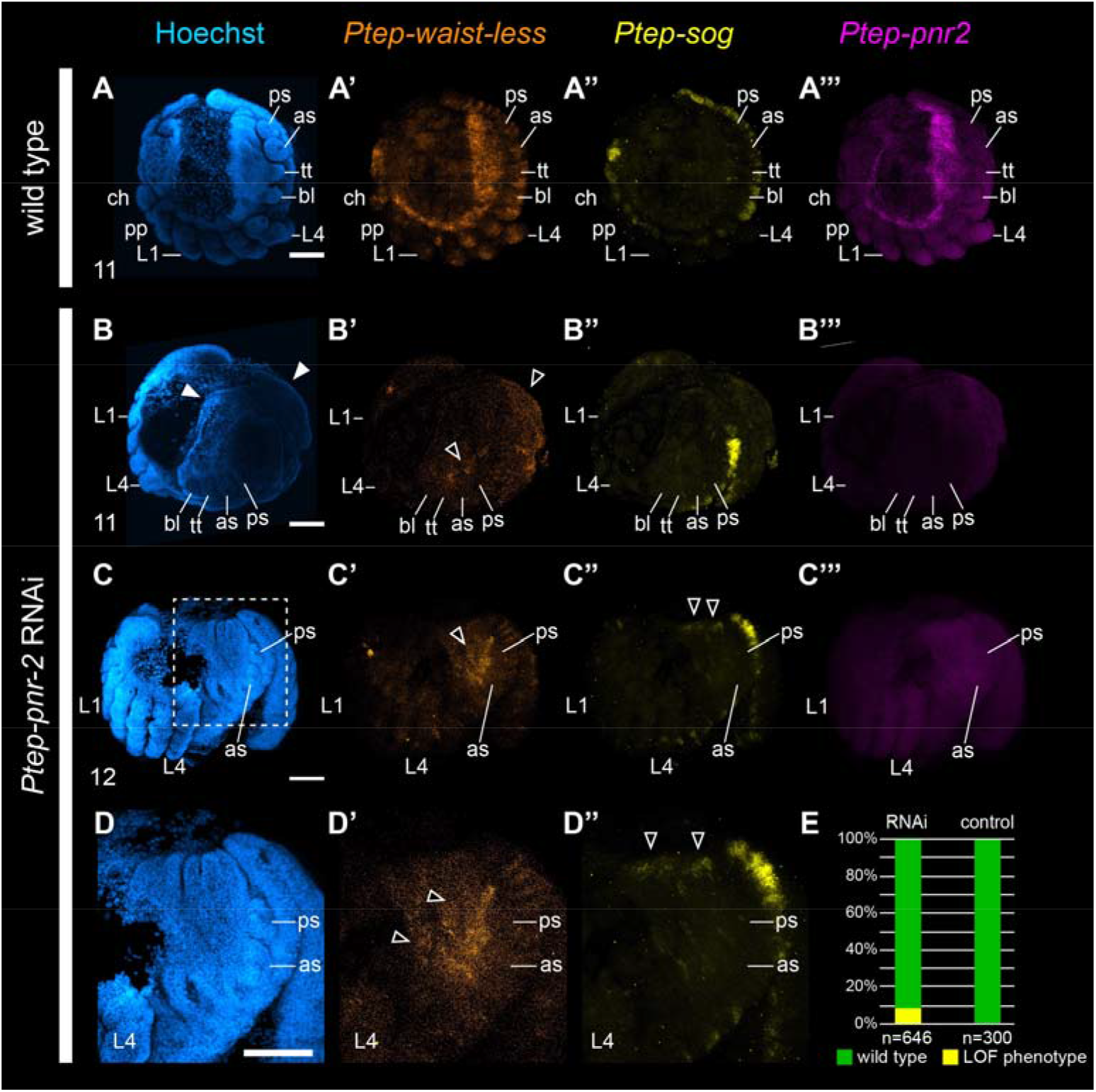
RNAi against *Ptep-pnr2* results in ectopic tissue formation in the opisthosoma and disrupts expression of *Ptep-waist-less* at the lateral boundary. A-A’’’. Wild type embryos at stage 11 (inversion) express *Ptep-waist-less* with a clear boundary of lateral expression in both body wall and migrating opisthosomal tissue (A’), expression of *Ptep-sog* is restricted to the ventral midline (A’’), and *Ptep-pnr2* is expressed in the lateral margins of the germ band with strongest expression concentrated in the opisthosoma and migrating tissues (A’’’) (n=9/9). B-C’’’. *Ptep-pnr2* loss-of function phenotypes exhibit expansion of the lateral opisthosomal territory (B, white arrowheads), with ectopic opisthosomal tissue exhibiting disrupted and non-uniform *Ptep-waist-less* expression (B’, C’, black arrowheads) along the lateral margin. (C) In later stages, ectopic expression of *Ptep-sog* was detected in the dorsal midline of the opisthosoma in *Ptep-pnr2* RNAi embryos (n=4/7) (C’’, D’’, black arrowheads). Expression of *Ptep-pnr2* was disrupted and indistinguishable from background in *Ptep-pnr2* RNAi embryos (B’’’, C’’’). D-D’’. Magnification of C-C’’ corresponding to the region outlined in C. E. Phenotypic distribution of *Ptep-pnr2* RNAi and control embryos. Abbreviations as in Figures 2 and 3. Scale bars: 100 μm.

## Discussion

### A taxon-restricted Iroquois copy is required for patterning the boundary between the tagmata of chelicerates

Comparative investigations of arthropod body plan evolution have historically focused on various aspects of morphogenesis, such as anteroposterior segmentation, neurogenesis, regionalization of body axes, and germ cell specification. Candidate gene approaches in spiders have featured prominently in such investigations, with *P. tepidariorum* serving as the leading model system representing Chelicerata. In some cases, the magnitude of the phylogenetic distance between chelicerates and insects has limited the informativeness of candidate gene suites that were established from the fruit fly literature. A separate challenge for an insect model-derived candidate gene approach is the evolution of taxon-restricted genes, as exemplified by the subdivision of *araucan* and *caupolican* (restricted to a derived group of dipterans), and by the abundance of gene duplicates resulting from WGD in groups like spiders. Here, we developed a tissue-specific transcriptomic profile of appendage-bearing segments in a large-bodied spider to circumvent these hurdles. Profiling for, and functional screening of, genes highly expressed in the spider posterior tagma resulted in the identification of *waist-less*, an Iroquois gene whose ortholog has been lost in the common ancestor of Pancrustacea. The high level of expression posterior to the PO boundary for *waist-less* and *pannier2*, as well as their roles in territory-specific dorso-ventral patterning, accorded with bioinformatic predictions of the differential gene expression analysis.

The phenotypic spectrum incurred by RNAi against *waist-less* was unexpected for an Iroquois homolog. The sparse existing data for Iroquois family genes in chelicerate taxa have encompassed only bioinformatic assays and whole mount gene expression surveys, leaving the function of this gene family unexplored in non-insect Arthropoda. Expression patterns of spider Iroquois family genes were previously interpreted to mean that these paralogs have undergone subfunctionalization after duplication in spider development, upon comparison to the expression pattern in a non-arachnopulmonate arachnid lineage (arachnids lacking a WGD event, such as harvestmen) [25]. However, that previous survey reported only one of the three Iroquois genes in the harvestman, and only four of the five in the spider *P. tepidariorum*. In addition, the expression dynamics of *waist-less* transcripts in single cell RNAseq datasets had previously been interpreted to mean that this spider gene played a role in antero-posterior segmentation and/or neural development [32].

By contrast, in the fruit fly *D. melanogaster*, the two homologs of *waist-less* (cyclorraphan fly-specific duplicates *araucan* and *caupolican*) are broadly pleiotropic, acting in multiple contexts that span dorso-lateral patterning of the body wall; heart, eye, and muscle development; extra-embryonic tissue specification; and neurogenesis and sensory structure development [44,47–52]. In many of these functional contexts, Irx copies of the *ara/caup* group have been demonstrated to serve redundant functions, including dorso-ventral patterning [48]. Nevertheless, a role in regionalized tissue maintenance (i.e., the discontinuous germ bands in loss-of-function phenotypes that were found in this study) was not known for any arthropod Iroquois homolog. Taken together with the present results, available data points for arthropod Iroquois homologs suggest that the dorso-ventral patterning function of *waist-less* in Chelicerata is partly conserved, whereas the regionalized tissue maintenance function at the boundary of the two spider tagmata reflects taxon-specific neofunctionalization (Fig. 7). Moreover, ectopic expression of *Ptep-pnr2* in the PO boundary upon *Ptep-waist-less* knockdown, as well as ectopic *Ptep-waist-less* and *Ptep-sog* expression upon *Ptep-pnr2* knockdown, is consistent with conservation of antagonistic regulatory interactions between *Iroquois* and *pannier* homologs in insects and spiders (and by extension, across Arthropoda, given the phylogenetic position of hexapods and chelicerates).

**Figure 7.**
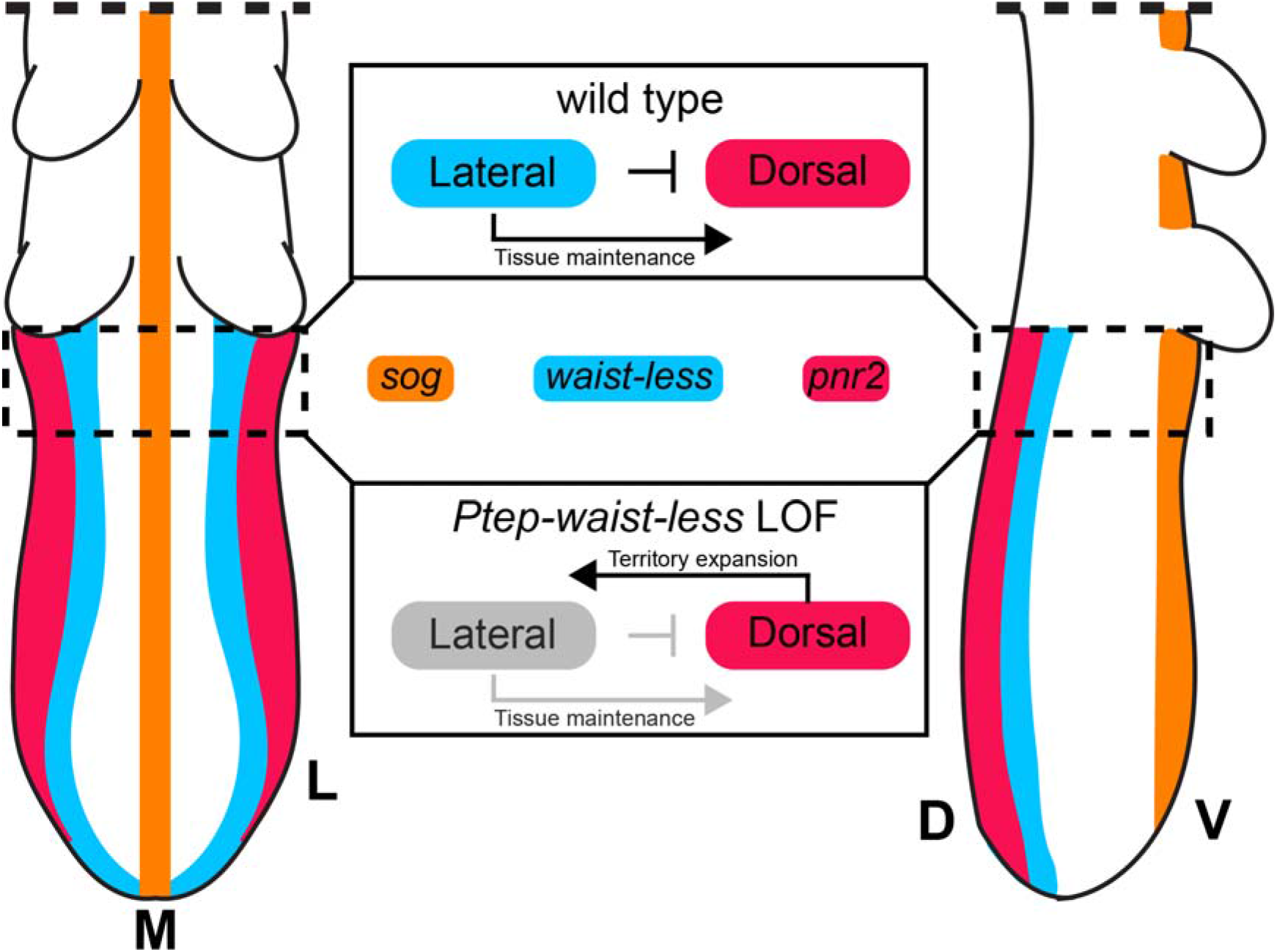
Simplified model of interactions between *Ptep-waist-less* and *Ptep-pnr2* in the anterior opisthosoma. Abbreviations: D, dorsal; L, lateral; M, medial; V, ventral.

### Association of *Iroquois3* with the prosoma-opisthosoma boundary predates the WGD of arachnopulmonates

Investigating the genetic architecture of the spider pedicel highlights the challenges of the candidate gene approach in emerging model systems; classic arthropod models (holometabolous insects) lack the pedicel, as well as other taxon-specific structures of interest. This has precluded developmental genetic investigations of iconic arachnid organs like spider venom glands, silk spigots, and fangs. The RNAseq datasets established herein for limb bud-bearing territories of embryonic spiders, spanning developmental stages most salient to development of posterior appendages, are anticipated to guide future investigations of book lung and (spider) spinneret development, two arachnid organs that have prompted prolonged debates over evolutionary origins and serial homology [53–55].

One potential limitation of spiders as model systems for study of chelicerate developmental biology is the incidence of whole genome duplication in the common ancestor of Arachnopulmonata, as the ancestral condition for Chelicerata is unambiguously an unduplicated genome [20,56,57]. In the context of the present study, the function of *waist-less* is of interest from the perspective of body plan evolution, but *waist-less* is itself a duplicated copy; groups like mites, ticks, and harvestmen possess only one homolog of *Iroquois3*. It is therefore not clear whether the function of *waist-less* reflects a dynamic conserved across Chelicerata, or whether it represents an arachnopulmonate-specific novelty. To discern between these possibilities, we examined the expression of *Iroquois2* and *Iroquois3* single-copy homologs in the harvestman *Phalangium opilio*. In stages corresponding to opisthosomal segment addition, *Iroquois2* was expressed in the dorso-lateral body wall throughout the germ band, whereas *Iroquois3* exhibited a more complex pattern with expression in the head, the distal territory of the appendages (strongest in L2 leg), the ventral ectoderm, and in the dorso-lateral body wall territory spanning the L4 segment to the posterior terminus (Fig. S9). Expression patterns are therefore closely comparable between harvestman *Iroquois3*, *P. tepidariorum waist-less*, and the bioinformatic predictions of the DGE datasets for the *waist-less* ortholog of the tarantula *A. hentzi*. By contrast, we found that other spider Iroquois homologs were not comparably expressed, either with respect to in situ hybridization assays (Fig. S10) or expression dynamics inferred from RNA-Seq (Fig. S11, S12, S13). These data suggest that an association between *Iroquois3* and the PO boundary predates the divergence of arachnopulmonates, and that an association with the PO boundary is retained only by the spider *waist-less* (*Iroquois3-2*) ortholog (but not *Iroquois3-1*).

The availability of these data for the spider and the harvestman may facilitate broader tests of how arachnopulmonate copies divergence as a function of phylogenetic distance. Specifically, the recently established availability of developmental genetic resources for non-spider arachnopulmonates like scorpions [22] and whip spiders [58] may enable investigation of whether Iroquois duplicates faithfully retain expression domains as a function of orthology, or whether they exhibit developmental system drift. The sum of these comparative datasets, extended to *pannier* homologs, may also aid in pinpointing whether tagma-specific regionalization of one gene preceded the regionalization of its regulatory partner. As a corollary, investigating the activity of *Iroquois3* and *pannier* in chelicerate taxa that have undergone reduction of the opisthosoma may aid in testing the inference of opisthosoma-specific activity of these two genes. Specifically, comparative data from Pycnogonida (sea spiders), which retain only a rudiment of the opisthosoma, may aid in understanding how tagmata evolve [59].

## Methods

### Field collection, sequencing, and differential gene expression analyses of tarantula embryos

Field collection protocols for *Aphonopelma hentzi* (Araneae: Theraphosidae) embryos and laboratory protocols for care were previously described by Setton et. al [27].Embryos used to generate the developmental transcriptomic resources were stored in Trizol Tri-reagent (Ambion Life Technologies, Waltham, MA, USA) prior to RNA extraction, following manufacturer’s protocols. Library preparation and stranded mRNA sequencing were performed at the University of Wisconsin-Madison Biotechnology Center on an Illumina HiSeq 2500 platform with 2×100 PE reads. The transcriptome spans developmental stages 9.1 to 13 (following Mittmann and Wolff, 2012; Setton et al., 2019) to juveniles (1^st^-2^nd^ instar post-hatching). This resource is available under accession numbers NCBI SRR13605914 and SRR13605915 [20].

Three sets of biological replicates for each tarantula appendage type were dissected at three different time points during embryogenesis (stages 9, 10, and 11 after Setton et al. 2019). Each experimental sample contained appendages from multiple individuals from the same clutch (n = 3 to 7 samples per appendage type). Total RNA was extracted from whole embryos using TRIzol Tri Reagent (Ambion Life Technologies, Waltham, MA, USA), following the manufacturer’s protocol. Libraries were prepared for sequencing using standard protocols for the Illumina NovaSeq 6000 platform with a 2×150 PE sequencing strategy (stages 9 and 10) or the Illumina HiSeq 2500 platform with 1×100 SE sequencing strategy (stage 11). Multiplexing was designed to recover an expected 15M reads per library for stages 9 and 10, and 12M reads per library for stage 11. Adaptor removal and quality trimming was conducted using Trimmomatic v 0.35 (Bolger et al. 2014) prior to analysis.

Reads were mapped to the *A. hentzi* developmental transcriptome using the density of reads mapped to the transcriptome as a proxy for transcript abundance, as implemented by salmon v. 0.9.1 under default parameters [60] . Differential gene expression analysis was performed using DESeq2 v. 1.14.1 [61])

### Gene tree analysis and orthology inference

BLAST and BLASTp searches were used to determine the identities of transcripts identified in DGE datasets. Orthology of *A. hentzi Iroquois* homologs was determined using previously published *P. tepidariorum Irx* sequences as queries [26,32] for tBLASTn searches, and hits with e-values < 10^-5^ were retained. All putative orthologs were verified using reciprocal BLAST searches. Multiple sequence alignment was conducted *de novo* with MAFFT v.7 with default parameters [62]. Identification of *A. hentzi pannier* homologs was determined using previously published *D. melanogaster pannier* and *grain* sequences as queries for tBLASTn searches, and hits with e-values < 10^-5^ were retained. Vertebrate sequences included the GATA1-3 (*grain*) group and GATA4-6 (*pnr*) group were taken from a previously published study [63]. All putative orthologs were verified via reciprocal BLAST searches, as with Iroquois orthologs. Multiple sequence alignment was conducted *de novo* with CLUSTAL Omega [64]

Phylogenetic reconstruction of Iroquois amino acid alignments consisted of maximum likelihood analysis with IQ-TREE, with automated model selection (-m MFP; chosen model: LG+G4) and 1000 ultrafast bootstrap resampling replicates [65]. Chelicerate sequences were pulled from previously published genome or transcriptome assemblies available on GenBank and insect sequences were added from a previous work on the Iroquois gene family [31]. Phylogenetic reconstruction of pannier amino acid alignments consisted of maximum likelihood analysis with IQ-TREE, with automated model selection (-m MFP; chosen model: DCMut+F+R5) and 1000 ultrafast bootstrap resampling replicates. All alignments, annotated tree files and log files are available as supplementary material.

### Cloning of orthologs and probe synthesis

Fragments of *Ptep*-*irx4* were amplified using standard PCR protocols and cloned using the TOPO® TA Cloning® Kit using One Shot® Top10 chemically competent *Escherichia coli* (Invitrogen, Carlsbad, CA, USA) following the manufacturer’s protocol, and their PCR product identities were verified via sequencing with M13 universal primers. All gene-specific primer sequences are provided in SI Appendix, Table S3. Upon completion of probe synthesis, the presence of the target sequence was checked using gel electrophoresis.

### House spider embryo collection, fixation, in situ hybridization, and imaging

Animals were maintained, and embryos fixed and assayed for gene expression, following established or minimally modified protocols for colorimetric *in situ* hybridization, as detailed previously [27,30,37]. PCRs for generating riboprobe templates, synthesis of DIG-labeled probes, and preservation of embryos all followed recently detailed procedures (Setton and Sharma 2018, 2021). Whole mount images were taken using a Nikon SMZ25 fluorescence stereomicroscope mounted with a DS-Fi2 digital color camera driven by Nikon Elements software.

For hybridization chain reaction (HCR) gene expression assays, probes were designed separately for each gene using an open-source probe design platform [66] with standard parameters and 20 probe pairs per gene returned; *Ptep-*sog was designed with the delay parameter set to 50 and the number of probe pairs was set to 30. Expression for *Ptep-Irx2-2* could not be surveyed using HCR due to the short sequence length (747bp). For instances where genes with regions of high sequence similarity were multiplexed into a single probe, the regions of highly similar sequence were identified using sequence alignments and then masked in Aliview v.1.28 prior to probe design [67]. All probes were designed to span a maximal amount of ORF and minimal UTR (SI Appendix, Table S4).

For HCR, embryos of *P. tepidariorum* were fixed by dechorionation in 50% bleach solution (Clorox brand) and fixed in a 3.2% paraformaldehyde (PFA) solution in PBS for 35 min. Vitelline membranes were manually removed using fine forceps during the fixation in PFA solution. Embryos were washed in PBS-Tween20 several times and serially dehydrated into 100% ethanol for storage at −20° C. The procedures for HCR, and all solutions therein, constitute minor modifications of a recently published protocol [68]. For spiders, we lowered the amount of probe hybridization solution to 148 μL and added in probe stocks at 2×-4× suggested concentration (1.6 μL to 3.2 μL probe per gene). Confocal imaging was conducted on a Zeiss LSM710 confocal microscope driven by Zen software.

### Double-stranded RNA synthesis and maternal RNA interference

Double-stranded RNA (dsRNA) was synthesized following the manufacturer’s protocol using a MEGAscript® T7 kit (Ambion/Life Technologies, Grand Island, NY, USA) from amplified PCR product. dsRNA quality was checked, and concentration adjusted to 2.5 μg/μl. For *Ptep-waist-less*, RNAi was performed with 20 μg of dsRNA of a 978 bp fragment, delivered over eight days to 32 virgin females, with 22 surviving to laying the second cocoon. Of these 22 females injected with *Ptep-waist-less* dsRNA, 13 produced at least one cocoon of embryos with phenotypes; embryos were collected from cocoons 2-5 as previously described (Setton and Sharma 2018, 2021). Negative controls with injected with an equal volume of deionized water, following established protocols in spiders [17]; 12 females were injected thus, with seven laying beyond cocoon 2. To rule out off-target effects of RNAi, we performed gene silencing using two non-overlapping fragments of *Ptep-waist-less* (473 bp and 405 bp fragments), delivered at the same concentration, and assessed the resulting phenotypic spectra to confirm identical phenotypes. Counts of phenotypes were obtained from a randomly selected group of embryos spanning clutches 3-5 of multiple females, for both RNAi and negative control experiments.

For *Ptep-pnr2* RNAi, dsRNA was injected at a concentration of 4 μg/μl for a total of 32 μg administered over 8 days, following optimization of dsRNA delivery for this gene. Three virgin females were injected comparably to the *Ptep-waist-less* experiments, with another two females injected as negative controls. Three out of four females laid egg sacs and embryos were collected from cocoons 2-5 as previously described (Setton and Sharma 2018, 2021). Counts of phenotypes were obtained from a randomly selected group of embryos spanning clutches 3-5 of multiple females, for both RNAi and negative controls experiments.

## Acknowledgements

Confocal microscopy was performed at the Newcomb Imaging Center, Department of Botany, University of Wisconsin–Madison. Guilherme Gainett assisted with the pipeline for differential gene expression analyses. Fieldwork in Colorado in 2018 was supported by an American Arachnological Society Student Research award to EVWS. EVWS was additionally supported by a National Science Foundation Graduate Research Fellowship in Biology (DGE-1747503). This material is based on work supported by the National Science Foundation under grants IOS-1552610 and IOS-2016141.

## References

1. Chipman A. Parallel evolution of segmentation by coLJoption of ancestral gene regulatory networks. BioEssays. 2010;32: 60–70.

2. Davidson EH, Erwin DH. Gene regulatory networks and the evolution of animal body plans. Science. 2006;311: 796–800. doi:10.1126/science.1113832

3. Murugesan SN, Connahs H, Matsuoka Y, Das Gupta M, Tiong GJL, Huq M, et al. Butterfly eyespots evolved via cooption of an ancestral gene-regulatory network that also patterns antennae, legs, and wings. Proc Natl Acad Sci. 2022;119: e2108661119. doi:10.1073/pnas.2108661119

4. Stahi R, Chipman AD. Blastoderm segmentation in Oncopeltus fasciatus and the evolution of insect segmentation mechanisms. Proc Biol Sci. 2016;283: 20161745–9. doi:10.1098/rspb.2016.1745

5. Chipman AD, Arthur W, Akam M. Early development and segment formation in the centipede, Strigamia maritima (Geophilomorpha). Evol Dev. 2004;6: 78–89.

6. McGregor AP, Pechmann M, Schwager EE, Feitosa NM, Kruck S, Aranda M, et al. Wnt8 is required for growth-zone establishment and development of opisthosomal segments in a spider. Curr Biol. 2008;18: 1619–1623. doi:10.1016/j.cub.2008.08.045

7. Pechmann M, Mcgregor AP, Schwager EE, Feitosa NM, Damen WGM. Dynamic gene expression is required for anterior regionalization in a spider. Proc Natl Acad Sci. 2009;106: 1468–1472. doi:10.1073/pnas.0811150106

8. Pechmann M, Prpic N-M. Appendage patterning in the South American bird spider Acanthoscurria geniculata (Araneae: Mygalomorphae). Dev Genes Evol. 2009;219: 189– 198. doi:10.1007/s00427-009-0279-7

9. Prpic N-M, Damen WGM. Notch-mediated segmentation of the appendages is a molecular phylotypic trait of the arthropods. Dev Biol. 2009;326: 262–271. doi:10.1016/j.ydbio.2008.10.049

10. Schoppmeier M, Damen WG. Double-stranded RNA interference in the spider Cupiennius saleiLJ: the role of Distal-less is evolutionarily conserved in arthropod appendage formation. Dev Genes Evol. 2001;211: 76–82. doi:10.1007/s004270000121

11. Iwasaki-Yokozawa S, Nanjo R, Akiyama-Oda Y, Oda H. Lineage-specific, fast-evolving GATA-like gene regulates zygotic gene activation to promote endoderm specification and pattern formation in the Theridiidae spider. BMC Biol. 2022;20: 223. doi:10.1186/s12915-022-01421-0

12. Posnien N, Zeng V, Schwager EE, Pechmann M, Hilbrant M, Keefe JD, et al. A comprehensive reference transcriptome resource for the common house spider Parasteatoda tepidariorum. PLoS ONE. 2014;9: e104885. doi:10.1371/journal.pone.0104885

13. Schomburg C, Turetzek N, Prpic N-M. Candidate gene screen for potential interaction partners and regulatory targets of the Hox gene labial in the spider Parasteatoda tepidariorum. 2020; 1–16. doi:10.1007/s00427-020-00656-7

14. Damen WGM. Evolutionary conservation and divergence of the segmentation process in arthropods. Dev Dyn. 2007;236: 1379–1391. doi:10.1002/(issn)1097-0177

15. Fusco G, Minelli A. Arthropod Segmentation and Tagmosis. Arthropod Biology and Evolution: Molecules, Development, Morphology. 2013. pp. 197–222.

16. Paese CLB, Schoenauer A, Leite DJ, Russell S, McGregor AP. A SoxB gene acts as an anterior gap gene and regulates posterior segment addition in a spider. eLife. 2018;7: e37567. doi:10.7554/eLife.37567

17. Pechmann M, Benton MA, Kenny NJ, Posnien N, Roth S. A novel role for Ets4 in axis specification and cell migration in the spider Parasteatoda tepidariorum. Elife. 2017;6: 1735. doi:10.7554/elife.27590

18. Turetzek N, Khadjeh S, Schomburg C, Prpic N-M. Rapid diversification of homothorax expression patterns after gene duplication in spiders. 2017; 1–12. doi:10.1186/s12862-017-1013-0

19. Ballesteros JA, Santibáñez-López CE, Baker CM, Benavides LR, Cunha TJ, Gainett G, et al. Comprehensive Species Sampling and Sophisticated Algorithmic Approaches Refute the Monophyly of Arachnida. Teeling E, editor. Mol Biol Evol. 2022;39: msac021. doi:10.1093/molbev/msac021

20. Ontano AZ, Gainett G, Aharon S, Ballesteros JA, Benavides LR, Corbett KF, et al. Taxonomic Sampling and Rare Genomic Changes Overcome Long-Branch Attraction in the Phylogenetic Placement of Pseudoscorpions. Mol Biol Evol. 2021;38: 2446–2467. doi:10.1093/molbev/msab038

21. Schwager EE, Sharma PP, Clarke T, Leite DJ, Wierschin T, Pechmann M, et al. The house spider genome reveals an ancient whole-genome duplication during arachnid evolution. BMC Biol. 2017;15: 62. doi:10.1186/s12915-017-0399-x

22. Sharma PP, Schwager EE, Extavour CG, Wheeler WC. Hox gene duplications correlate with posterior heteronomy in scorpions. Proc R Soc B Biol Sci. 2014;281: 20140661. doi:10.1016/j.cub.2009.06.061

23. Benton MA, Pechmann M, Frey N, Stappert D, Conrads KH, Chen Y-T, et al. Toll Genes Have an Ancestral Role in Axis Elongation. Curr Biol. 2016;26: 1609–1615. doi:10.1016/j.cub.2016.04.055

24. Turetzek N, Pechmann M, Schomburg C, Schneider J, Prpic N-M. Neofunctionalization of a Duplicate dachshundGene Underlies the Evolution of a Novel Leg Segment in Arachnids. Mol Biol Evol. 2015;33: 109–121. doi:10.1093/molbev/msv200

25. Leite DJ, Baudouin-Gonzalez L, Iwasaki-Yokozawa S, Lozano-Fernandez J, Turetzek N, Akiyama-Oda Y, et al. Homeobox Gene Duplication and Divergence in Arachnids. O’Connell MJ, editor. Mol Biol Evol. 2018;35: 2240–2253. doi:10.1093/molbev/msy125

26. Aase-Remedios ME, Janssen R, Leite DJ, Sumner-Rooney L, McGregor AP. Evolution of the spider homeobox gene repertoire by tandem and whole genome duplication. Evolutionary Biology; 2023 May. doi:10.1101/2023.05.26.542232

27. Setton EVW, Hendrixson BE, Sharma PP. Embryogenesis in a Colorado population of Aphonopelma hentzi (Girard, 1852) (Araneae: Mygalomorphae: Theraphosidae): establishing a promising system for the study of mygalomorph development. J Arachnol. 2019;47: 209–216.

28. Akiyama-Oda Y. Axis specification in the spider embryo: dpp is required for radial-to-axial symmetry transformation and sog for ventral patterning. Development. 2006;133: 2347– 2357. doi:10.1242/dev.02400

29. Setton EVW, Sharma PP. A conserved role for arrow in posterior axis patterning across Arthropoda. Dev Biol. 2021;475: 91–105. doi:10.1016/j.ydbio.2021.02.006

30. Setton EVW, Sharma PP. Cooption of an appendage-patterning gene cassette in the head segmentation of arachnids. Proc Natl Acad Sci. 2018;115: E3491–E3500. doi:10.1073/pnas.1720193115

31. Kerner P, Ikmi A, Coen D, Vervoort M. Evolutionary history of the iroquois/Irx genes in metazoans. BMC Evol Biol. 2009;9: 74. doi:10.1186/1471-2148-9-74

32. Leite DJ, Schönauer A, Blakeley G, Harper A, Garcia-Castro H, Baudouin-Gonzalez L, et al. An atlas of spider development at single-cell resolution provides new insights into arthropod embryogenesis. bioRxiv. 2022. doi:10.1101/2022.06.09.495456

33. Damen WGM. Parasegmental organization of the spider embryo implies that the parasegment is an evolutionary conserved entity in arthropod embryogenesis. Development. 2002;129: 1239–1250.

34. Pechmann M, Khadjeh S, Turetzek N, Mcgregor AP, Damen WGM, Prpic N-M. Novel Function of Distal-less as a Gap Gene during Spider Segmentation. PLoS Genet. 2011;7: e1002342. doi:10.1371/journal.pgen.1002342.g004

35. Oda H, Iwasaki-Yokozawa S, Usui T, Akiyama-Oda Y. Experimental duplication of bilaterian body axes in spider embryos: Holm’s organizer and self-regulation of embryonic fields. Dev Genes Evol. 2020;230: 49–63. doi:10.1007/s00427-019-00631-x

36. Mittmann B, Wolff C. Embryonic development and staging of the cobweb spider Parasteatoda tepidariorum C. L. Koch, 1841 (syn.: Achaearanea tepidariorum; Araneomorphae; Theridiidae). Dev Genes Evol. 2012;222: 189–216. doi:10.1007/s00427-012-0401-0

37. Akiyama-Oda Y, Oda H. Early patterning of the spider embryo: a cluster of mesenchymal cells at the cumulus produces Dpp signals received by germ disc epithelial cells. Development. 2003;130: 1735–1747. doi:10.1242/dev.00390

38. Ashe HL, Mannervik M, Levine M. Dpp signaling thresholds in the dorsal ectoderm of the *Drosophila* embryo. Development. 2000;127: 3305–3312. doi:10.1242/dev.127.15.3305

39. Heitzler P, Haenlin M, Ramain P, Calleja M, Simpson P. A Genetic Analysis of *pannier*, a Gene Necessary for Viability of Dorsal Tissues and Bristle Positioning in Drosophila. Genetics. 1996;143: 1271–1286. doi:10.1093/genetics/143.3.1271

40. Herranz H, Morata G. The functions of *pannier* during *Drosophila* embryogenesis. Development. 2001;128: 4837–4846. doi:10.1242/dev.128.23.4837

41. Jürgens G, Wieschaus E, Nüsslein-Volhard C, Kluding H. Mutations affecting the pattern of the larval cuticle inDrosophila melanogaster. Wilhelm Rouxs Arch Dev Biol. 1984;193: 283–295. doi:10.1007/BF00848157

42. Sharma R, Beermann A, Schröder R. The dynamic expression of extraembryonic marker genes in the beetle Tribolium castaneum reveals the complexity of serosa and amnion formation in a short germ insect. Gene Expr Patterns. 2013;13: 362–371. doi:10.1016/j.gep.2013.07.002

43. Winick J, Abel T, Leonard MW, M. Michelson A, Chardon-Loriaux I, Holmgren RA, et al. A GATA family transcription factor is expressed along the embryonic dorsoventral axis in *Drosophila melanogaster*. Development. 1993;119: 1055–1065. doi:10.1242/dev.119.4.1055

44. Calleja M, Herranz H, Estella C, Casal J, Lawrence P, Simpson P, et al. Generation of medial and lateral dorsal body domains by the *pannier* gene of *Drosophila*. Development. 2000;127: 3971–3980. doi:10.1242/dev.127.18.3971

45. Letizia A, Barrio R, Campuzano S. Antagonistic and cooperative actions of the EGFR and Dpp pathways on the *iroquois* genes regulate *Drosophila* mesothorax specification and patterning. Development. 2007;134: 1337–1346. doi:10.1242/dev.02823

46. Treffkorn S, Mayer G, Janssen R. Review of extra-embryonic tissues in the closest arthropod relatives, onychophorans and tardigrades. Philos Trans R Soc B Biol Sci. 2022;377: 20210270. doi:10.1098/rstb.2021.0270

47. Carrasco-Rando M, Tutor AS, Prieto-Sánchez S, González-Pérez E, Barrios N, Letizia A, et al. Drosophila Araucan and Caupolican Integrate Intrinsic and Signalling Inputs for the Acquisition by Muscle Progenitors of the Lateral Transverse Fate. Perrimon N, editor. PLoS Genet. 2011;7: e1002186. doi:10.1371/journal.pgen.1002186

48. Cavodeassi F, Modolell J, Gómez-Skarmeta JL. The Iroquois family of genes: from body building to neural patterning. Development. 2001;128: 2847–2855. doi:10.1242/dev.128.15.2847

49. del Corral RD, Aroca P, Gomez-Skarmeta JL, Cavodeassi F, Modolell J. The Iroquois homeodomain proteins are required to specify body wall identity in Drosophila. Genes Dev. 1999;13: 1754–1761. doi:10.1101/gad.13.13.1754

50. Ikmi A, Netter S, Coen D. Prepatterning the Drosophila notum: The three genes of the iroquois complex play intrinsically distinct roles. Dev Biol. 2008;317: 634–648. doi:10.1016/j.ydbio.2007.12.034

51. Leyns L, Gómez-Skarmeta J-L, Dambly-Chaudière C. iroquois: a prepattern gene that controls the formation of bristles on the thorax ofDrosophila. Mech Dev. 1996;59: 63–72. doi:10.1016/0925-4773(96)00577-1

52. Mirzoyan Z, Pandur P. The Iroquois Complex Is Required in the Dorsal Mesoderm to Ensure Normal Heart Development in Drosophila. Singh A, editor. PLoS ONE. 2013;8: e76498. doi:10.1371/journal.pone.0076498

53. Damen W, Saridaki T, Averof M. Diverse Adaptations of an Ancestral Gill A Common Evolutionary Origin for Wings, Breathing Organs, and Spinnerets. Curr Biol. 2002;12: 1711–1716.

54. Sharma PP. Chelicerates and the Conquest of Land: A View of Arachnid Origins Through an Evo-Devo Spyglass. Integr Comp Biol. 2017;57: 510–522. doi:10.1093/icb/icx078

55. Shultz JW. The origin of the spinning apparatus in spiders. Biol Rev. 1987;62: 89–113. doi:10.1111/j.1469-185X.1987.tb01263.x

56. Gainett G, González VL, Ballesteros JA, Setton EVW, Baker CM, Barolo Gargiulo L, et al. The genome of a daddy-long-legs (Opiliones) illuminates the evolution of arachnid appendages. Proc R Soc B Biol Sci. 2021;288: 20211168. doi:10.1098/rspb.2021.1168

57. Schwager EE, Schoppmeier M, Pechmann M, Damen WG. Duplicated Hox genes in the spider Cupiennius salei. Front Zool. 2007;4: 10. doi:10.1186/1742-9994-4-10

58. Gainett G, Sharma PP. Genomic resources and toolkits for developmental study of whip spiders (Amblypygi) provide insights into arachnid genome evolution and antenniform leg patterning. EvoDevo. 2020;11: 18. doi:10.1186/s13227-020-00163-w

59. Arango CP. Morphological phylogenetics of the sea spiders (Arthropoda: Pycnogonida). Org Divers Evol. 2002;2: 107–125.

60. Patro R, Duggal G, Love MI, Irizarry RA, Kingsford C. Salmon provides fast and bias-aware quantification of transcript expression. Nat Methods. 2017;14: 417–419. doi:10.1038/nmeth.4197

61. Love MI, Huber W, Anders S. Moderated estimation of fold change and dispersion for RNA-seq data with DESeq2. Genome Biol. 2014;15: 31–21. doi:10.1186/s13059-014-0550-8

62. Katoh K, Standley DM. MAFFT multiple sequence alignment software version 7: improvements in performance and usability. Mol Biol Evol. 2013;30: 772–780. doi:10.1093/molbev/mst010

63. Gillis WJ, Bowerman B, Schneider SQ. Ectoderm- and endomesoderm-specific GATA transcription factors in the marine annelid Platynereis dumerilli: Polychaete GATA factors. Evol Dev. 2007;9: 39–50. doi:10.1111/j.1525-142X.2006.00136.x

64. Sievers F, Wilm A, Dineen D, Gibson TJ, Karplus K, Li W, et al. Fast, scalable generation of highLJquality protein multiple sequence alignments using Clustal Omega. Mol Syst Biol. 2011;7: 539. doi:10.1038/msb.2011.75

65. Nguyen L-T, Schmidt HA, Haeseler A von, Minh BQ. IQ-TREE: A Fast and Effective Stochastic Algorithm for Estimating Maximum-Likelihood Phylogenies. Mol Biol Evol. 2014;32: 268–274. doi:10.1093/molbev/msu300

66. Kuehn E, Clausen DS, Null RW, Metzger BM, Willis AD, Özpolat BD. Segment number threshold determines juvenile onset of germline cluster expansion in Platynereis dumerilii. J Exp Zoolog B Mol Dev Evol. 2022;338: 225–240. doi:10.1002/jez.b.23100

67. Larsson A. AliView: a fast and lightweight alignment viewer and editor for large datasets. Bioinformatics. 2014;30: 3276–3278. doi:10.1093/bioinformatics/btu531

68. Bruce HS, Jerz G, Kelly SR, McCarthy J, Pomerantz A, Senevirathne G, et al. Hybridization Chain Reaction (HCR) In Situ Protocol. protocols.io; 2021. Available: https://dx.doi.org/10.17504/protocols.io.bunznvf6

