## Supporting Information for "A taxon-restricted duplicate of *Iroquois3* is required for patterning the spider waist"

Author contributions: EVWS and PPS conceptualized the project. EVWS and PPS designed experiments. Bioinformatic work was conducted by EVWS, JAB, and PPS. Colorimetric gene expression assays and imaging were performed by POB, BCK, and EVWS. Fluorescent gene expression and confocal microscopy was conducted by EVWS. EVWS and PPS wrote the manuscript. Funding was acquired by PPS. All authors approved the manuscript.

Keywords: body plan | gene duplication | homeodomain | Iroquois | tagmosis

### Supporting Information

**Table S1.** List of genes screened via RNAi in *Parasteatoda tepidariorum*.

| **Gene** | **Number injected** | **Number survived to laying (C1)** | **Number survived to laying (C2)** | **Embryos?** | **Phenotype?** |
| --- | --- | --- | --- | --- | --- |
| *waist-less* | 32 | 23 | 22 | Yes | Yes |
| *pnr2* | 4 | 3 | 3 | Yes | Yes |
| *biniou* | 12 | 9 | 9 | Yes | No |
| *GATA-A* | 11 | 11 | 8 | Yes | No |
| *Hand2-2* | 4 | 4 | 3 | Yes | No |
| *Sox8* | 7 | 7 | 6 | Yes | No |
| *piopio* | 4 | 3 | 3 | Yes | No |
| *Loc86968* | 4 | 3 | 1 | Yes | No |
| *Loc101009* | 3 | 2 | 2 | Yes | No |
| *Pax9-1* | 3 | 2 | 2 | Yes | No |
| *SDPEF* | 3 | 0 | 0 | Yes | No |
| *spaetzle* | 3 | 0 | 0 | Yes | No |
| *Loc95745* | 3 | 0 | 0 | Yes | No |
| *Mab21-1* | 4 | 4 | 1 | Yes | No |

**Table S2.** List of genomes referenced for synteny analysis.

| **Species** | **Taxon** | **GenBank asembly** |
| --- | --- | --- |
| *Ixodes scapularis* | apulmonate arachnids | GCF_016920785.2 |
| *Phalangium opilio* | apulmonate arachnids | GCA_019434445.1 |
| *Dysdera silvatica* | Arachnopulmonata | GCA_006491805.2 |
| *Parasteatoda tepidariorum* | Arachnopulmonata | GCF_000365465.3 |
| *Glomeris maerens* | Myriapoda | GCA_023279145.1 |
| *Rhysida immarginata* | Myriapoda | GCA_023313115.1 |
| *Thereuonema tuberculata* | Myriapoda | GCA_023159025.1 |
| *Niponia nodulosa* | Myriapoda | GCA_023159045.1 |
| *Eriocheir sinensis* | crustaceans | GCF_024679095.1 |
| *Daphnia magna* | crustaceans | GCF_020631705.1 |
| *Drosophila melanogaser* | Hexapoda | GCF_000001215.4 |
| *Tribolium castaneum* | Hexapoda | GCF_000002335.3 |
| *Epiperipatus broadwayi* | Onychophora | GCA_028023455.1 |
| *Ramazzottius varieornatus* | Tardigrada | GCA_001949185.1 |

**Table S3.** List of primer sequences used for gene cloning and/or riboprobe synthesis.

| **Primers** |  |  |  |  |  |
| --- | --- | --- | --- | --- | --- |
| All primers ordered with T7 linker sequences: | | |  |  |  |
| Forward primer:    ggccgcgg | |  |  |  |  |
| Reverse primer:    cccggggc | |  |  |  |  |
| Genes with Phenotypes | |  |  |  |  |
| **Species** | **Gene** | **Forward Primer** | **Reverse Primer** | **Length (bp)** | ***P. tepidariorum* source sequence** |
| *P. tepidariorum* | waist-less | CGTACGGAGATCGGTTTGAT | CCCATTTGGGCATAAAATTG | 978 | XM_016073862.1 |
| *P. tepidariorum* | waist-less 5' fragment | CGTACGGAGATCGGTTTGAT | TTCAGGGAATTTGGTTTCTGA | 473 | XM_016073862.1 |
| *P. tepidariorum* | waist-less 3' fragment | AGACTGCCACCTCAGAGAGC | CCCCATTGGGCATAAAATTG | 405 | XM_016073862.1 |
| *P. tepidariorum* | pannier-2 | ATGGACATGGAACTGCACAA | AAGTAGGCGGTGACGAAGAA | 842 | XM_016052354.2 |
| Genes used for expression assays | |  |  |  |  |
| **Species** | **Gene** | **Forward Primer** | **Reverse Primer** | **Length (bp)** | ***P. tepidariorum* source sequence** |
| *P. tepidariorum* | Scr-1 | AATTGCGAGGTTGTTCTTC | CAGACACAGGGCATGAGCTA | 750 | FM956097.1 |
| *P. tepidariorum* | Dll | ATGCCCAGGCTTACCCTATT | TGTCCCATGAGGAGATAGGC | 819 | FM876233.2 |
| *P. tepidariorum* | en | CTGCTTGACATTGCCTGAAA | GAATCTGCTGGCATTCCATT | 792 | AB125741.1 |
| Genes resulting in no phenotype | |  |  |  |  |
| **Species** | **Gene** | **Forward Primer** | **Reverse Primer** | **Length (bp)** | ***P. tepidariorum* source sequence** |
| *P. tepidariorum* | biniou | AAGGCCAGGAAAAGGACATT | ACTGTTTCGGAGGTCACAGG | 803 | XM_016074416.1 |
| *P. tepidariorum* | Hand2-2 | GGTCCTGGTGGTGAAACATT | ACGGGACCAACTCGTAATCA | 799 | XM_016061345.1 |
| *P. tepidariorum* | Mab21-1 | ATGGACATTTTGACGGCTTC | CGCGGATTTTCCTTTAAACA | 778 | XM_016057696.1 |
| *P. tepidariorum* | Pax9-1 | CCTGCGACATCTCAAGACAG | AACATTTCCTGTTGTCGCCTA | 790 | XM_016054880.1 |
| *P. tepidariorum* | piopio | TCATGTTCTCCACCCAGTCA | TCTTGAGGCACAGTTTGACG | 763 | XM_016048904.1 |
| *P. tepidariorum* | SAM_pointed_domain-containing_Ets_transcription_factor | ACAATGGCCGATTTTGAGAG | CAACGTCACAGGATTCATCG | 858 | XM_016067906.1 |
| *P. tepidariorum* | Sox8 | GTGGGGGTTCTCCCAGTAAT | AACATAAGGGTCTGCCGTTG | 859 | XM_016058966.1 |
| *P. tepidariorum* | spaetzle3 | TTTGTGGGAGAAAGGGAGAA | CCAGATGCGATTTCTTTCGT | 770 | XM_016052139.1 |
| *P. tepidariorum* | uncharacterized protein | TGCGAGTTCTTGGAATGATG | TCAGATTTTCTTAGTTCGGTGACA | 656 | XM_016063495.1 |
| *P. tepidariorum* | uncharacterized protein | ACACGGAGAATGTGCCTACC | CAGTGAAACCAAGCGAGACA | 795 | XM_016068055.2 |
| *P. tepidariorum* | uncharacterized protein | GACCCTGAAGCAATTTCTGG | CAAATCGCATGCAAATCTTTT | 573 | XM_016051911.1 |
| *P. tepidariorum* | GATA-A | ATTCCCATTGTCGAGTTTCG | AGCTCAGGTTTGACGATGCT | 851 | LC379619.1 |

**Table S4.** List of HCR probe sequences.

| **Initiator and gene name** | **Sequence** |
| --- | --- |
| B2_Ptep_waist-less | AAATCTTCTAATTTCGTCCTATTGGGT |
| B2_Ptep_waist-less | TTAGCTCTCACACCGACTTGGAATAAA |
| B2_Ptep_waist-less | AAGCTGGAGCACTTGTAAAACCAACGG |
| B2_Ptep_waist-less | GATGTAGAAACTGTATGATACATAGAA |
| B2_Ptep_waist-less | AACTGGCTGAAATGTCCTTATGTCTAG |
| B2_Ptep_waist-less | ATGAAAAAGAGGATGTCCTCAAGAAAA |
| B2_Ptep_waist-less | AAATAGTCTATAGGATTATGCCTCAGA |
| B2_Ptep_waist-less | ATGTCCCATTTGGGCATAAAATTGTAA |
| B2_Ptep_waist-less | AACTACCATTCAGAAGGCGATTGTAAT |
| B2_Ptep_waist-less | TGTGAACTGTCACCACAAGGTAAAAAA |
| B2_Ptep_waist-less | AAAGGAGGATGGTGACACAGAAGCTAA |
| B2_Ptep_waist-less | TGGGTGGTAAAACGGATGGGGATCCAA |
| B2_Ptep_waist-less | AAGAATTTGTCACCATGTGGGAAATTG |
| B2_Ptep_waist-less | ACTTCGCTCGTGGGAGGTGACACTAAA |
| B2_Ptep_waist-less | AAGACGATAAAGATGATTCGGCTGAAC |
| B2_Ptep_waist-less | AACTCTCCTGTTCTCCAAAGGGGTGAA |
| B2_Ptep_waist-less | AAGTGAAGGATGGTATAAGTGTTCATT |
| B2_Ptep_waist-less | CCTGCGGAGGAAGAGGCCTGTGGTAAA |
| B2_Ptep_waist-less | AAATCAGTTTGGTGGTCGAAGAACTTT |
| B2_Ptep_waist-less | ACCCAACCTTCTTAATTTTTGGTTCAA |
| B2_Ptep_waist-less | AAGCGCTAGTGACCAGATTTTTGGTTT |
| B2_Ptep_waist-less | GTGGTGGGCTCTCTGAGGTGGCAGTAA |
| B2_Ptep_waist-less | AAAAGTTCAGGGAATTTGGTTTCTGAG |
| B2_Ptep_waist-less | AATTGATTCTGATTTGATGTGTTCAAA |
| B2_Ptep_waist-less | AATCGGGAGCACTGAGGCTGTCACCAA |
| B2_Ptep_waist-less | TCAAGCGCTTTCATTGGAGACCTGTAA |
| B2_Ptep_waist-less | AACATCGTCCTTCATACTGTCTCGTCT |
| B2_Ptep_waist-less | GGGAGGTGCATTTCTGTTGTCGGTTAA |
| B2_Ptep_waist-less | AATTGATTTTCATTCGAATCGCTGAGT |
| B2_Ptep_waist-less | TCTACTGGAGTCACTTTCCTGGCATAA |
| B2_Ptep_waist-less | AATCATCTTTATCATCATCTTCATCCA |
| B2_Ptep_waist-less | CTCTTAAGGTCATCCCCACTCTGCTAA |
| B2_Ptep_waist-less | AAATCTCCGTACGGAGAATATGCGGCG |
| B2_Ptep_waist-less | TTTTTCACTATCAACAGGATCAAACAA |
| B2_Ptep_waist-less | AAAGCCCGAGGATCTTTTATGTCATAA |
| B2_Ptep_waist-less | GTGTCCTGGTGGTAGGGATGCCCAGAA |
| B2_Ptep_waist-less | AAGTGGCATGCTGTTTGAGGCTGGATA |
| B2_Ptep_waist-less | GATGGGCCTGACTTACTAGTGTGTCAA |
| B2_Ptep_waist-less | AATTGAGGAGATGCAGTATAGGCGAAA |
| B2_Ptep_waist-less | GCTGGGGTGCTGGCCTGTCATCAGAAA |
| **Initiator and gene name** | **Sequence** |
| B1_Ptep_pnr-2 | GAGGAGGGCAGCAAACGGAAGTATGGCGCGGGCATTCTATTTGAC |
| B1_Ptep_pnr-2 | TTAACCATCACGAATCTCATGTTTGTAGAAGAGTCTTCCTTTACG |
| B1_Ptep_pnr-2 | GAGGAGGGCAGCAAACGGAAGTCCATGCTGTGATGACTGTTATAG |
| B1_Ptep_pnr-2 | TTGGTGGTGATGGCTTGAACTGGAGTAGAAGAGTCTTCCTTTACG |
| B1_Ptep_pnr-2 | GAGGAGGGCAGCAAACGGAAAGCAGGGGTTGAATTGTTGCTTACT |
| B1_Ptep_pnr-2 | AGTTGCATCGCAAGATGCTGGAGAATAGAAGAGTCTTCCTTTACG |
| B1_Ptep_pnr-2 | GAGGAGGGCAGCAAACGGAATGGTGGTCTGTTTACCCCATGTAAT |
| B1_Ptep_pnr-2 | TTTTTGAATTCCTTCCTTTTTCATTTAGAAGAGTCTTCCTTTACG |
| B1_Ptep_pnr-2 | GAGGAGGGCAGCAAACGGAACAGATATTGCCGGATCTAGCAATCC |
| B1_Ptep_pnr-2 | CATCAGAACAATATGGATGTTGCAATAGAAGAGTCTTCCTTTACG |
| B1_Ptep_pnr-2 | GAGGAGGGCAGCAAACGGAAGTCAAAGCTGAAATTCCATAACCAG |
| B1_Ptep_pnr-2 | GAATTTTGAGCAAGATTCAGTCCAGTAGAAGAGTCTTCCTTTACG |
| B1_Ptep_pnr-2 | GAGGAGGGCAGCAAACGGAAAAGCAGCGCTGGAGGGAGATGATGG |
| B1_Ptep_pnr-2 | CGAGACTTCCAGTTGACGAAGCATATAGAAGAGTCTTCCTTTACG |
| B1_Ptep_pnr-2 | GAGGAGGGCAGCAAACGGAAATAGTTGGAGCACGTTCCCTGTTTC |
| B1_Ptep_pnr-2 | ACCATGACTGTATCACTGGAACTGCTAGAAGAGTCTTCCTTTACG |
| B1_Ptep_pnr-2 | GAGGAGGGCAGCAAACGGAAGTCGACTGTTCTCTATGATTGATGG |
| B1_Ptep_pnr-2 | CGACTATTAGATGTTTCATTGGCTGTAGAAGAGTCTTCCTTTACG |
| B1_Ptep_pnr-2 | GAGGAGGGCAGCAAACGGAATCATTGCGCAAGAAGAAGTATTCTC |
| B1_Ptep_pnr-2 | TTGCTAAGCTGTTGTTGTTATTCGATAGAAGAGTCTTCCTTTACG |
| B1_Ptep_pnr-2 | GAGGAGGGCAGCAAACGGAAAGCATGTTCTGTAACATTTGCATGG |
| B1_Ptep_pnr-2 | GTTACAATGATATGAATCTGAGACGTAGAAGAGTCTTCCTTTACG |
| B1_Ptep_pnr-2 | GAGGAGGGCAGCAAACGGAATGCTTTGATGGTCAGTTTTGGACAG |
| B1_Ptep_pnr-2 | TAGCATGGTGGTGTGAATCCCATTCTAGAAGAGTCTTCCTTTACG |
| B1_Ptep_pnr-2 | GAGGAGGGCAGCAAACGGAACAGTTCCATGTCCATCGTTTCTATT |
| B1_Ptep_pnr-2 | TTTTTGCCATTTGAGTAGTTTCTTGTAGAAGAGTCTTCCTTTACG |
| B1_Ptep_pnr-2 | GAGGAGGGCAGCAAACGGAATAATCTTCAACAGCGAGAACATTTG |
| B1_Ptep_pnr-2 | TTTTTGACTGCTGGAGGAACTAAACTAGAAGAGTCTTCCTTTACG |
| B1_Ptep_pnr-2 | GAGGAGGGCAGCAAACGGAAATTACTGTCATCTCTTGATACGGTT |
| B1_Ptep_pnr-2 | TTGTACTGCTTTATTCACTCCCGCGTAGAAGAGTCTTCCTTTACG |
| B1_Ptep_pnr-2 | GAGGAGGGCAGCAAACGGAAAAGATTCTTTGGGAAGAACGGGCTG |
| B1_Ptep_pnr-2 | TACCAAAATCATTTACATTACTGTCTAGAAGAGTCTTCCTTTACG |
| B1_Ptep_pnr-2 | GAGGAGGGCAGCAAACGGAAAATGTGCTGGCTCCAGCACTAAATA |
| B1_Ptep_pnr-2 | GTATAGAATTTTTGGTCATAGAATTTAGAAGAGTCTTCCTTTACG |
| B1_Ptep_pnr-2 | GAGGAGGGCAGCAAACGGAAAAATTATGGGCAGAAGTAAATACGG |
| B1_Ptep_pnr-2 | TGTGCTTCATGATTGTTAACTAACGTAGAAGAGTCTTCCTTTACG |
| B1_Ptep_pnr-2 | GAGGAGGGCAGCAAACGGAACTAAAGTTTCTGGCTTCATATTGCG |
| B1_Ptep_pnr-2 | CAAAGTTCTTATAAGACAAGTTGCGTAGAAGAGTCTTCCTTTACG |
| B1_Ptep_pnr-2 | GAGGAGGGCAGCAAACGGAACTTTCGACGATTTGAGGATAGGATT |
| B1_Ptep_pnr-2 | ATGCTGTTTAAATTCTTCGAAGCTTTAGAAGAGTCTTCCTTTACG |
| B1_Ptep_pnr-2 | GAGGAGGGCAGCAAACGGAACTTTCGACGATTTGAGGATAGGATT |
| B1_Ptep_pnr-2 | ATGCTGTTTAAATTCTTCGAAGCTTTAGAAGAGTCTTCCTTTACG |
| B1_Ptep_pnr-2 | GAGGAGGGCAGCAAACGGAACTTTCGACGATTTGAGGATAGGATT |
| B1_Ptep_pnr-2 | ATGCTGTTTAAATTCTTCGAAGCTTTAGAAGAGTCTTCCTTTACG |
| B1_Ptep_pnr-2 | GAGGAGGGCAGCAAACGGAACTTTCGACGATTTGAGGATAGGATT |
| B1_Ptep_pnr-2 | ATGCTGTTTAAATTCTTCGAAGCTTTAGAAGAGTCTTCCTTTACG |
| **Initiator and gene name** | **Sequence** |
| B3_Ptep_sog | GTCCCTGCCTCTATATCTTTGTGTATAAAAGCATAGCACCACCGA |
| B3_Ptep_sog | TTTTTGTTATAAAATGGGTAGCCCATTCCACTCAACTTTAACCCG |
| B3_Ptep_sog | GTCCCTGCCTCTATATCTTTCTCAGTTTTGTTTCTCTTTGGGTTG |
| B3_Ptep_sog | CAACAAATGCCTGTTGGATGATGGATTCCACTCAACTTTAACCCG |
| B3_Ptep_sog | GTCCCTGCCTCTATATCTTTGACACAGAATCTATGCATCTATCAC |
| B3_Ptep_sog | CCTTCCGCGGATCTTGAAGACGATTTTCCACTCAACTTTAACCCG |
| B3_Ptep_sog | GTCCCTGCCTCTATATCTTTATGACATTTAACACATTTTTCAGCT |
| B3_Ptep_sog | CTTACACTTGGCTCGTCCGTCTTTATTCCACTCAACTTTAACCCG |
| B3_Ptep_sog | GTCCCTGCCTCTATATCTTTTAACAGTTGATTTTTCACCACTTTG |
| B3_Ptep_sog | TGCAACCTCCAGCATTCAATATCTCTTCCACTCAACTTTAACCCG |
| B3_Ptep_sog | GTCCCTGCCTCTATATCTTTCTATAAGCATCTTTCTCAGCACAAG |
| B3_Ptep_sog | TTTTTGCAACAATCGTTATCTTGCTTTCCACTCAACTTTAACCCG |
| B3_Ptep_sog | GTCCCTGCCTCTATATCTTTGGGAGGAACATAGGGATGCCATCTG |
| B3_Ptep_sog | ACAAAGAGAGCATTTGCTGAATCCATTCCACTCAACTTTAACCCG |
| B3_Ptep_sog | GTCCCTGCCTCTATATCTTTAACATTCTCCTTCCATCGTCATAGG |
| B3_Ptep_sog | CATTAGAGTTACTTGGGCAGAAGGGTTCCACTCAACTTTAACCCG |
| B3_Ptep_sog | GTCCCTGCCTCTATATCTTTTCACCTTTCCAATGATCTCCATCAT |
| B3_Ptep_sog | CAAGAGCACATGGTGCATTCTTCATTTCCACTCAACTTTAACCCG |
| B3_Ptep_sog | GTCCCTGCCTCTATATCTTTTCTTGTCTTGGAGGATGCATCGTAA |
| B3_Ptep_sog | GTATCCATCCTGATATGTATTCCCATTCCACTCAACTTTAACCCG |
| B3_Ptep_sog | GTCCCTGCCTCTATATCTTTAGTGTAGCCTTGTTGTATGAAAGCC |
| B3_Ptep_sog | GTTAAGTTTCGATGTCACAACCAGGTTCCACTCAACTTTAACCCG |
| B3_Ptep_sog | GTCCCTGCCTCTATATCTTTTGATATTTTCCTGGTCTCACTTGCA |
| B3_Ptep_sog | TTTTTCAAGACACGCTGTAGAGGGTTTCCACTCAACTTTAACCCG |
| B3_Ptep_sog | GTCCCTGCCTCTATATCTTTCCAAGCTATTCCACCTGCAGTTGTG |
| B3_Ptep_sog | AAGAATGCAATCTTTATCAATGGATTTCCACTCAACTTTAACCCG |
| B3_Ptep_sog | GTCCCTGCCTCTATATCTTTGAACTTTGTCAGAACGTGTAACACG |
| B3_Ptep_sog | GATGAACTTGGCCTCGCAACTGCATTTCCACTCAACTTTAACCCG |
| B3_Ptep_sog | GTCCCTGCCTCTATATCTTTACTGTATTAGGTGTGTCTGCAAGTA |
| B3_Ptep_sog | CGATATTCTGGAGTAGCTGAATGTATTCCACTCAACTTTAACCCG |
| B3_Ptep_sog | GTCCCTGCCTCTATATCTTTTAGGTTTTGTGAAACTTGTTGTAAC |
| B3_Ptep_sog | GAATGTGCCATTTCCCCATGAATTATTCCACTCAACTTTAACCCG |
| B3_Ptep_sog | GTCCCTGCCTCTATATCTTTACTTATAATTTATGAATCCTTCTGA |
| B3_Ptep_sog | ATTCAATGCTTGGATCGCTGACGTGTTCCACTCAACTTTAACCCG |
| B3_Ptep_sog | GTCCCTGCCTCTATATCTTTCATGATGCGTCCGCTCTGGAGGCGT |
| B3_Ptep_sog | TTCAAACACGTTGCAAGTAACCTTGTTCCACTCAACTTTAACCCG |
| B3_Ptep_sog | GTCCCTGCCTCTATATCTTTATCAATGGCACATCTCGGATATCTT |
| B3_Ptep_sog | TCCATATCTTCATAGAAAAGGCTAATTCCACTCAACTTTAACCCG |
| B3_Ptep_sog | GTCCCTGCCTCTATATCTTTAGTTTCATTCGCCAAATCGCCAGAT |
| B3_Ptep_sog | AATGTTCGCCATACCACCAGTACCTTTCCACTCAACTTTAACCCG |
| B3_Ptep_sog | GTCCCTGCCTCTATATCTTTTGTGAGCTAGTTACCAATATAAAGA |
| B3_Ptep_sog | ATTCTCCCTCCAACAATTCCATCAGTTCCACTCAACTTTAACCCG |
| B3_Ptep_sog | GTCCCTGCCTCTATATCTTTGGGTGCAATGGGAATTTCATGTTCT |
| B3_Ptep_sog | ACAAACTTTTGATCCTTGAACATGGTTCCACTCAACTTTAACCCG |
| B3_Ptep_sog | GTCCCTGCCTCTATATCTTTAAAACAAAGTAACCACGTGCTGCTC |
| B3_Ptep_sog | ATGGTATAATGAAGATCTCGTTTAGTTCCACTCAACTTTAACCCG |
| B3_Ptep_sog | GTCCCTGCCTCTATATCTTTCTTCATCTTCTGATTCTAGCTTCCG |
| B3_Ptep_sog | GAGCCGTATATTCTTTCAGAATTTTTTCCACTCAACTTTAACCCG |
| B3_Ptep_sog | GTCCCTGCCTCTATATCTTTCATGTTGGTTTCGGGCAATCATTCT |
| B3_Ptep_sog | CTTTGGGGAAGTAAGACAGGATCGTTTCCACTCAACTTTAACCCG |
| B3_Ptep_sog | GTCCCTGCCTCTATATCTTTTTCTCCTTTTTCGCTGAACCGGTAT |
| B3_Ptep_sog | TATTTTTGCATCGCACTTTCGACATTTCCACTCAACTTTAACCCG |
| B3_Ptep_sog | GTCCCTGCCTCTATATCTTTAAATGGAGGACCCAAGTCAGGCCTC |
| B3_Ptep_sog | TTCGCATCGCATGCAATACAGCATGTTCCACTCAACTTTAACCCG |
| B3_Ptep_sog | GTCCCTGCCTCTATATCTTTCCAAACTGACAATGAGTTTGTTTTT |
| B3_Ptep_sog | CTCTCCTCCAGTTCGTAAGTATTGTTTCCACTCAACTTTAACCCG |
| B3_Ptep_sog | GTCCCTGCCTCTATATCTTTATTCATTTTTGAATAGAGTTGCTTC |
| B3_Ptep_sog | CAATCCACACACACTTTGTTTCCGTTTCCACTCAACTTTAACCCG |
| B3_Ptep_sog | GTCCCTGCCTCTATATCTTTTCTAACAACTAAAAACATACGGCAC |
| B3_Ptep_sog | GTTTTCCCCTAGATTTTCTTTCGCATTCCACTCAACTTTAACCCG |
| **Initiator and gene name** | **Sequence** |
| B1_Ptep_Irx1_multiplexed | GAGGAGGGCAGCAAACGGAATCCAGCGCTTATGACAGTGCTACCC |
| B1_Ptep_Irx1_multiplexed | TTATGATAAAACACGACCTTGTGTCTAGAAGAGTCTTCCTTTACG |
| B1_Ptep_Irx1_multiplexed | GAGGAGGGCAGCAAACGGAAGAGGTTAGGTTGAACACTGAGGGAG |
| B1_Ptep_Irx1_multiplexed | GGTTGCAGAGAGTTGTTGCTCAGTTTAGAAGAGTCTTCCTTTACG |
| B1_Ptep_Irx1_multiplexed | GAGGAGGGCAGCAAACGGAACCGTAATCTATTCAGTGAGGACACT |
| B1_Ptep_Irx1_multiplexed | AGCAGTCGGAAAGGCCGCTGCAGCTTAGAAGAGTCTTCCTTTACG |
| B1_Ptep_Irx1_multiplexed | GAGGAGGGCAGCAAACGGAAGGATCCTCCGACACTGGTCACAACA |
| B1_Ptep_Irx1_multiplexed | ACACGAGTAAACGGCCGACAGAGGATAGAAGAGTCTTCCTTTACG |
| B1_Ptep_Irx1_multiplexed | GAGGAGGGCAGCAAACGGAAAGCACCAGGCGGTAGAGCAGACGAA |
| B1_Ptep_Irx1_multiplexed | TCTCTGAAGAGGACTCAGTCTGGAATAGAAGAGTCTTCCTTTACG |
| B1_Ptep_Irx1_multiplexed | GAGGAGGGCAGCAAACGGAAACAATGGACCATATTCGAGGTTTTT |
| B1_Ptep_Irx1_multiplexed | GGCCCTTGTGTTGACGTAGCCGTGTTAGAAGAGTCTTCCTTTACG |
| B1_Ptep_Irx1_multiplexed | GAGGAGGGCAGCAAACGGAATCGAAGCTGGGTGGGGTAGTGAAGA |
| B1_Ptep_Irx1_multiplexed | GCTGGCTACTTTGTGGTGGTAGTTCTAGAAGAGTCTTCCTTTACG |
| B1_Ptep_Irx1_multiplexed | GAGGAGGGCAGCAAACGGAACAATGAGGCATCAGAGATAGGACTT |
| B1_Ptep_Irx1_multiplexed | CTGCGAGGATTCAGAGTCTTCTGATTAGAAGAGTCTTCCTTTACG |
| B1_Ptep_Irx1_multiplexed | GAGGAGGGCAGCAAACGGAATTCTCTTTGTCGGAGCTGTTTGTGA |
| B1_Ptep_Irx1_multiplexed | ACACGATGCGTCTTCAGGAGGGAACTAGAAGAGTCTTCCTTTACG |
| B1_Ptep_Irx1_multiplexed | GAGGAGGGCAGCAAACGGAAACGACTTGCAGTCATCATCCACGTC |
| B1_Ptep_Irx1_multiplexed | TGTGGCCGTCTGTACTCGTAGATGCTAGAAGAGTCTTCCTTTACG |
| B1_Ptep_Irx1_multiplexed | GAGGAGGGCAGCAAACGGAATTGCCTAGGATGGTGATGATCGTCC |
| B1_Ptep_Irx1_multiplexed | GAAATCCATCTTACCATCGTCGGAATAGAAGAGTCTTCCTTTACG |
| B1_Ptep_Irx1_multiplexed | GAGGAGGGCAGCAAACGGAAGATCTTTTACCATCGGACTCTTTTA |
| B1_Ptep_Irx1_multiplexed | CTCCGATCTACATCGGAATCTTTATTAGAAGAGTCTTCCTTTACG |
| B1_Ptep_Irx1_multiplexed | GAGGAGGGCAGCAAACGGAACCGTTTGGAAAGGGATAACCAGCAA |
| B1_Ptep_Irx1_multiplexed | TTTTTACCGTTTAAATCAACACCATTAGAAGAGTCTTCCTTTACG |
| B1_Ptep_Irx1_multiplexed | GAGGAGGGCAGCAAACGGAAAAGCTGCATATGGTAAACTTCTCCA |
| B1_Ptep_Irx1_multiplexed | CGGCCGAATCGTATGGGTAATACATTAGAAGAGTCTTCCTTTACG |
| B1_Ptep_Irx1_multiplexed | GAGGAGGGCAGCAAACGGAAATTAGCCGTTGGAGAATAGAATGCG |
| B1_Ptep_Irx1_multiplexed | TTCTAGGTTGTCCTTTAGGTCAAAGTAGAAGAGTCTTCCTTTACG |
| B1_Ptep_Irx1_multiplexed | GAGGAGGGCAGCAAACGGAAGGGTGGTTGCAGTTGATCCGTTATA |
| B1_Ptep_Irx1_multiplexed | TTCCGGACTGTCCAGCCATCAGCAGTAGAAGAGTCTTCCTTTACG |
| B1_Ptep_Irx1_multiplexed | GAGGAGGGCAGCAAACGGAAAGATGGACAGTCATACGTATGGATG |
| B1_Ptep_Irx1_multiplexed | AAATTGGGGACATGACATTTTCAAGTAGAAGAGTCTTCCTTTACG |
| B1_Ptep_Irx1_multiplexed | GAGGAGGGCAGCAAACGGAATAGAAGTCCATACAATGTGTCTCTC |
| B1_Ptep_Irx1_multiplexed | GGTCGTCCGGTTCTCCAGGAAATCCTAGAAGAGTCTTCCTTTACG |
| B1_Ptep_Irx1_multiplexed | GAGGAGGGCAGCAAACGGAATCGTTTCATTAAGCCATGCAATTAC |
| B1_Ptep_Irx1_multiplexed | TTACCACACTAGGTTTCTGAAGTCATAGAAGAGTCTTCCTTTACG |
| B1_Ptep_Irx1_multiplexed | GAGGAGGGCAGCAAACGGAATATTTGAATTCCAAACTATAAGACC |
| B1_Ptep_Irx1_multiplexed | TTGCGTCGAAATAAACGAATCGGAATAGAAGAGTCTTCCTTTACG |
| B2_Ptep_Irx3-1_multiplexed | CCTCGTAAATCCTCATCAAATAACGAGGCGGAATGTTCGAGAACC |
| B2_Ptep_Irx3-1_multiplexed | TTATTCACTATGTTTAACATTCGTTAAATCATCCAGTAAACCGCC |
| B2_Ptep_Irx3-1_multiplexed | CCTCGTAAATCCTCATCAAAAATCCGTTCGAATCGAAAAATTCCG |
| B2_Ptep_Irx3-1_multiplexed | TGACAGCACTTTTGAAGTGTTCTATAAATCATCCAGTAAACCGCC |
| B2_Ptep_Irx3-1_multiplexed | CCTCGTAAATCCTCATCAAATGATTTGTTGTCATACTCTAAACAG |
| B2_Ptep_Irx3-1_multiplexed | TGTGTCACAATCCGCTTCGTAAACTAAATCATCCAGTAAACCGCC |
| B2_Ptep_Irx3-1_multiplexed | CCTCGTAAATCCTCATCAAATTCTGAGGCACACTTTATATCATTC |
| B2_Ptep_Irx3-1_multiplexed | GGGGTTGTCCATCATGCACGATCCTAAATCATCCAGTAAACCGCC |
| B2_Ptep_Irx3-1_multiplexed | CCTCGTAAATCCTCATCAAAGGCATGAAATCTGTAAAAGAGGACA |
| B2_Ptep_Irx3-1_multiplexed | TTTTTATTGCAACACAACTCACTGTAAATCATCCAGTAAACCGCC |
| B2_Ptep_Irx3-1_multiplexed | CCTCGTAAATCCTCATCAAAGGACATTTTCTAAACAGAAATCCAT |
| B2_Ptep_Irx3-1_multiplexed | TCGAGTTCACGGAACACTCTGCTTTAAATCATCCAGTAAACCGCC |
| B2_Ptep_Irx3-1_multiplexed | CCTCGTAAATCCTCATCAAACAAGCGACCAAATTTTTGGTCTGCT |
| B2_Ptep_Irx3-1_multiplexed | GGGGACTATTGGATGTGGCAGTTCTAAATCATCCAGTAAACCGCC |
| B2_Ptep_Irx3-1_multiplexed | CCTCGTAAATCCTCATCAAAGGAAACGTCTTTTGCACTAGGTTTT |
| B2_Ptep_Irx3-1_multiplexed | GGAATCGAAGGTTTCCGAAAACCTAAAATCATCCAGTAAACCGCC |
| B2_Ptep_Irx3-1_multiplexed | CCTCGTAAATCCTCATCAAATCCTAACTGGGCTAGAATGTTCTGA |
| B2_Ptep_Irx3-1_multiplexed | CCTGCTGGAATCCCGGGTAGCAGGAAAATCATCCAGTAAACCGCC |
| B2_Ptep_Irx3-1_multiplexed | CCTCGTAAATCCTCATCAAAAGTTCGATAACATTCCAGAGCACAT |
| B2_Ptep_Irx3-1_multiplexed | GATGGAATCTTCTGATGAAAGACTTAAATCATCCAGTAAACCGCC |
| B2_Ptep_Irx3-1_multiplexed | CCTCGTAAATCCTCATCAAAAAGGTTCTTTGCTGTGGAATGTTTC |
| B2_Ptep_Irx3-1_multiplexed | GGGGAGATGTGGAATAATCGGCAGGAAATCATCCAGTAAACCGCC |
| B2_Ptep_Irx3-1_multiplexed | CCTCGTAAATCCTCATCAAACTTCTGCTGTATCGTCTGACTTCAG |
| B2_Ptep_Irx3-1_multiplexed | AAAAATCGTTCAAGTTGCTGCACTCAAATCATCCAGTAAACCGCC |
| B2_Ptep_Irx3-1_multiplexed | CCTCGTAAATCCTCATCAAACTGTCTGTATAGGTTGTACAGGAGG |
| B2_Ptep_Irx3-1_multiplexed | TTTTTAACACTCTCCATAGAACCGTAAATCATCCAGTAAACCGCC |
| B2_Ptep_Irx3-1_multiplexed | CCTCGTAAATCCTCATCAAATCCATGAGTTTCGTGATTCTTTCAA |
| B2_Ptep_Irx3-1_multiplexed | TAGGATCGTATGGGTAACAGGATGCAAATCATCCAGTAAACCGCC |
| B2_Ptep_Irx3-1_multiplexed | CCTCGTAAATCCTCATCAAAAGTGGTTGTGTTGGCTGCCAGTGAT |
| B2_Ptep_Irx3-1_multiplexed | GTAAGGAGCACCCAGTGTTGAGTAGAAATCATCCAGTAAACCGCC |
| B2_Ptep_Irx3-1_multiplexed | CCTCGTAAATCCTCATCAAAAGACCGAAGGATGCCAGCCGAGGTA |
| B2_Ptep_Irx3-1_multiplexed | ATGTGATCAGCGTATGAAGAACCGTAAATCATCCAGTAAACCGCC |
| B2_Ptep_Irx3-1_multiplexed | CCTCGTAAATCCTCATCAAATACCTGTGTTGAAGAAGGTGGATAT |
| B2_Ptep_Irx3-1_multiplexed | GCAACACGGTCCGCCGCTGCTCACCAAATCATCCAGTAAACCGCC |
| B2_Ptep_Irx3-1_multiplexed | CCTCGTAAATCCTCATCAAACGATTTGGACGGATTAGTGGGGTGT |
| B2_Ptep_Irx3-1_multiplexed | TAACCGAATTGTGAATAGGACATTAAAATCATCCAGTAAACCGCC |
| B2_Ptep_Irx3-1_multiplexed | CCTCGTAAATCCTCATCAAATCATTCGGCCTTTCTTGAACCGATC |
| B2_Ptep_Irx3-1_multiplexed | GAAAGGGAGATCGGCCTCACAGTGTAAATCATCCAGTAAACCGCC |
| B2_Ptep_Irx3-1_multiplexed | CCTCGTAAATCCTCATCAAAATTCCGTTATTCCAACTGTATCCCT |
| B2_Ptep_Irx3-1_multiplexed | CAAAAACATGGCGATCATCGGTCAAAAATCATCCAGTAAACCGCC |
| B3_Ptep_waist-less_multiplexed | GTCCCTGCCTCTATATCTTTATCTTCTAATTTCGTCCTATTGGGT |
| B3_Ptep_waist-less_multiplexed | TTAGCTCTCACACCGACTTGGAATATTCCACTCAACTTTAACCCG |
| B3_Ptep_waist-less_multiplexed | GTCCCTGCCTCTATATCTTTGCTGGAGCACTTGTAAAACCAACGG |
| B3_Ptep_waist-less_multiplexed | GATGTAGAAACTGTATGATACATAGTTCCACTCAACTTTAACCCG |
| B3_Ptep_waist-less_multiplexed | GTCCCTGCCTCTATATCTTTCTGGCTGAAATGTCCTTATGTCTAG |
| B3_Ptep_waist-less_multiplexed | ATGAAAAAGAGGATGTCCTCAAGAATTCCACTCAACTTTAACCCG |
| B3_Ptep_waist-less_multiplexed | GTCCCTGCCTCTATATCTTTATAGTCTATAGGATTATGCCTCAGA |
| B3_Ptep_waist-less_multiplexed | ATGTCCCATTTGGGCATAAAATTGTTTCCACTCAACTTTAACCCG |
| B3_Ptep_waist-less_multiplexed | GTCCCTGCCTCTATATCTTTCTACCATTCAGAAGGCGATTGTAAT |
| B3_Ptep_waist-less_multiplexed | TGTGAACTGTCACCACAAGGTAAAATTCCACTCAACTTTAACCCG |
| B3_Ptep_waist-less_multiplexed | GTCCCTGCCTCTATATCTTTAGGAGGATGGTGACACAGAAGCTAA |
| B3_Ptep_waist-less_multiplexed | TGGGTGGTAAAACGGATGGGGATCCTTCCACTCAACTTTAACCCG |
| B3_Ptep_waist-less_multiplexed | GTCCCTGCCTCTATATCTTTGAATTTGTCACCATGTGGGAAATTG |
| B3_Ptep_waist-less_multiplexed | ACTTCGCTCGTGGGAGGTGACACTATTCCACTCAACTTTAACCCG |
| B3_Ptep_waist-less_multiplexed | GTCCCTGCCTCTATATCTTTGACGATAAAGATGATTCGGCTGAAC |
| B3_Ptep_waist-less_multiplexed | AACTCTCCTGTTCTCCAAAGGGGTGTTCCACTCAACTTTAACCCG |
| B3_Ptep_waist-less_multiplexed | GTCCCTGCCTCTATATCTTTGTGAAGGATGGTATAAGTGTTCATT |
| B3_Ptep_waist-less_multiplexed | CCTGCGGAGGAAGAGGCCTGTGGTATTCCACTCAACTTTAACCCG |
| B3_Ptep_waist-less_multiplexed | GTCCCTGCCTCTATATCTTTATCAGTTTGGTGGTCGAAGAACTTT |
| B3_Ptep_waist-less_multiplexed | ACCCAACCTTCTTAATTTTTGGTTCTTCCACTCAACTTTAACCCG |
| B3_Ptep_waist-less_multiplexed | GTCCCTGCCTCTATATCTTTGCGCTAGTGACCAGATTTTTGGTTT |
| B3_Ptep_waist-less_multiplexed | GTGGTGGGCTCTCTGAGGTGGCAGTTTCCACTCAACTTTAACCCG |
| B3_Ptep_waist-less_multiplexed | GTCCCTGCCTCTATATCTTTAAGTTCAGGGAATTTGGTTTCTGAG |
| B3_Ptep_waist-less_multiplexed | AATTGATTCTGATTTGATGTGTTCATTCCACTCAACTTTAACCCG |
| B3_Ptep_waist-less_multiplexed | GTCCCTGCCTCTATATCTTTTCGGGAGCACTGAGGCTGTCACCAA |
| B3_Ptep_waist-less_multiplexed | TCAAGCGCTTTCATTGGAGACCTGTTTCCACTCAACTTTAACCCG |
| B3_Ptep_waist-less_multiplexed | GTCCCTGCCTCTATATCTTTCATCGTCCTTCATACTGTCTCGTCT |
| B3_Ptep_waist-less_multiplexed | GGGAGGTGCATTTCTGTTGTCGGTTTTCCACTCAACTTTAACCCG |
| B3_Ptep_waist-less_multiplexed | GTCCCTGCCTCTATATCTTTTTGATTTTCATTCGAATCGCTGAGT |
| B3_Ptep_waist-less_multiplexed | TCTACTGGAGTCACTTTCCTGGCATTTCCACTCAACTTTAACCCG |
| B3_Ptep_waist-less_multiplexed | GTCCCTGCCTCTATATCTTTTCATCTTTATCATCATCTTCATCCA |
| B3_Ptep_waist-less_multiplexed | CTCTTAAGGTCATCCCCACTCTGCTTTCCACTCAACTTTAACCCG |
| B3_Ptep_waist-less_multiplexed | GTCCCTGCCTCTATATCTTTATCTCCGTACGGAGAATATGCGGCG |
| B3_Ptep_waist-less_multiplexed | TTTTTCACTATCAACAGGATCAAACTTCCACTCAACTTTAACCCG |
| B3_Ptep_waist-less_multiplexed | GTCCCTGCCTCTATATCTTTAGCCCGAGGATCTTTTATGTCATAA |
| B3_Ptep_waist-less_multiplexed | GTGTCCTGGTGGTAGGGATGCCCAGTTCCACTCAACTTTAACCCG |
| B3_Ptep_waist-less_multiplexed | GTCCCTGCCTCTATATCTTTGTGGCATGCTGTTTGAGGCTGGATA |
| B3_Ptep_waist-less_multiplexed | GATGGGCCTGACTTACTAGTGTGTCTTCCACTCAACTTTAACCCG |
| B3_Ptep_waist-less_multiplexed | GTCCCTGCCTCTATATCTTTTTGAGGAGATGCAGTATAGGCGAAA |
| B3_Ptep_waist-less_multiplexed | GCTGGGGTGCTGGCCTGTCATCAGATTCCACTCAACTTTAACCCG |
| **Initiator and gene name** | **Sequence** |
| B1_Ptep_Irx2-1_multiplexed | GAGGAGGGCAGCAAACGGAATCCCGTTGCAGAATAATCCTCAGGT |
| B1_Ptep_Irx2-1_multiplexed | TTATTGCAGGGATGTCGCCTTGAAATAGAAGAGTCTTCCTTTACG |
| B1_Ptep_Irx2-1_multiplexed | GAGGAGGGCAGCAAACGGAACACCGATAGCAGATGATTGATATAG |
| B1_Ptep_Irx2-1_multiplexed | TACTGTTTTGAATACTAGCAGATACTAGAAGAGTCTTCCTTTACG |
| B1_Ptep_Irx2-1_multiplexed | GAGGAGGGCAGCAAACGGAATGAGCCGGATACCTTCATGTTTGGT |
| B1_Ptep_Irx2-1_multiplexed | ACCACCTTGTCCACCCAGGAAGCCATAGAAGAGTCTTCCTTTACG |
| B1_Ptep_Irx2-1_multiplexed | GAGGAGGGCAGCAAACGGAAGGCACAAAACCTTGAGCAGATGAAC |
| B1_Ptep_Irx2-1_multiplexed | GTTTGAGGTGGTGTGTCCGTCTGAGTAGAAGAGTCTTCCTTTACG |
| B1_Ptep_Irx2-1_multiplexed | GAGGAGGGCAGCAAACGGAAAGGATGATAACTTCCAAAGCCACCT |
| B1_Ptep_Irx2-1_multiplexed | GGGCGATGCGAATGACCCACCAGAGTAGAAGAGTCTTCCTTTACG |
| B1_Ptep_Irx2-1_multiplexed | GAGGAGGGCAGCAAACGGAAGGATTCACCGAAAACCATCCACAGT |
| B1_Ptep_Irx2-1_multiplexed | GATGCTGAGGCATTGGTCACTTGGTTAGAAGAGTCTTCCTTTACG |
| B1_Ptep_Irx2-1_multiplexed | GAGGAGGGCAGCAAACGGAATCTCTTGAGTTGGACCACCAAGCAG |
| B1_Ptep_Irx2-1_multiplexed | CTAACATGTCCATCATACTGCCGGATAGAAGAGTCTTCCTTTACG |
| B1_Ptep_Irx2-1_multiplexed | GAGGAGGGCAGCAAACGGAACCAAAGACCATATTTTTGGTTTATT |
| B1_Ptep_Irx2-1_multiplexed | GGGGACTTTTACTCGTGGCTGTGTCTAGAAGAGTCTTCCTTTACG |
| B1_Ptep_Irx2-1_multiplexed | GAGGAGGGCAGCAAACGGAACTTCGACAGCAGAAGGCGAGCTTTT |
| B1_Ptep_Irx2-1_multiplexed | TCATGGATCGACCGGACATGTCCACTAGAAGAGTCTTCCTTTACG |
| B1_Ptep_Irx2-1_multiplexed | GAGGAGGGCAGCAAACGGAAAATTAGGCGATGAGGATGAGTCAGT |
| B1_Ptep_Irx2-1_multiplexed | TGCGCATGAGGGCCTGAATGTCCGGTAGAAGAGTCTTCCTTTACG |
| B1_Ptep_Irx2-1_multiplexed | GAGGAGGGCAGCAAACGGAAAGTACTGTCCCTAGTACTGTTTTCT |
| B1_Ptep_Irx2-1_multiplexed | TGTGAATTCGGAGGTAGCGATGAGGTAGAAGAGTCTTCCTTTACG |
| B1_Ptep_Irx2-1_multiplexed | GAGGAGGGCAGCAAACGGAATTTCTGTGGTGAGGCAGGTGGGGAT |
| B1_Ptep_Irx2-1_multiplexed | TGTTGAGCACCGTTTGGAGTTATTTTAGAAGAGTCTTCCTTTACG |
| B1_Ptep_Irx2-1_multiplexed | GAGGAGGGCAGCAAACGGAACGTAATGTACTGCACCATCAATGGA |
| B1_Ptep_Irx2-1_multiplexed | GTCGTCCAGTTATGTGATTACCTGATAGAAGAGTCTTCCTTTACG |
| B1_Ptep_Irx2-1_multiplexed | GAGGAGGGCAGCAAACGGAATTCGTCTTCGCTTTCACATTTTGTC |
| B1_Ptep_Irx2-1_multiplexed | ATCACACGATGCTGTTCTACGATCGTAGAAGAGTCTTCCTTTACG |
| B1_Ptep_Irx2-1_multiplexed | GAGGAGGGCAGCAAACGGAACTGTCCATCCTGTTCCTTTTAGTTG |
| B1_Ptep_Irx2-1_multiplexed | CTGGACTCCCCATCAACGCGATCGTTAGAAGAGTCTTCCTTTACG |
| B1_Ptep_Irx2-1_multiplexed | GAGGAGGGCAGCAAACGGAACATCGTCATAATCCTTGACGTCATT |
| B1_Ptep_Irx2-1_multiplexed | CATCACCGTCATCTTCGACGTCATCTAGAAGAGTCTTCCTTTACG |
| B1_Ptep_Irx2-1_multiplexed | GAGGAGGGCAGCAAACGGAATACCCGTAAGCAGCCAGGCTTGGAT |
| B1_Ptep_Irx2-1_multiplexed | TTTTTGCCATTGAAATCTAAACCACTAGAAGAGTCTTCCTTTACG |
| B1_Ptep_Irx2-1_multiplexed | GAGGAGGGCAGCAAACGGAACTCTCCACGGGTCGCCAGCATCTTT |
| B1_Ptep_Irx2-1_multiplexed | AATAATAAGAAGCAGATGGCGTAATTAGAAGAGTCTTCCTTTACG |
| B1_Ptep_Irx2-1_multiplexed | GAGGAGGGCAGCAAACGGAAAAAGGCTGAAGATTCAACTCCCGTG |
| B1_Ptep_Irx2-1_multiplexed | ACCATAGGCTGTACTCAAAGAGGGATAGAAGAGTCTTCCTTTACG |
| B1_Ptep_Irx2-1_multiplexed | GAGGAGGGCAGCAAACGGAACTAGGATGGTAGGTTGTAAGTAGCC |
| B1_Ptep_Irx2-1_multiplexed | TTGGGATAGGCTTGATGCTCGTAGATAGAAGAGTCTTCCTTTACG |
| B2_Ptep_Irx3_multiplexed | CCTCGTAAATCCTCATCAAATAACGAGGCGGAATGTTCGAGAACC |
| B2_Ptep_Irx3_multiplexed | TTATTCACTATGTTTAACATTCGTTAAATCATCCAGTAAACCGCC |
| B2_Ptep_Irx3_multiplexed | CCTCGTAAATCCTCATCAAAAATCCGTTCGAATCGAAAAATTCCG |
| B2_Ptep_Irx3_multiplexed | TGACAGCACTTTTGAAGTGTTCTATAAATCATCCAGTAAACCGCC |
| B2_Ptep_Irx3_multiplexed | CCTCGTAAATCCTCATCAAATGATTTGTTGTCATACTCTAAACAG |
| B2_Ptep_Irx3_multiplexed | TGTGTCACAATCCGCTTCGTAAACTAAATCATCCAGTAAACCGCC |
| B2_Ptep_Irx3_multiplexed | CCTCGTAAATCCTCATCAAATTCTGAGGCACACTTTATATCATTC |
| B2_Ptep_Irx3_multiplexed | GGGGTTGTCCATCATGCACGATCCTAAATCATCCAGTAAACCGCC |
| B2_Ptep_Irx3_multiplexed | CCTCGTAAATCCTCATCAAAGGCATGAAATCTGTAAAAGAGGACA |
| B2_Ptep_Irx3_multiplexed | TTTTTATTGCAACACAACTCACTGTAAATCATCCAGTAAACCGCC |
| B2_Ptep_Irx3_multiplexed | CCTCGTAAATCCTCATCAAAGGACATTTTCTAAACAGAAATCCAT |
| B2_Ptep_Irx3_multiplexed | TCGAGTTCACGGAACACTCTGCTTTAAATCATCCAGTAAACCGCC |
| B2_Ptep_Irx3_multiplexed | CCTCGTAAATCCTCATCAAACAAGCGACCAAATTTTTGGTCTGCT |
| B2_Ptep_Irx3_multiplexed | GGGGACTATTGGATGTGGCAGTTCTAAATCATCCAGTAAACCGCC |
| B2_Ptep_Irx3_multiplexed | CCTCGTAAATCCTCATCAAAGGAAACGTCTTTTGCACTAGGTTTT |
| B2_Ptep_Irx3_multiplexed | GGAATCGAAGGTTTCCGAAAACCTAAAATCATCCAGTAAACCGCC |
| B2_Ptep_Irx3_multiplexed | CCTCGTAAATCCTCATCAAATCCTAACTGGGCTAGAATGTTCTGA |
| B2_Ptep_Irx3_multiplexed | CCTGCTGGAATCCCGGGTAGCAGGAAAATCATCCAGTAAACCGCC |
| B2_Ptep_Irx3_multiplexed | CCTCGTAAATCCTCATCAAAAGTTCGATAACATTCCAGAGCACAT |
| B2_Ptep_Irx3_multiplexed | GATGGAATCTTCTGATGAAAGACTTAAATCATCCAGTAAACCGCC |
| B2_Ptep_Irx3_multiplexed | CCTCGTAAATCCTCATCAAAAAGGTTCTTTGCTGTGGAATGTTTC |
| B2_Ptep_Irx3_multiplexed | GGGGAGATGTGGAATAATCGGCAGGAAATCATCCAGTAAACCGCC |
| B2_Ptep_Irx3_multiplexed | CCTCGTAAATCCTCATCAAACTTCTGCTGTATCGTCTGACTTCAG |
| B2_Ptep_Irx3_multiplexed | AAAAATCGTTCAAGTTGCTGCACTCAAATCATCCAGTAAACCGCC |
| B2_Ptep_Irx3_multiplexed | CCTCGTAAATCCTCATCAAACTGTCTGTATAGGTTGTACAGGAGG |
| B2_Ptep_Irx3_multiplexed | TTTTTAACACTCTCCATAGAACCGTAAATCATCCAGTAAACCGCC |
| B2_Ptep_Irx3_multiplexed | CCTCGTAAATCCTCATCAAATCCATGAGTTTCGTGATTCTTTCAA |
| B2_Ptep_Irx3_multiplexed | TAGGATCGTATGGGTAACAGGATGCAAATCATCCAGTAAACCGCC |
| B2_Ptep_Irx3_multiplexed | CCTCGTAAATCCTCATCAAAAGTGGTTGTGTTGGCTGCCAGTGAT |
| B2_Ptep_Irx3_multiplexed | GTAAGGAGCACCCAGTGTTGAGTAGAAATCATCCAGTAAACCGCC |
| B2_Ptep_Irx3_multiplexed | CCTCGTAAATCCTCATCAAAAGACCGAAGGATGCCAGCCGAGGTA |
| B2_Ptep_Irx3_multiplexed | ATGTGATCAGCGTATGAAGAACCGTAAATCATCCAGTAAACCGCC |
| B2_Ptep_Irx3_multiplexed | CCTCGTAAATCCTCATCAAATACCTGTGTTGAAGAAGGTGGATAT |
| B2_Ptep_Irx3_multiplexed | GCAACACGGTCCGCCGCTGCTCACCAAATCATCCAGTAAACCGCC |
| B2_Ptep_Irx3_multiplexed | CCTCGTAAATCCTCATCAAACGATTTGGACGGATTAGTGGGGTGT |
| B2_Ptep_Irx3_multiplexed | TAACCGAATTGTGAATAGGACATTAAAATCATCCAGTAAACCGCC |
| B2_Ptep_Irx3_multiplexed | CCTCGTAAATCCTCATCAAATCATTCGGCCTTTCTTGAACCGATC |
| B2_Ptep_Irx3_multiplexed | GAAAGGGAGATCGGCCTCACAGTGTAAATCATCCAGTAAACCGCC |
| B2_Ptep_Irx3_multiplexed | CCTCGTAAATCCTCATCAAAATTCCGTTATTCCAACTGTATCCCT |
| B2_Ptep_Irx3_multiplexed | CAAAAACATGGCGATCATCGGTCAAAAATCATCCAGTAAACCGCC |
| B3_Ptep_waist-less_multiplexed | GTCCCTGCCTCTATATCTTTATCTTCTAATTTCGTCCTATTGGGT |
| B3_Ptep_waist-less_multiplexed | TTAGCTCTCACACCGACTTGGAATATTCCACTCAACTTTAACCCG |
| B3_Ptep_waist-less_multiplexed | GTCCCTGCCTCTATATCTTTGCTGGAGCACTTGTAAAACCAACGG |
| B3_Ptep_waist-less_multiplexed | GATGTAGAAACTGTATGATACATAGTTCCACTCAACTTTAACCCG |
| B3_Ptep_waist-less_multiplexed | GTCCCTGCCTCTATATCTTTCTGGCTGAAATGTCCTTATGTCTAG |
| B3_Ptep_waist-less_multiplexed | ATGAAAAAGAGGATGTCCTCAAGAATTCCACTCAACTTTAACCCG |
| B3_Ptep_waist-less_multiplexed | GTCCCTGCCTCTATATCTTTATAGTCTATAGGATTATGCCTCAGA |
| B3_Ptep_waist-less_multiplexed | ATGTCCCATTTGGGCATAAAATTGTTTCCACTCAACTTTAACCCG |
| B3_Ptep_waist-less_multiplexed | GTCCCTGCCTCTATATCTTTCTACCATTCAGAAGGCGATTGTAAT |
| B3_Ptep_waist-less_multiplexed | TGTGAACTGTCACCACAAGGTAAAATTCCACTCAACTTTAACCCG |
| B3_Ptep_waist-less_multiplexed | GTCCCTGCCTCTATATCTTTAGGAGGATGGTGACACAGAAGCTAA |
| B3_Ptep_waist-less_multiplexed | TGGGTGGTAAAACGGATGGGGATCCTTCCACTCAACTTTAACCCG |
| B3_Ptep_waist-less_multiplexed | GTCCCTGCCTCTATATCTTTGAATTTGTCACCATGTGGGAAATTG |
| B3_Ptep_waist-less_multiplexed | ACTTCGCTCGTGGGAGGTGACACTATTCCACTCAACTTTAACCCG |
| B3_Ptep_waist-less_multiplexed | GTCCCTGCCTCTATATCTTTGACGATAAAGATGATTCGGCTGAAC |
| B3_Ptep_waist-less_multiplexed | AACTCTCCTGTTCTCCAAAGGGGTGTTCCACTCAACTTTAACCCG |
| B3_Ptep_waist-less_multiplexed | GTCCCTGCCTCTATATCTTTGTGAAGGATGGTATAAGTGTTCATT |
| B3_Ptep_waist-less_multiplexed | CCTGCGGAGGAAGAGGCCTGTGGTATTCCACTCAACTTTAACCCG |
| B3_Ptep_waist-less_multiplexed | GTCCCTGCCTCTATATCTTTATCAGTTTGGTGGTCGAAGAACTTT |
| B3_Ptep_waist-less_multiplexed | ACCCAACCTTCTTAATTTTTGGTTCTTCCACTCAACTTTAACCCG |
| B3_Ptep_waist-less_multiplexed | GTCCCTGCCTCTATATCTTTGCGCTAGTGACCAGATTTTTGGTTT |
| B3_Ptep_waist-less_multiplexed | GTGGTGGGCTCTCTGAGGTGGCAGTTTCCACTCAACTTTAACCCG |
| B3_Ptep_waist-less_multiplexed | GTCCCTGCCTCTATATCTTTAAGTTCAGGGAATTTGGTTTCTGAG |
| B3_Ptep_waist-less_multiplexed | AATTGATTCTGATTTGATGTGTTCATTCCACTCAACTTTAACCCG |
| B3_Ptep_waist-less_multiplexed | GTCCCTGCCTCTATATCTTTTCGGGAGCACTGAGGCTGTCACCAA |
| B3_Ptep_waist-less_multiplexed | TCAAGCGCTTTCATTGGAGACCTGTTTCCACTCAACTTTAACCCG |
| B3_Ptep_waist-less_multiplexed | GTCCCTGCCTCTATATCTTTCATCGTCCTTCATACTGTCTCGTCT |
| B3_Ptep_waist-less_multiplexed | GGGAGGTGCATTTCTGTTGTCGGTTTTCCACTCAACTTTAACCCG |
| B3_Ptep_waist-less_multiplexed | GTCCCTGCCTCTATATCTTTTTGATTTTCATTCGAATCGCTGAGT |
| B3_Ptep_waist-less_multiplexed | TCTACTGGAGTCACTTTCCTGGCATTTCCACTCAACTTTAACCCG |
| B3_Ptep_waist-less_multiplexed | GTCCCTGCCTCTATATCTTTTCATCTTTATCATCATCTTCATCCA |
| B3_Ptep_waist-less_multiplexed | CTCTTAAGGTCATCCCCACTCTGCTTTCCACTCAACTTTAACCCG |
| B3_Ptep_waist-less_multiplexed | GTCCCTGCCTCTATATCTTTATCTCCGTACGGAGAATATGCGGCG |
| B3_Ptep_waist-less_multiplexed | TTTTTCACTATCAACAGGATCAAACTTCCACTCAACTTTAACCCG |
| B3_Ptep_waist-less_multiplexed | GTCCCTGCCTCTATATCTTTAGCCCGAGGATCTTTTATGTCATAA |
| B3_Ptep_waist-less_multiplexed | GTGTCCTGGTGGTAGGGATGCCCAGTTCCACTCAACTTTAACCCG |
| B3_Ptep_waist-less_multiplexed | GTCCCTGCCTCTATATCTTTGTGGCATGCTGTTTGAGGCTGGATA |
| B3_Ptep_waist-less_multiplexed | GATGGGCCTGACTTACTAGTGTGTCTTCCACTCAACTTTAACCCG |
| B3_Ptep_waist-less_multiplexed | GTCCCTGCCTCTATATCTTTTTGAGGAGATGCAGTATAGGCGAAA |
| B3_Ptep_waist-less_multiplexed | GCTGGGGTGCTGGCCTGTCATCAGATTCCACTCAACTTTAACCCG |
| **Initiator and gene name** | **Sequence** |
| B1_Ptep_pnr1_multiplexed | GAGGAGGGCAGCAAACGGAATCTAGATTCCGACGTTGCAGAACTG |
| B1_Ptep_pnr1_multiplexed | TTATCTGTTCATATCGTTATAACAATAGAAGAGTCTTCCTTTACG |
| B1_Ptep_pnr1_multiplexed | GAGGAGGGCAGCAAACGGAATTTGCGTAACTTCCATGTTGCATTT |
| B1_Ptep_pnr1_multiplexed | GAATGTTGGTAGGAACTGTCTACGCTAGAAGAGTCTTCCTTTACG |
| B1_Ptep_pnr1_multiplexed | GAGGAGGGCAGCAAACGGAAATATCGTTTTGTGATGCGATGAGGA |
| B1_Ptep_pnr1_multiplexed | TTGCTGTGTGCTGCGTGCACGATGATAGAAGAGTCTTCCTTTACG |
| B1_Ptep_pnr1_multiplexed | GAGGAGGGCAGCAAACGGAATCCTGCCACTTGTGTACCCGAAGTT |
| B1_Ptep_pnr1_multiplexed | TTGTGGTAAGACAGAATGACAACCGTAGAAGAGTCTTCCTTTACG |
| B1_Ptep_pnr1_multiplexed | GAGGAGGGCAGCAAACGGAAATGGTGATTGCAAGGATGGCGTTGT |
| B1_Ptep_pnr1_multiplexed | GCGGAGAGCTGTGTATGTGGCTTGTTAGAAGAGTCTTCCTTTACG |
| B1_Ptep_pnr1_multiplexed | GAGGAGGGCAGCAAACGGAAGAAGACTGTTCTGGTTTCACAGTTG |
| B1_Ptep_pnr1_multiplexed | TTCTGAAGATCAATAATATCAGAGGTAGAAGAGTCTTCCTTTACG |
| B1_Ptep_pnr1_multiplexed | GAGGAGGGCAGCAAACGGAATTTTACGAGTCTGTATTCCAGCTTT |
| B1_Ptep_pnr1_multiplexed | CAGTTTTGTTAGTGTTCTTTGGTTTTAGAAGAGTCTTCCTTTACG |
| B1_Ptep_pnr1_multiplexed | GAGGAGGGCAGCAAACGGAAGCTGGCAGCCGTCTTTGTGGTTTAC |
| B1_Ptep_pnr1_multiplexed | TTTTTGGAGAGTCCCATCCTACGAGTAGAAGAGTCTTCCTTTACG |
| B1_Ptep_pnr1_multiplexed | GAGGAGGGCAGCAAACGGAAGTCAAATTCTTGTCTTCTAGATAGG |
| B1_Ptep_pnr1_multiplexed | TTTATTGAATGGGTGATAATCTCCTTAGAAGAGTCTTCCTTTACG |
| B1_Ptep_pnr1_multiplexed | GAGGAGGGCAGCAAACGGAATTTTCTAGGCTTCCCCAATGCCCAT |
| B1_Ptep_pnr1_multiplexed | CTCTGCGAAGATTGTAGGGCTGCCCTAGAAGAGTCTTCCTTTACG |
| B1_Ptep_pnr1_multiplexed | GAGGAGGGCAGCAAACGGAACATTGCTACGCTGGAAGTAGGACGT |
| B1_Ptep_pnr1_multiplexed | CCATGTATGAATGATATGCACTTAATAGAAGAGTCTTCCTTTACG |
| B1_Ptep_pnr1_multiplexed | GAGGAGGGCAGCAAACGGAATGACAAAGATGGAGCCTGAAATCCA |
| B1_Ptep_pnr1_multiplexed | ATCTCGGTTCACACCGCCAATGGAGTAGAAGAGTCTTCCTTTACG |
| B1_Ptep_pnr1_multiplexed | GAGGAGGGCAGCAAACGGAATATGAGTCAGAAAGGGGAACTCCAG |
| B1_Ptep_pnr1_multiplexed | CGAACTGCAGACAATTGGGCATGATTAGAAGAGTCTTCCTTTACG |
| B1_Ptep_pnr1_multiplexed | GAGGAGGGCAGCAAACGGAAATGTCCAGGTATTAGGATTGCTTGT |
| B1_Ptep_pnr1_multiplexed | TCCGACCACCATGATGATCACCAGATAGAAGAGTCTTCCTTTACG |
| B1_Ptep_pnr1_multiplexed | GAGGAGGGCAGCAAACGGAACACCATTGGTCGCGAATTTGGTATG |
| B1_Ptep_pnr1_multiplexed | TGGTAACTGACCACCTTGATGAAATTAGAAGAGTCTTCCTTTACG |
| B1_Ptep_pnr1_multiplexed | GAGGAGGGCAGCAAACGGAATCTTGGTACGCGGGCATATGAATGG |
| B1_Ptep_pnr1_multiplexed | ATTGAACTGGGATGGTGCAAGTAACTAGAAGAGTCTTCCTTTACG |
| B1_Ptep_pnr1_multiplexed | GAGGAGGGCAGCAAACGGAATTGCCCACTCTGTTTTAGGATAGTT |
| B1_Ptep_pnr1_multiplexed | ACATGTTCTGGAACATAGTGGTTCCTAGAAGAGTCTTCCTTTACG |
| B1_Ptep_pnr1_multiplexed | GAGGAGGGCAGCAAACGGAATTGCACTTTTGTTGAAGAGAAGAAC |
| B1_Ptep_pnr1_multiplexed | ACGTTTCGTGAAATGGCGAAACTTATAGAAGAGTCTTCCTTTACG |
| B1_Ptep_pnr1_multiplexed | GAGGAGGGCAGCAAACGGAAAACCATATAGACGAATTCAAGCAGC |
| B1_Ptep_pnr1_multiplexed | GGCGTTAATATTCCATAAGCGTTTTTAGAAGAGTCTTCCTTTACG |
| B1_Ptep_pnr1_multiplexed | GAGGAGGGCAGCAAACGGAATATTTCTTTGACACAGCGTTCGACC |
| B1_Ptep_pnr1_multiplexed | CTACAATTAGAAAATAGGTGACATATAGAAGAGTCTTCCTTTACG |
| B2_Ptep_pnr2_multiplexed | CCTCGTAAATCCTCATCAAAGTATGGCGCGGGCATTCTATTTGAC |
| B2_Ptep_pnr2_multiplexed | TTAACCATCACGAATCTCATGTTTGAAATCATCCAGTAAACCGCC |
| B2_Ptep_pnr2_multiplexed | CCTCGTAAATCCTCATCAAAGTCCATGCTGTGATGACTGTTATAG |
| B2_Ptep_pnr2_multiplexed | TTGGTGGTGATGGCTTGAACTGGAGAAATCATCCAGTAAACCGCC |
| B2_Ptep_pnr2_multiplexed | CCTCGTAAATCCTCATCAAAAGCAGGGGTTGAATTGTTGCTTACT |
| B2_Ptep_pnr2_multiplexed | AGTTGCATCGCAAGATGCTGGAGAAAAATCATCCAGTAAACCGCC |
| B2_Ptep_pnr2_multiplexed | CCTCGTAAATCCTCATCAAATGGTGGTCTGTTTACCCCATGTAAT |
| B2_Ptep_pnr2_multiplexed | TTTTTGAATTCCTTCCTTTTTCATTAAATCATCCAGTAAACCGCC |
| B2_Ptep_pnr2_multiplexed | CCTCGTAAATCCTCATCAAACAGATATTGCCGGATCTAGCAATCC |
| B2_Ptep_pnr2_multiplexed | CATCAGAACAATATGGATGTTGCAAAAATCATCCAGTAAACCGCC |
| B2_Ptep_pnr2_multiplexed | CCTCGTAAATCCTCATCAAAGTCAAAGCTGAAATTCCATAACCAG |
| B2_Ptep_pnr2_multiplexed | GAATTTTGAGCAAGATTCAGTCCAGAAATCATCCAGTAAACCGCC |
| B2_Ptep_pnr2_multiplexed | CCTCGTAAATCCTCATCAAAAAGCAGCGCTGGAGGGAGATGATGG |
| B2_Ptep_pnr2_multiplexed | CGAGACTTCCAGTTGACGAAGCATAAAATCATCCAGTAAACCGCC |
| B2_Ptep_pnr2_multiplexed | CCTCGTAAATCCTCATCAAAATAGTTGGAGCACGTTCCCTGTTTC |
| B2_Ptep_pnr2_multiplexed | ACCATGACTGTATCACTGGAACTGCAAATCATCCAGTAAACCGCC |
| B2_Ptep_pnr2_multiplexed | CCTCGTAAATCCTCATCAAAGTCGACTGTTCTCTATGATTGATGG |
| B2_Ptep_pnr2_multiplexed | CGACTATTAGATGTTTCATTGGCTGAAATCATCCAGTAAACCGCC |
| B2_Ptep_pnr2_multiplexed | CCTCGTAAATCCTCATCAAATCATTGCGCAAGAAGAAGTATTCTC |
| B2_Ptep_pnr2_multiplexed | TTGCTAAGCTGTTGTTGTTATTCGAAAATCATCCAGTAAACCGCC |
| B2_Ptep_pnr2_multiplexed | CCTCGTAAATCCTCATCAAAAGCATGTTCTGTAACATTTGCATGG |
| B2_Ptep_pnr2_multiplexed | GTTACAATGATATGAATCTGAGACGAAATCATCCAGTAAACCGCC |
| B2_Ptep_pnr2_multiplexed | CCTCGTAAATCCTCATCAAATGCTTTGATGGTCAGTTTTGGACAG |
| B2_Ptep_pnr2_multiplexed | TAGCATGGTGGTGTGAATCCCATTCAAATCATCCAGTAAACCGCC |
| B2_Ptep_pnr2_multiplexed | CCTCGTAAATCCTCATCAAACAGTTCCATGTCCATCGTTTCTATT |
| B2_Ptep_pnr2_multiplexed | TTTTTGCCATTTGAGTAGTTTCTTGAAATCATCCAGTAAACCGCC |
| B2_Ptep_pnr2_multiplexed | CCTCGTAAATCCTCATCAAATAATCTTCAACAGCGAGAACATTTG |
| B2_Ptep_pnr2_multiplexed | TTTTTGACTGCTGGAGGAACTAAACAAATCATCCAGTAAACCGCC |
| B2_Ptep_pnr2_multiplexed | CCTCGTAAATCCTCATCAAAATTACTGTCATCTCTTGATACGGTT |
| B2_Ptep_pnr2_multiplexed | TTGTACTGCTTTATTCACTCCCGCGAAATCATCCAGTAAACCGCC |
| B2_Ptep_pnr2_multiplexed | CCTCGTAAATCCTCATCAAAAAGATTCTTTGGGAAGAACGGGCTG |
| B2_Ptep_pnr2_multiplexed | TACCAAAATCATTTACATTACTGTCAAATCATCCAGTAAACCGCC |
| B2_Ptep_pnr2_multiplexed | CCTCGTAAATCCTCATCAAAAATGTGCTGGCTCCAGCACTAAATA |
| B2_Ptep_pnr2_multiplexed | GTATAGAATTTTTGGTCATAGAATTAAATCATCCAGTAAACCGCC |
| B2_Ptep_pnr2_multiplexed | CCTCGTAAATCCTCATCAAAAAATTATGGGCAGAAGTAAATACGG |
| B2_Ptep_pnr2_multiplexed | TGTGCTTCATGATTGTTAACTAACGAAATCATCCAGTAAACCGCC |
| B2_Ptep_pnr2_multiplexed | CCTCGTAAATCCTCATCAAACTAAAGTTTCTGGCTTCATATTGCG |
| B2_Ptep_pnr2_multiplexed | CAAAGTTCTTATAAGACAAGTTGCGAAATCATCCAGTAAACCGCC |
| B2_Ptep_pnr2_multiplexed | CCTCGTAAATCCTCATCAAACTTTCGACGATTTGAGGATAGGATT |
| B2_Ptep_pnr2_multiplexed | ATGCTGTTTAAATTCTTCGAAGCTTAAATCATCCAGTAAACCGCC |
| B3_Ptep_pnr3_multiplexed | GTCCCTGCCTCTATATCTTTGTTTCAAGAACCATTCCATTTTCTA |
| B3_Ptep_pnr3_multiplexed | GGGGCCAAACTGATAACAGAGGGGTTTCCACTCAACTTTAACCCG |
| B3_Ptep_pnr3_multiplexed | GTCCCTGCCTCTATATCTTTGAAAATGGGGCAGCTGTTGAGATAA |
| B3_Ptep_pnr3_multiplexed | CCGTTTCGTTTGGGGAGTCCAAGGGTTCCACTCAACTTTAACCCG |
| B3_Ptep_pnr3_multiplexed | GTCCCTGCCTCTATATCTTTAGATGAGGTAGAGCTAGTGGTTGAA |
| B3_Ptep_pnr3_multiplexed | CCCCTCCATGCTCACCTGTGGAAGATTCCACTCAACTTTAACCCG |
| B3_Ptep_pnr3_multiplexed | GTCCCTGCCTCTATATCTTTCTGGAGCCAATCCCTGATGAATGTC |
| B3_Ptep_pnr3_multiplexed | TTCGGAGATTTGCTGGAAGATCCATTTCCACTCAACTTTAACCCG |
| B3_Ptep_pnr3_multiplexed | GTCCCTGCCTCTATATCTTTTAGGCTTTCTTTTTCTTGTCTGAAT |
| B3_Ptep_pnr3_multiplexed | TTCCCTTGGATGAAGATCCACTGCTTTCCACTCAACTTTAACCCG |
| B3_Ptep_pnr3_multiplexed | GTCCCTGCCTCTATATCTTTCACTTGGTGTAATTTGTAATACAAT |
| B3_Ptep_pnr3_multiplexed | ATCCTTTCTCATTGAGATGGGCCGATTCCACTCAACTTTAACCCG |
| B3_Ptep_pnr3_multiplexed | GTCCCTGCCTCTATATCTTTGTGGTCAGCCTTCTTTGAGGCCTTA |
| B3_Ptep_pnr3_multiplexed | TTTTTGGTCATACCAACTCTACGACTTCCACTCAACTTTAACCCG |
| B3_Ptep_pnr3_multiplexed | GTCCCTGCCTCTATATCTTTCTGGTGGTGGATCACTATATTGAGA |
| B3_Ptep_pnr3_multiplexed | TTTTTGCTTCACTGATTCTACCATATTCCACTCAACTTTAACCCG |
| B3_Ptep_pnr3_multiplexed | GTCCCTGCCTCTATATCTTTAGTCATAGAAGGTAAAGTCGACCAA |
| B3_Ptep_pnr3_multiplexed | TCTGCCATCTTGTAGGTACGAAGACTTCCACTCAACTTTAACCCG |
| B3_Ptep_pnr3_multiplexed | GTCCCTGCCTCTATATCTTTGAGCTTACGTAACTGCTATAACTGC |
| B3_Ptep_pnr3_multiplexed | TTCTCTGCTGACGTCAGTAAGTAAGTTCCACTCAACTTTAACCCG |
| B3_Ptep_pnr3_multiplexed | GTCCCTGCCTCTATATCTTTCTAAAATGGGGCTTCCCAAGCCAGA |
| B3_Ptep_pnr3_multiplexed | TTCCATTAGCGCTCACTTCATTATTTTCCACTCAACTTTAACCCG |
| B3_Ptep_pnr3_multiplexed | GTCCCTGCCTCTATATCTTTGCCCCACATGGGAAGAGTGCTAGCA |
| B3_Ptep_pnr3_multiplexed | AGTGCCTGCGTTGTTATTGTTAAGATTCCACTCAACTTTAACCCG |
| B3_Ptep_pnr3_multiplexed | GTCCCTGCCTCTATATCTTTATTGGGCTGTATACAACACTTTGAG |
| B3_Ptep_pnr3_multiplexed | TTCACCTCTTGTGAAGGATGTTGAATTCCACTCAACTTTAACCCG |
| B3_Ptep_pnr3_multiplexed | GTCCCTGCCTCTATATCTTTAGCATGGACCATTTGTACATTTTCG |
| B3_Ptep_pnr3_multiplexed | TATAGCTCTGGGATTATCACTTTCATTCCACTCAACTTTAACCCG |
| B3_Ptep_pnr3_multiplexed | GTCCCTGCCTCTATATCTTTGGAGATGTTGATCCACCAGGAGTAG |
| B3_Ptep_pnr3_multiplexed | CAGGAAGCACTTTTCAAATAACTATTTCCACTCAACTTTAACCCG |
| B3_Ptep_pnr3_multiplexed | GTCCCTGCCTCTATATCTTTCATAACTGACGCCATACTGTCCCAT |
| B3_Ptep_pnr3_multiplexed | AAGTCGGTGTCGGACTATAACTTCCTTCCACTCAACTTTAACCCG |
| B3_Ptep_pnr3_multiplexed | GTCCCTGCCTCTATATCTTTTACTAATTGTTCTCCAGCTTGTGTG |
| B3_Ptep_pnr3_multiplexed | GCCATTATTGTCTCCTAGAGTTTGATTCCACTCAACTTTAACCCG |
| B3_Ptep_pnr3_multiplexed | GTCCCTGCCTCTATATCTTTAATTTGGATAGCATCGGCCGTGTGA |
| B3_Ptep_pnr3_multiplexed | CGATGACTCCACGGCAGCTGAATAGTTCCACTCAACTTTAACCCG |
| B3_Ptep_pnr3_multiplexed | GTCCCTGCCTCTATATCTTTTGTAGAATCGCAGTGTTGTGAGTTT |
| B3_Ptep_pnr3_multiplexed | CCGGCCCGATGTCGGGGAAGCATCTTTCCACTCAACTTTAACCCG |
| B3_Ptep_pnr3_multiplexed | GTCCCTGCCTCTATATCTTTTGCTGCTGGGGTGGGGTCGGTAATG |
| B3_Ptep_pnr3_multiplexed | CTCAGATCTTCTGTAGATCGGGAATTTCCACTCAACTTTAACCCG |
| **Initiator and gene name** | **Sequence** |
| B2_Popi_irx2_multiplexed | CCTCGTAAATCCTCATCAAATCTTCGGAGAGGGAAAGCCTCTGAT |
| B2_Popi_irx2_multiplexed | GAAGCGCACGGAGACGAAGATTCGTAAATCATCCAGTAAACCGCC |
| B2_Popi_irx2_multiplexed | CCTCGTAAATCCTCATCAAATCTAACGTTCTTGACATCGTGAAGG |
| B2_Popi_irx2_multiplexed | CCCCGGTACTTCGCACATCATCGATAAATCATCCAGTAAACCGCC |
| B2_Popi_irx2_multiplexed | CCTCGTAAATCCTCATCAAAGTAATTGTTCAACTGATGTCCGACT |
| B2_Popi_irx2_multiplexed | ATGGTCACTCGGAGCCCCTACACCAAAATCATCCAGTAAACCGCC |
| B2_Popi_irx2_multiplexed | CCTCGTAAATCCTCATCAAACAATGTTTCTGATACGGTTGGTTCG |
| B2_Popi_irx2_multiplexed | GGGTAGTTGAAAGTTTCTCTGTTCAAAATCATCCAGTAAACCGCC |
| B2_Popi_irx2_multiplexed | CCTCGTAAATCCTCATCAAAAGCGGCTAAAGACCATATTTTTGGC |
| B2_Popi_irx2_multiplexed | GGGTGGAGGACTCTTACTGGTAGCCAAATCATCCAGTAAACCGCC |
| B2_Popi_irx2_multiplexed | CCTCGTAAATCCTCATCAAAACGCGTCTTTAATGAAAGCGTCCAA |
| B2_Popi_irx2_multiplexed | TCTCGCCCGACGAGGTGGATGACAGAAATCATCCAGTAAACCGCC |
| B2_Popi_irx2_multiplexed | CCTCGTAAATCCTCATCAAATTTGTTTGTTCCGATGATACTTGGT |
| B2_Popi_irx2_multiplexed | GTAAAGGCCGAGGTGCCCGTTAAAAAAATCATCCAGTAAACCGCC |
| B2_Popi_irx2_multiplexed | CCTCGTAAATCCTCATCAAAGATGAAGCGGAAGGTACCGATGGTA |
| B2_Popi_irx2_multiplexed | GTCGAATGATGAGGCCGAAATGGTCAAATCATCCAGTAAACCGCC |
| B2_Popi_irx2_multiplexed | CCTCGTAAATCCTCATCAAACAAACGAGATCCATTTTGAGACTCG |
| B2_Popi_irx2_multiplexed | ATAGCCGCTGTGCAATCCCAACGTTAAATCATCCAGTAAACCGCC |
| B2_Popi_irx2_multiplexed | CCTCGTAAATCCTCATCAAATTCGATCATCGTATTCGTCATTCCT |
| B2_Popi_irx2_multiplexed | GTGTCCTGCCAAGACACGCTAACCTAAATCATCCAGTAAACCGCC |
| B2_Popi_irx2_multiplexed | CCTCGTAAATCCTCATCAAAAAAATCACCTCCTTGGTCTTTCGCT |
| B2_Popi_irx2_multiplexed | TTGTCCAATCTTTCTTGCTTTGTTGAAATCATCCAGTAAACCGCC |
| B2_Popi_irx2_multiplexed | CCTCGTAAATCCTCATCAAACAGTTTGACTATCATCTCCGTGGTC |
| B2_Popi_irx2_multiplexed | CATCTTCGGATCTTCTACTACCAGAAAATCATCCAGTAAACCGCC |
| B2_Popi_irx2_multiplexed | CCTCGTAAATCCTCATCAAAAACCTCCGTACCCATACGCTGCCAA |
| B2_Popi_irx2_multiplexed | TTTTTCGTCTTGCCCCATTAAAATCAAATCATCCAGTAAACCGCC |
| B2_Popi_irx2_multiplexed | CCTCGTAAATCCTCATCAAATTGATGAGCATTCGCCGGATGAGCA |
| B2_Popi_irx2_multiplexed | TGGGTCGTAGTAATACGAAGCTGTAAAATCATCCAGTAAACCGCC |
| B2_Popi_irx2_multiplexed | CCTCGTAAATCCTCATCAAATCTTTCATTCCAAATCCAGTACTCA |
| B2_Popi_irx2_multiplexed | GAAGACAACGTTCCCCTCCACGGATAAATCATCCAGTAAACCGCC |
| B2_Popi_irx2_multiplexed | CCTCGTAAATCCTCATCAAATAGCCAAACTAGCGTATCCGTTACT |
| B2_Popi_irx2_multiplexed | TTGGATAAAAGGCGGAAGAATCAACAAATCATCCAGTAAACCGCC |
| B2_Popi_irx2_multiplexed | CCTCGTAAATCCTCATCAAAGTGGATGAACTTGATGCGAGTGATT |
| B2_Popi_irx2_multiplexed | TCCCGTAGAATCCGGCGGCCGTCAAAAATCATCCAGTAAACCGCC |
| B2_Popi_irx2_multiplexed | CCTCGTAAATCCTCATCAAACGCCGTGGATATCAACCTGGATTCG |
| B2_Popi_irx2_multiplexed | GGAATGGATAGAAGGGTGGTGATACAAATCATCCAGTAAACCGCC |
| B2_Popi_irx2_multiplexed | CCTCGTAAATCCTCATCAAAGGCGATTGGGGTGAAGGTGAAGAAA |
| B2_Popi_irx2_multiplexed | GAAGACACGGTGGCTTCAACGCAAGAAATCATCCAGTAAACCGCC |
| B2_Popi_irx2_multiplexed | CCTCGTAAATCCTCATCAAAACCACCGGTTGATACGGTCAACATT |
| B2_Popi_irx2_multiplexed | CGTGCAACTGGCACCACCAGTAACAAAATCATCCAGTAAACCGCC |
| B3_popi_irx3_multiplexed | GTCCCTGCCTCTATATCTTTATCAGTTCTTAGATGGTCCAAGGCG |
| B3_popi_irx3_multiplexed | TTATTGGACTTGAGCCGAACCGTCGTTCCACTCAACTTTAACCCG |
| B3_popi_irx3_multiplexed | GTCCCTGCCTCTATATCTTTGTCGATCGTGAGTTTGAGATTCGCG |
| B3_popi_irx3_multiplexed | GTGATGTTCTGACCAAAGGAGGCGGTTCCACTCAACTTTAACCCG |
| B3_popi_irx3_multiplexed | GTCCCTGCCTCTATATCTTTCATGCCATGAGGCGAGGTAATATCA |
| B3_popi_irx3_multiplexed | AAGATCGTAAGCCATGGCTGACGAGTTCCACTCAACTTTAACCCG |
| B3_popi_irx3_multiplexed | GTCCCTGCCTCTATATCTTTTGCGACACTCGACGATGAAATAATG |
| B3_popi_irx3_multiplexed | TAAGCCCTAGACGCCTCATTTCCCTTTCCACTCAACTTTAACCCG |
| B3_popi_irx3_multiplexed | GTCCCTGCCTCTATATCTTTCGGTCGTTGCTGGATCGAAAGCCGA |
| B3_popi_irx3_multiplexed | ATTGTTCCTTGGATTCGTCGTCGCCTTCCACTCAACTTTAACCCG |
| B3_popi_irx3_multiplexed | GTCCCTGCCTCTATATCTTTACCGCGTGTCCCATCGTTCCGTTCA |
| B3_popi_irx3_multiplexed | ACTGGTGACGTCGACGAACAGTTTTTTCCACTCAACTTTAACCCG |
| B3_popi_irx3_multiplexed | GTCCCTGCCTCTATATCTTTTGGCGTAATGGTCTTCGTAGTGTCC |
| B3_popi_irx3_multiplexed | AGTTTTTAGCACTCGTCACTCCGTTTTCCACTCAACTTTAACCCG |
| B3_popi_irx3_multiplexed | GTCCCTGCCTCTATATCTTTCATATCGCCGTCCATTTCCGACTTA |
| B3_popi_irx3_multiplexed | GTTGGCCACGTTGGCTCGTCCGCAATTCCACTCAACTTTAACCCG |
| B3_popi_irx3_multiplexed | GTCCCTGCCTCTATATCTTTTTGATACCGCTCGGTGTCGCGGGTA |
| B3_popi_irx3_multiplexed | TATTTAGGATCTCGATATTCCTGTCTTCCACTCAACTTTAACCCG |
| B3_popi_irx3_multiplexed | GTCCCTGCCTCTATATCTTTGCTAAGGACCATATTTTCGGTCTAT |
| B3_popi_irx3_multiplexed | GGCGGACTGTCCGAAGTGGCCGTGTTTCCACTCAACTTTAACCCG |
| B3_popi_irx3_multiplexed | GTCCCTGCCTCTATATCTTTGCAAACCACCTCTTAACAAGAGATT |
| B3_popi_irx3_multiplexed | ACGGAGGGGTGACGTGGTGACCGTGTTCCACTCAACTTTAACCCG |
| B3_popi_irx3_multiplexed | GTCCCTGCCTCTATATCTTTCAAGGGTGTCGACAACGGGTTGACG |
| B3_popi_irx3_multiplexed | TTGTTGCAAATGAAGGTTATGCGAATTCCACTCAACTTTAACCCG |
| B3_popi_irx3_multiplexed | GTCCCTGCCTCTATATCTTTCTACTCGTTGTGCCAATTCCCCTGT |
| B3_popi_irx3_multiplexed | TCCGCCGAACTTGAAGAGTTCTCCGTTCCACTCAACTTTAACCCG |
| B3_popi_irx3_multiplexed | GTCCCTGCCTCTATATCTTTAATCTCTCGACAACTCGTCTTCTTG |
| B3_popi_irx3_multiplexed | GATGATGGTGCATCTTGTAGTCGGATTCCACTCAACTTTAACCCG |
| B3_popi_irx3_multiplexed | GTCCCTGCCTCTATATCTTTGCCCGATCGCTCATCGTGAATGGAG |
| B3_popi_irx3_multiplexed | CATATGAGAACCCGTGTTCATTTTGTTCCACTCAACTTTAACCCG |
| B3_popi_irx3_multiplexed | GTCCCTGCCTCTATATCTTTACCGTCGACGCCGTCATTCGTCTTT |
| B3_popi_irx3_multiplexed | TTCATCGCCGCTGTCCGATTTGTCATTCCACTCAACTTTAACCCG |
| B3_popi_irx3_multiplexed | GTCCCTGCCTCTATATCTTTTTTCAAATCGTAGGGACTTGGCAAT |
| B3_popi_irx3_multiplexed | CAGGGATCCCCACGCTCCTCTACCGTTCCACTCAACTTTAACCCG |
| B3_popi_irx3_multiplexed | GTCCCTGCCTCTATATCTTTAGAGCGGCTGTTACGTAACTGGTTT |
| B3_popi_irx3_multiplexed | GAATAGAAGGCGGAGGCGTTGGTTCTTCCACTCAACTTTAACCCG |
| B3_popi_irx3_multiplexed | GTCCCTGCCTCTATATCTTTCTTCCACTGGTCGGGGTACTGTTAT |
| B3_popi_irx3_multiplexed | TCCGGAGACATGACACCGCTAGTAATTCCACTCAACTTTAACCCG |
| B3_popi_irx3_multiplexed | GTCCCTGCCTCTATATCTTTGTCCGGTCATAAGAAGCTGAGAGGA |
| B3_popi_irx3_multiplexed | AGCAAGTTGTGGCAACTGGAGGTTGTTCCACTCAACTTTAACCCG |

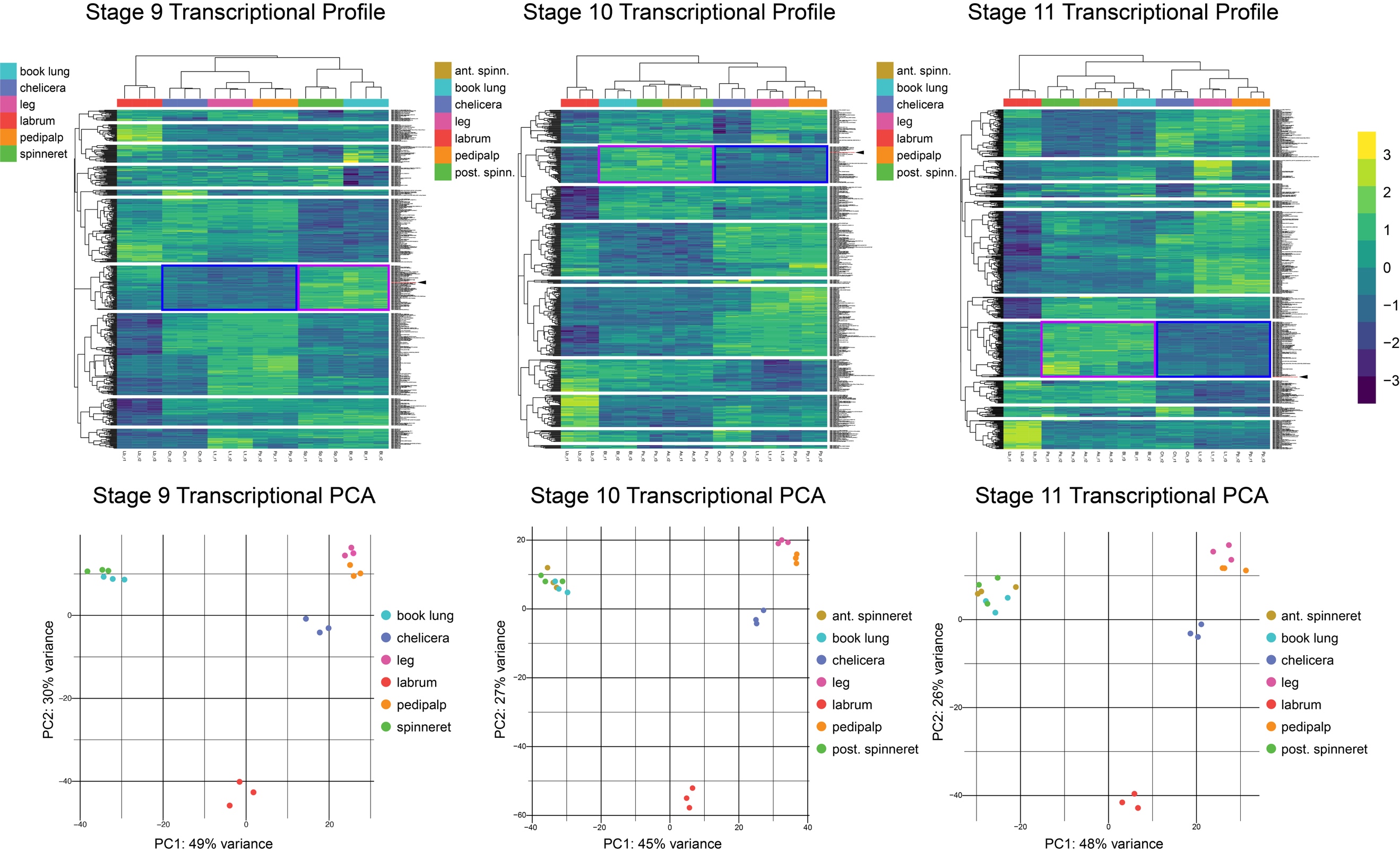

**Figure S1. Transcriptional profiles and principal components analysis of territory-specific gene expression in embryos of the tarantula *Aphonopelma hentzi*.** From left to right: stage 9, stage 10, and stage 11. In upper row, boxes highlight genes that are highly expressed in opisthosomal segments (purple) and lowly expressed in prosomal segments (blue). Arrowhead indicates tarantula ortholog of *waist-less*.

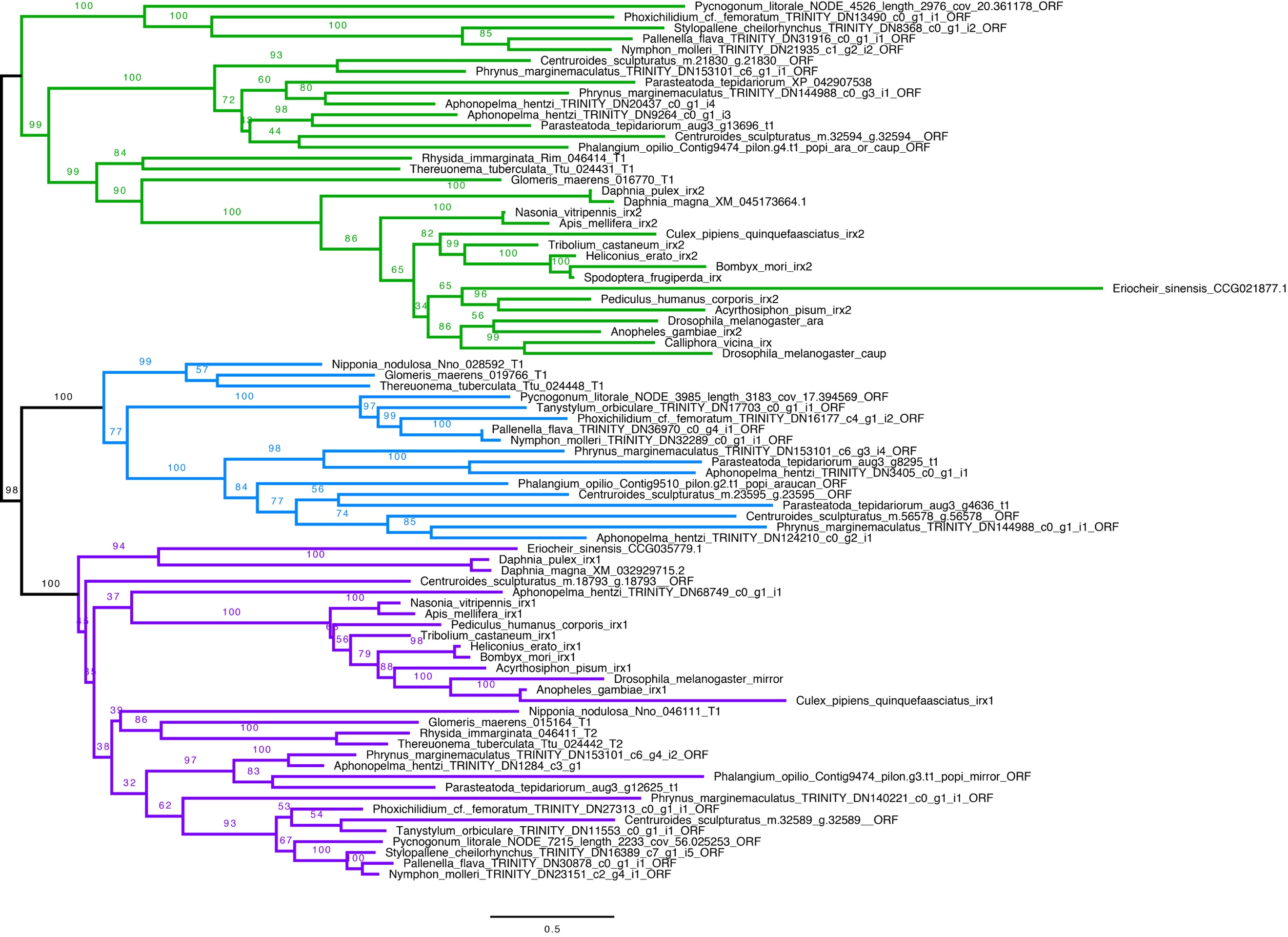

**Figure S2. Maximum likelihood tree topology of panarthropod Iroquois homologs.** Colors correspond to major phylogenetic lineages. Numbers on nodes correspond to bootstrap resampling frequencies.

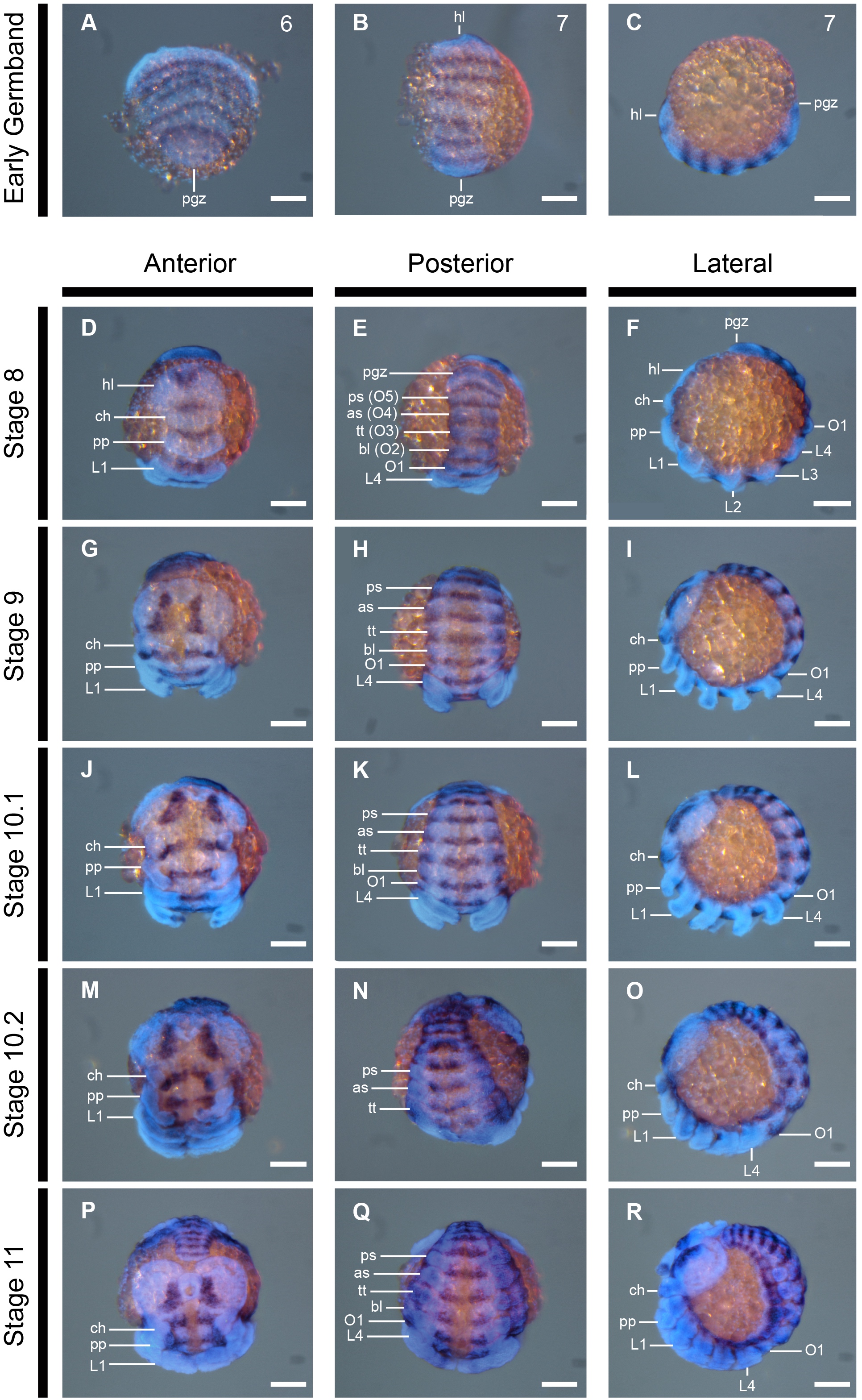

**Figure S3. Wild type expression of *waist-less* during embryogenesis of the cobweb spider *Parasteatoda tepidariorum*.** All panels constitute merged images of Hoechst and DIG-labeled *in situ* hybridization for *Ptep-waist-less*. Abbreviations as in Figure 2. Scale bar: 100 μm.

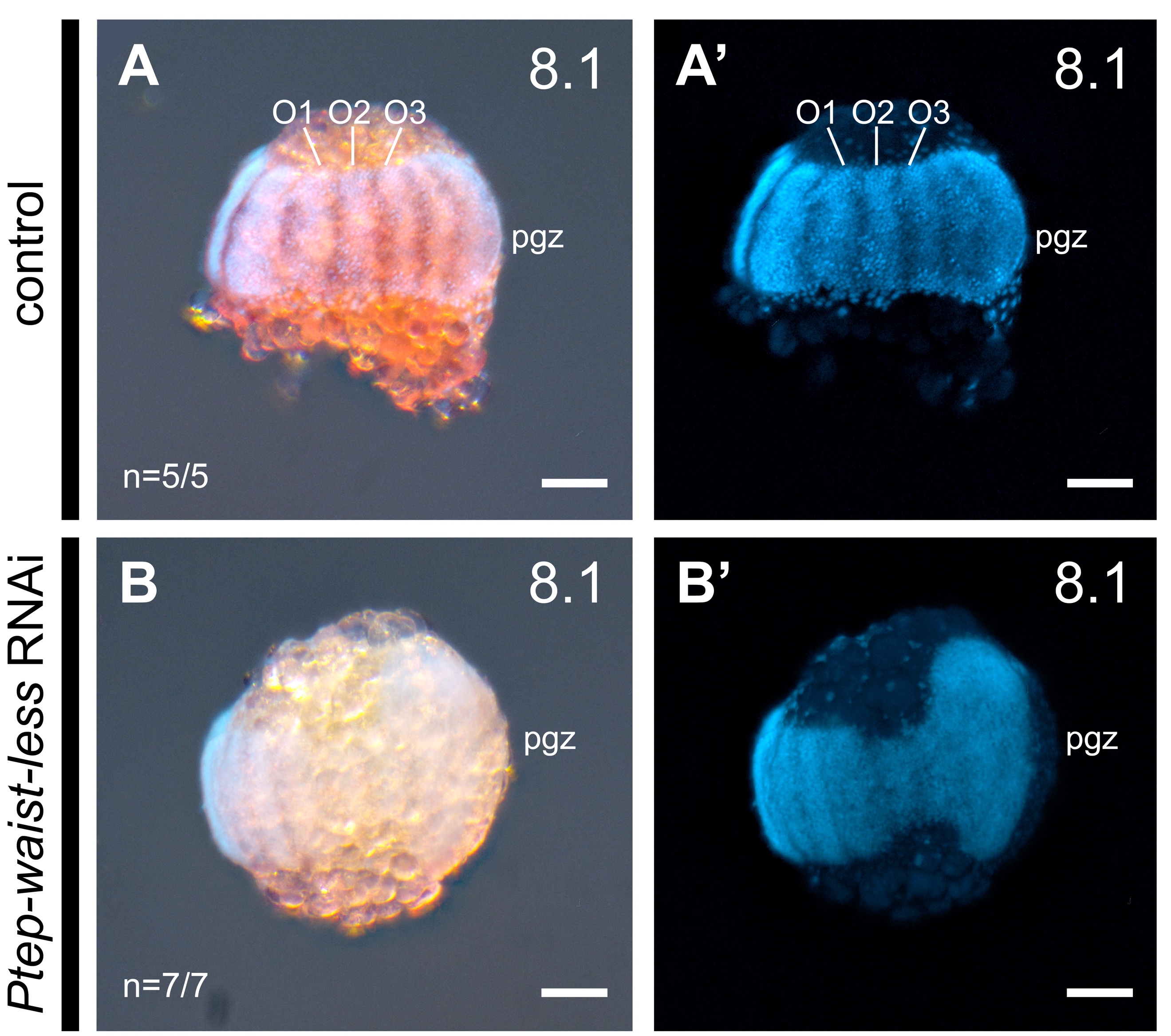

**Figure S4. Validation of RNAi using colorimetric *in situ* hybridization.** Upper row: negative control embryo showing wild type expression of *Ptep-waist-less.* Lower row: RNAi embryo showing diminution of *Ptep-waist-less* expression and abnormal development of germ band in the territory abutting the posterior growth zone. Scale bar: 100 μm.

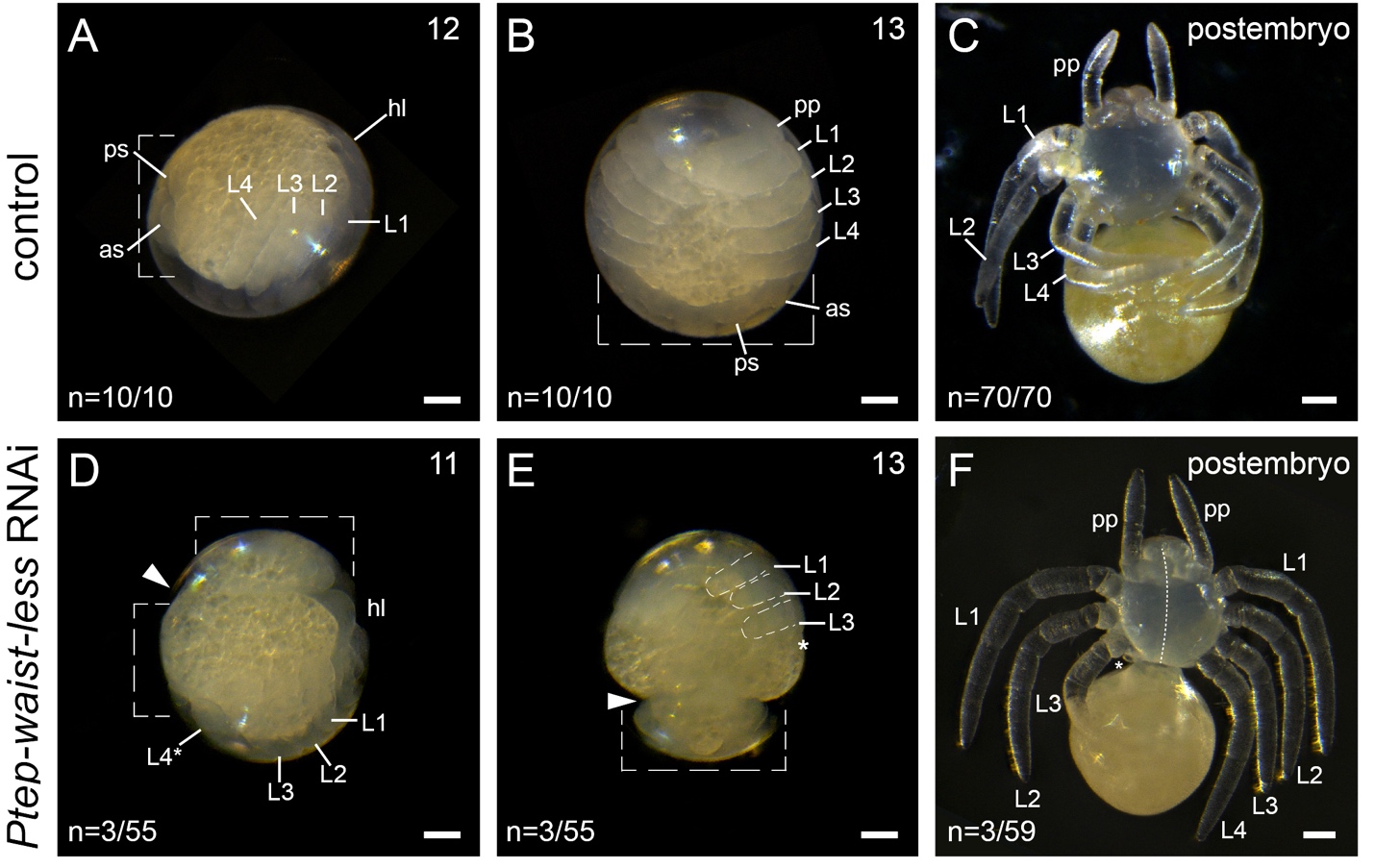

**Figure S5. Effects of *Ptep-waist-less* RNAi in late stages of embryogenesis and at hatching.** (A-C) Negative control embryos exhibiting wild type development. (D) Stage 11 Class I RNAi embryo exhibiting discontinuous germ band and aberrant disposition of opisthosoma. (E) Stage 13 Class II RNAi embryo exhibiting anomalous development of the pedicel territory and constriction of germ band due to missing tissue between tagmata. (F) Postembryo from RNAi experiment with mosaic phenotype, exhibiting loss of L4 (asterisk) and smaller prosoma on affected side (note position of dotted line in midline of the prosoma). Scale bar: 100 μm.

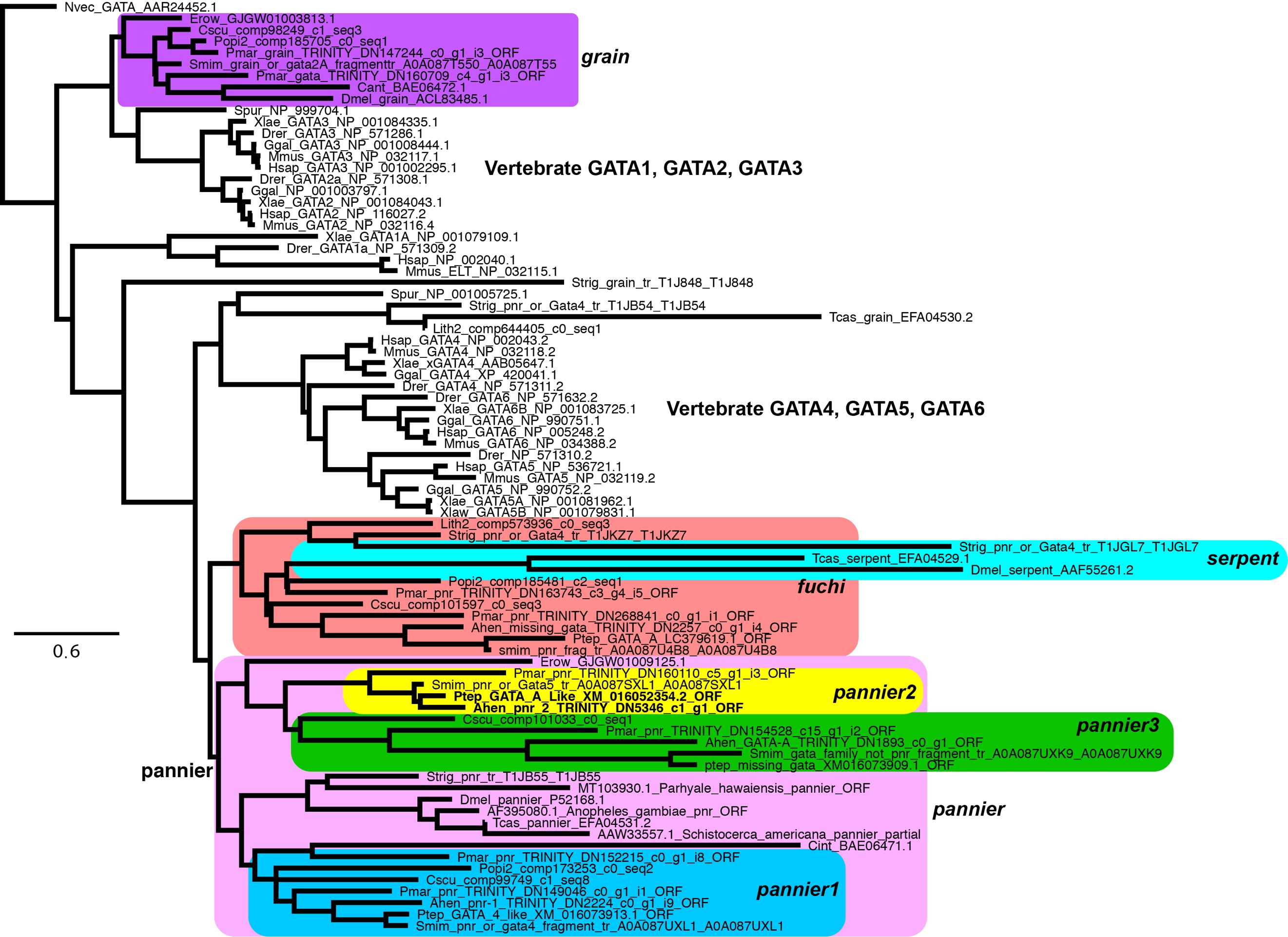

**Figure S6. Maximum likelihood tree topology of GATA homologs.** Colors correspond to previously identified paralogs of GATA. Numbers on nodes correspond to bootstrap resampling frequencies. Boldface text indicates *pannier2* copies of spiders.

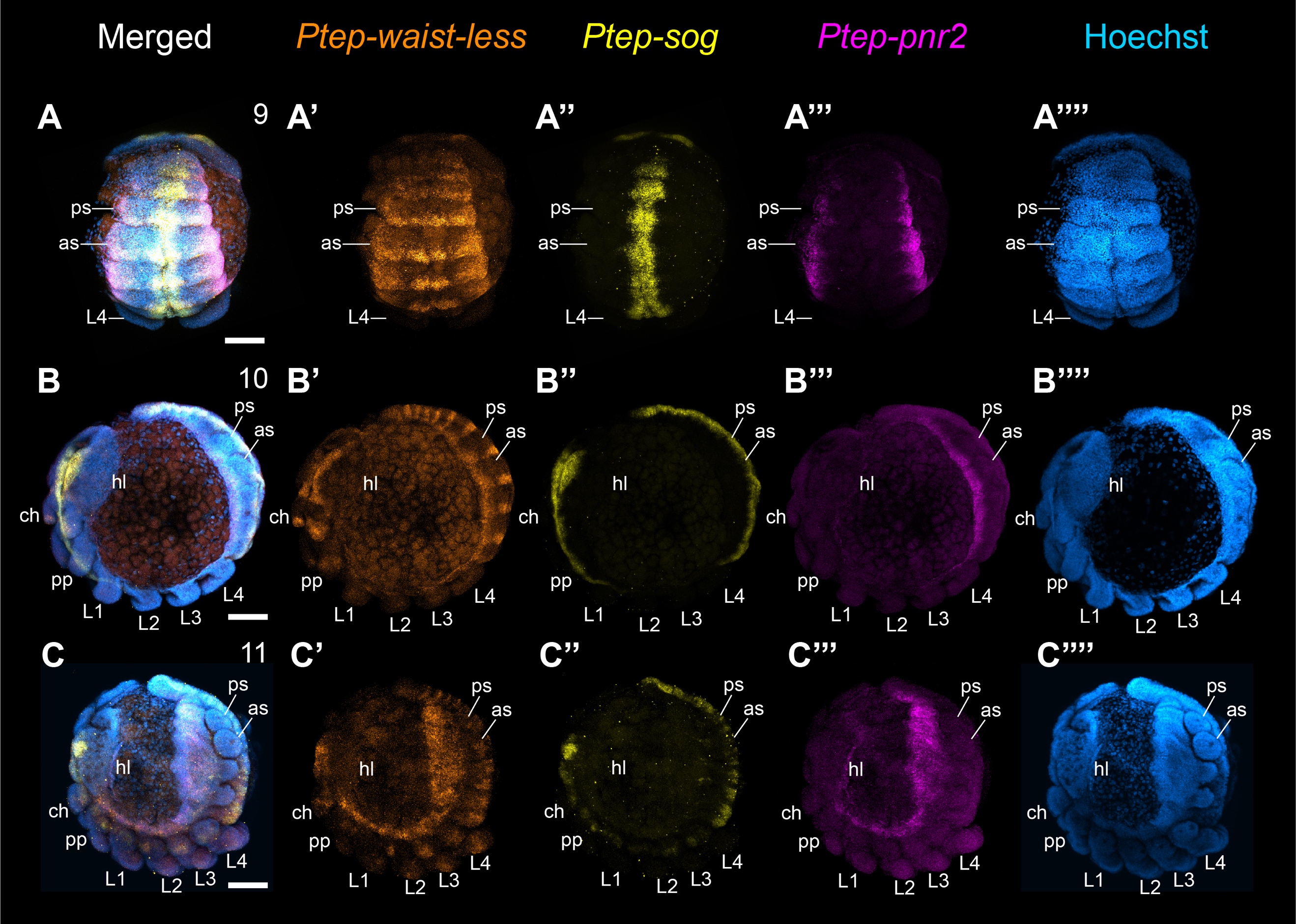

**Figure S7. Wild type expression of *Ptep-pnr2* shown in context with *Ptep-waist-less* and the ventral midline marker *Ptep-sog.*** Across all three stages surveyed for DGE in *A. hentzi*, *Ptep-pnr2* is expressed in the lateral edge of the germ band, which will become the dorsal part of the spider. In accordance with the DGE data, the strongest expression of *Ptep-pnr2* is in the opisthosoma (n=12/12). Abbreviations as in Figure 2. Scale bars: 100 μm.

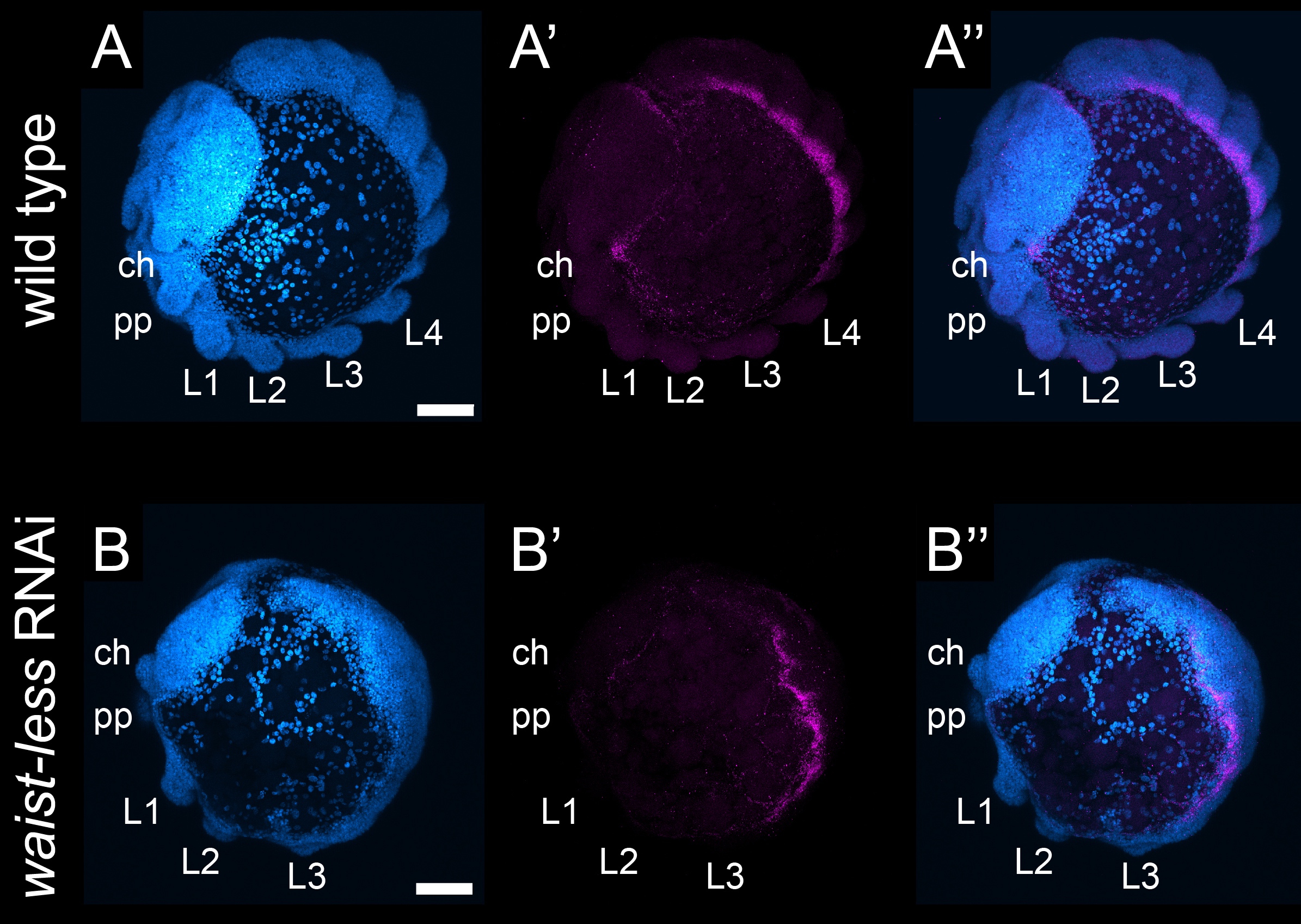

**Figure S8. Expression of *Ptep-pnr2* in a Class I *Ptep-waist-less* loss-of-function phenotype is disrupted in the opisthosoma.** In *Ptep-waist-less* RNAi embryos, the expression of *Ptep-pnr2* is no longer cleanly defined in the lateral edge of the opisthosoma and becomes blurred (n=6/9; wild type n=12/12). Abbreviations as in Figure 2. Scale bars: 100 μm.

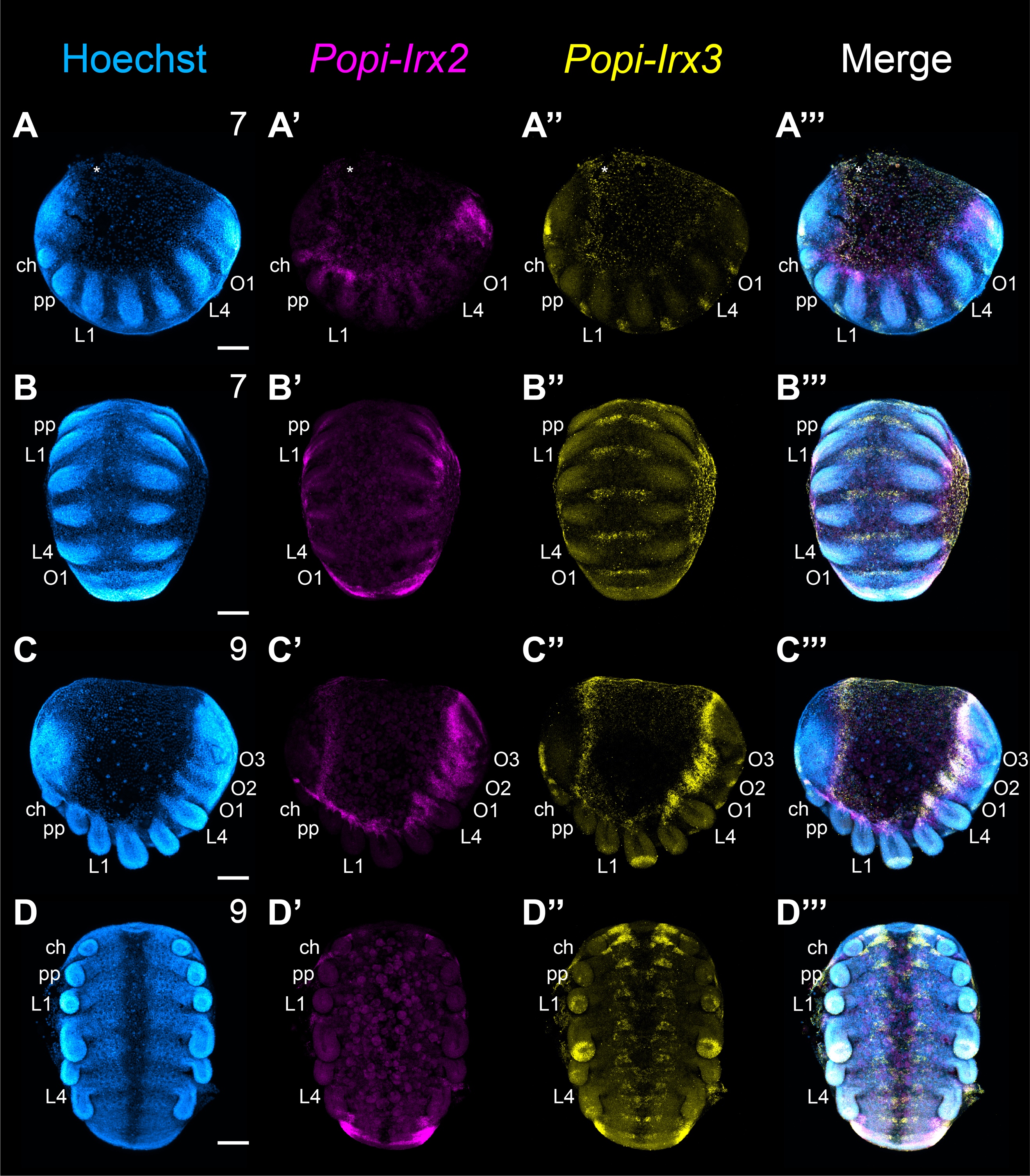

**Figure S9. Expression *Irx2* and *Irx3* in the harvestman *P. opilio*.** A-B’’’. Expression of *Popi-Irx2* and *Popi-Irx3* in stage 7 embryos. *Popi-Irx2* is expressed in the lateral margin of the germ band, with concentrated expression in the posterior terminus. Expression is not recovered in the head lobes or in the ventral ectoderm (A’, B’). *Popi-Irx3* is absent from the body wall at this stage, with expression limited to the head lobe and segmentally iterated parts of the ventral ectoderm (A’’, B’’). C-D’’’. Expression of *Popi-Irx2* and *Popi-Irx3* in stage 9 embryos. *Popi-Irx2* is restricted to the body wall and enriched in the posterior growth zone. (C’, D’). *Popi-Irx3* domains are retained in the head and ventral ectoderrm, with increased complexity of expression in the central nervous system. New regions of expression are now present in the distal portions of the prosomal appendages, with the exception of the chelicera, and an enrichment of expression in the distal L2 territory. Cheliceral expression is limited to a small area at the base of the appendage. Additionally, *Popi-Irx3* is expressed in the body wall and in a manner similar to *Ptep-waist-less*, with expression most concentrated from L4 to the posterior terminus (C’’, D’’). Asterisks in A-B’’’ mark mechanical damage to yolk. Abbreviations as in Figure 2. Scale bars: 100 μm.

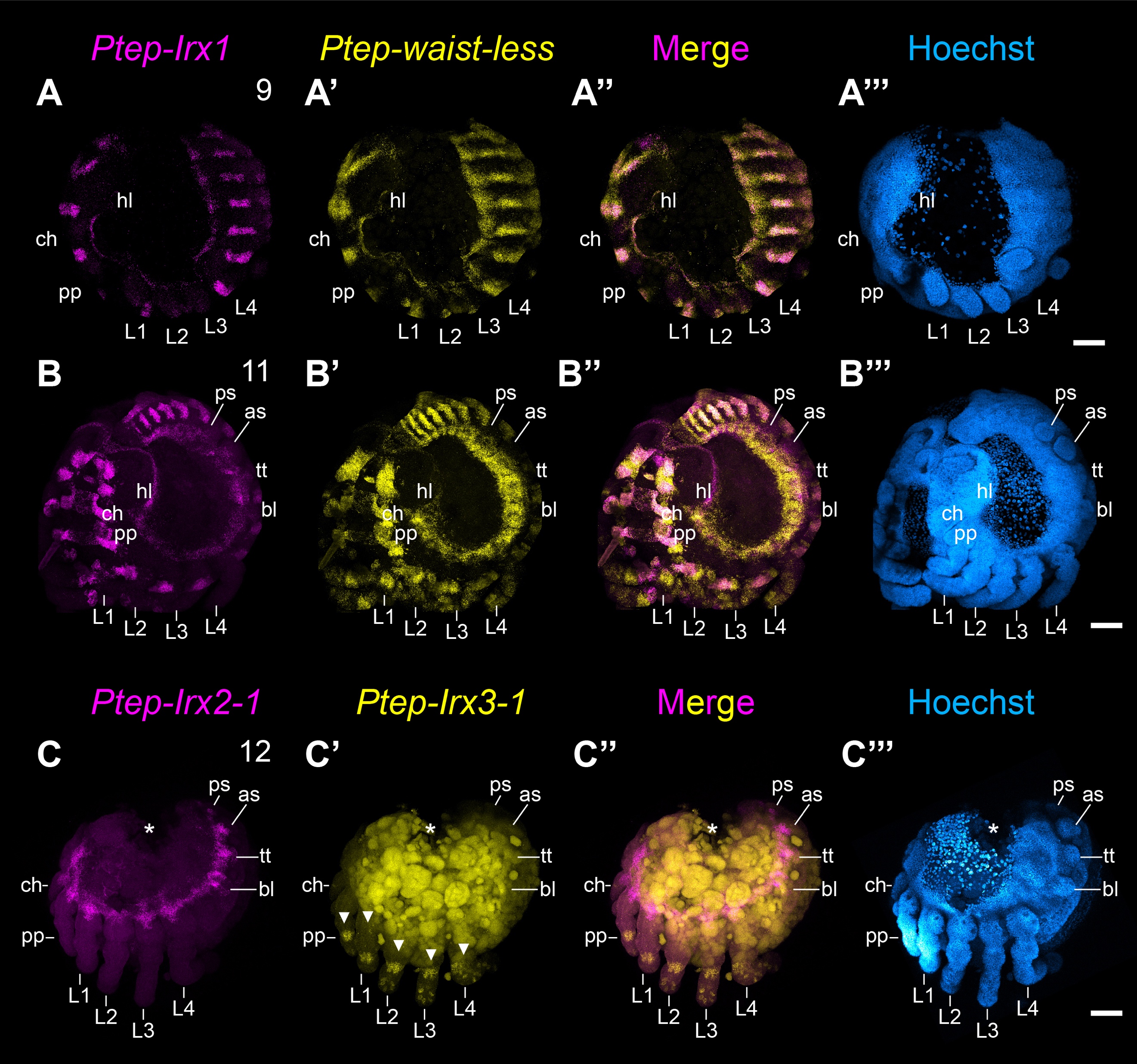

**Figure S10. Expression of *Iroquois* homologs in *P. tepidariorum*.** A-B’’’. *Ptep-Irx1* and *Ptep-waist-less* are similarly expressed but have distinct expression domains. *Ptep-waist-less* is enriched in body wall tissue of the opisthosoma, has a broader expression territory in the head and has additional expression domains in the legs, as compared to *Ptep-Irx1* (A’’, B’’). C-C’’. *Ptep-Irx2-1* and *Ptep-Irx3-1* are not comparably expressed to *Ptep-waist-less*. *Ptep-Irx2-1* is restricted to a uniform band of expression along the lateral margin of the germ band, with slight protrusions into the proximal-most regions of developing appendages (C). Expression of *Ptep-Irx3-1* is restricted to two non-overlapping expression domains in the developing legs and pedipalps, one distal and one medial. *Ptep-Irx3-1* is notably absent from the chelicera (C’). Asterisks in C-C’’’ mark mechanical damage to yolk; note autofluorescence in C’ and C’’ for *Ptep-Irx3-1*. Abbreviations as in Figure 2. Scale bars: 100 μm.

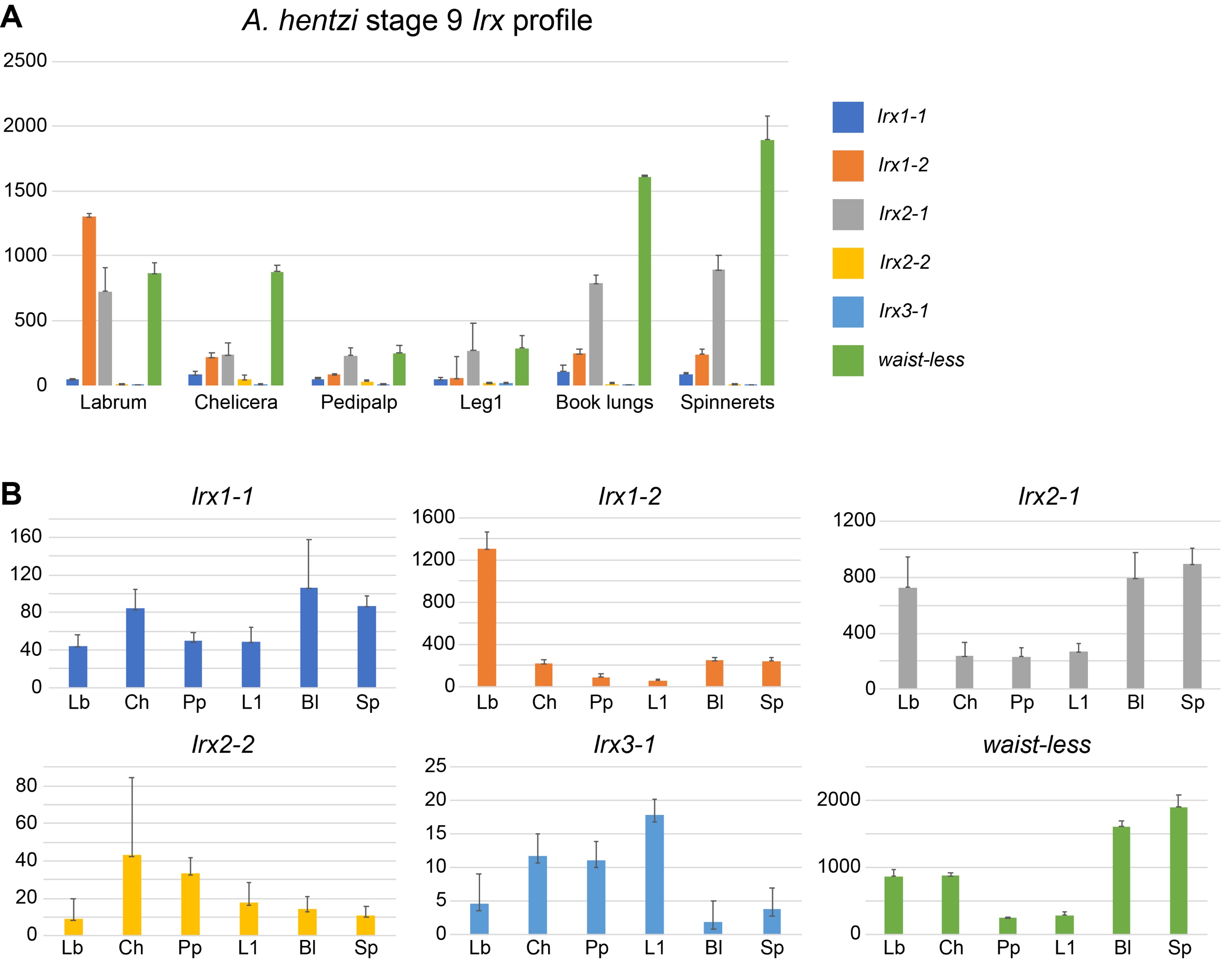

**Figure S11. Differential gene expression profile of *Iroquois* homologs in *A. hentzi* embryos at stage 9.** A. Expression levels for each *Iroquois* homolog by RNA-seq library (tissue type) in transcripts per million (TPM). B. Individual expression profiles of homologs by tissue type (magnified from panel A) show *Ahen-waist-less* is not comparably expressed to *Ahen-Irx3-*1 or other *Iroquois* homologs. Transcripts of *Ahen-waist-less* are highly enriched in RNA-seq libraries of opisthosomal tissue, to the exclusion of all prosomal regions sampled. Abbreviations: bl, book lung; ch, chelicera; lb, labrum; L1, first walking leg; pp, pedipalp; sp, spinnerets.

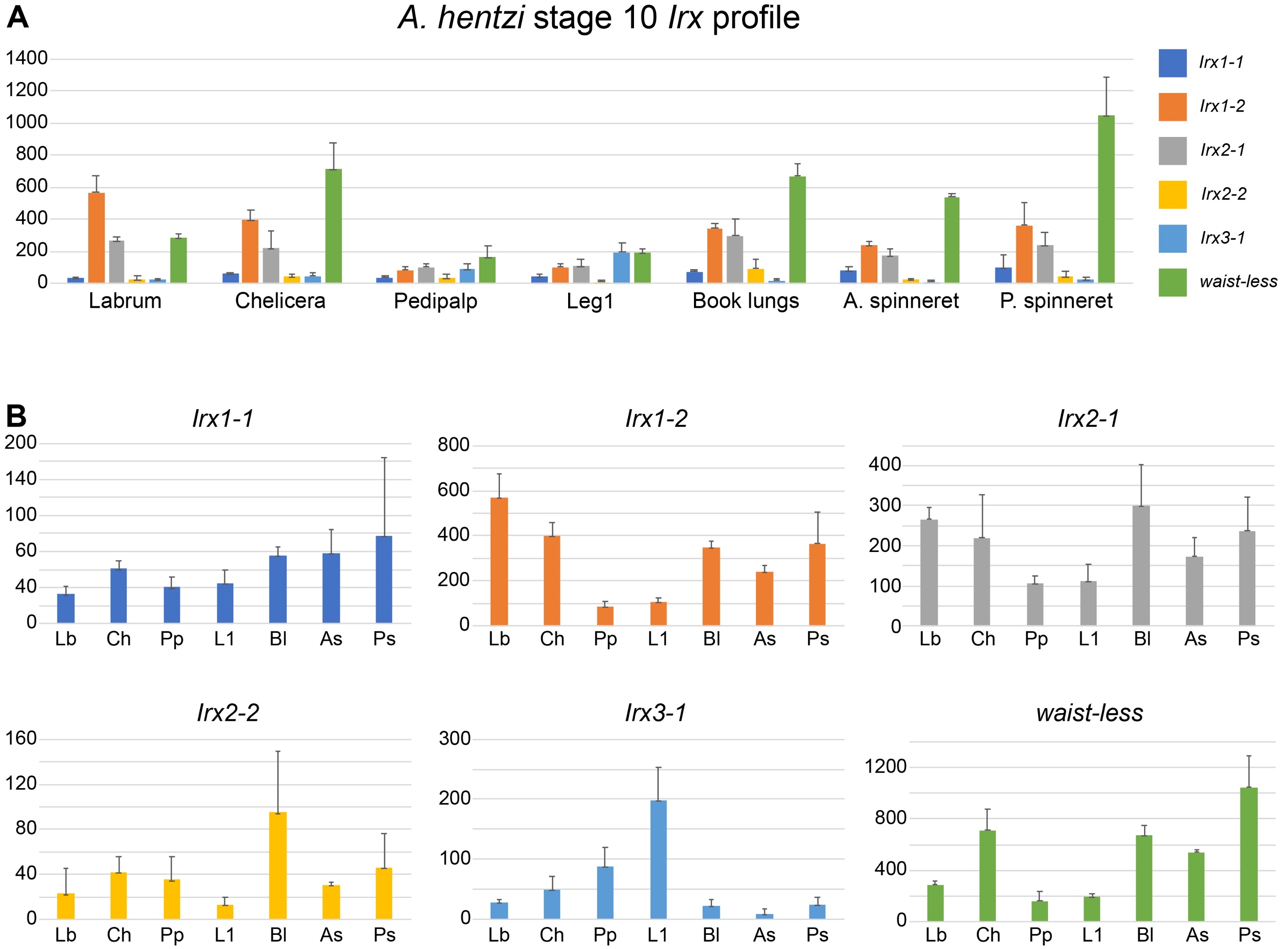

**Figure S12**. **Differential gene expression profile of *Iroquois* homologs in *A. hentzi* embryos at stage 10.** A. Expression levels for each *Iroquois* homolog by RNA-seq library (tissue type) in transcripts per million (TPM). B. Individual expression profiles of homologs by tissue type (magnified from panel A) show *Ahen-waist-less* is not comparably expressed to *Ahen-Irx3-*1 or other *Iroquois* homologs. Transcripts of *Ahen-waist-less* are enriched in RNA-seq libraries of opisthosomal tissue. Abbreviations: as, anterior spinneret; bl, book lung; ch, chelicera; lb, labrum; L1, first walking leg; pp, pedipalp; ps, posterior spinneret.

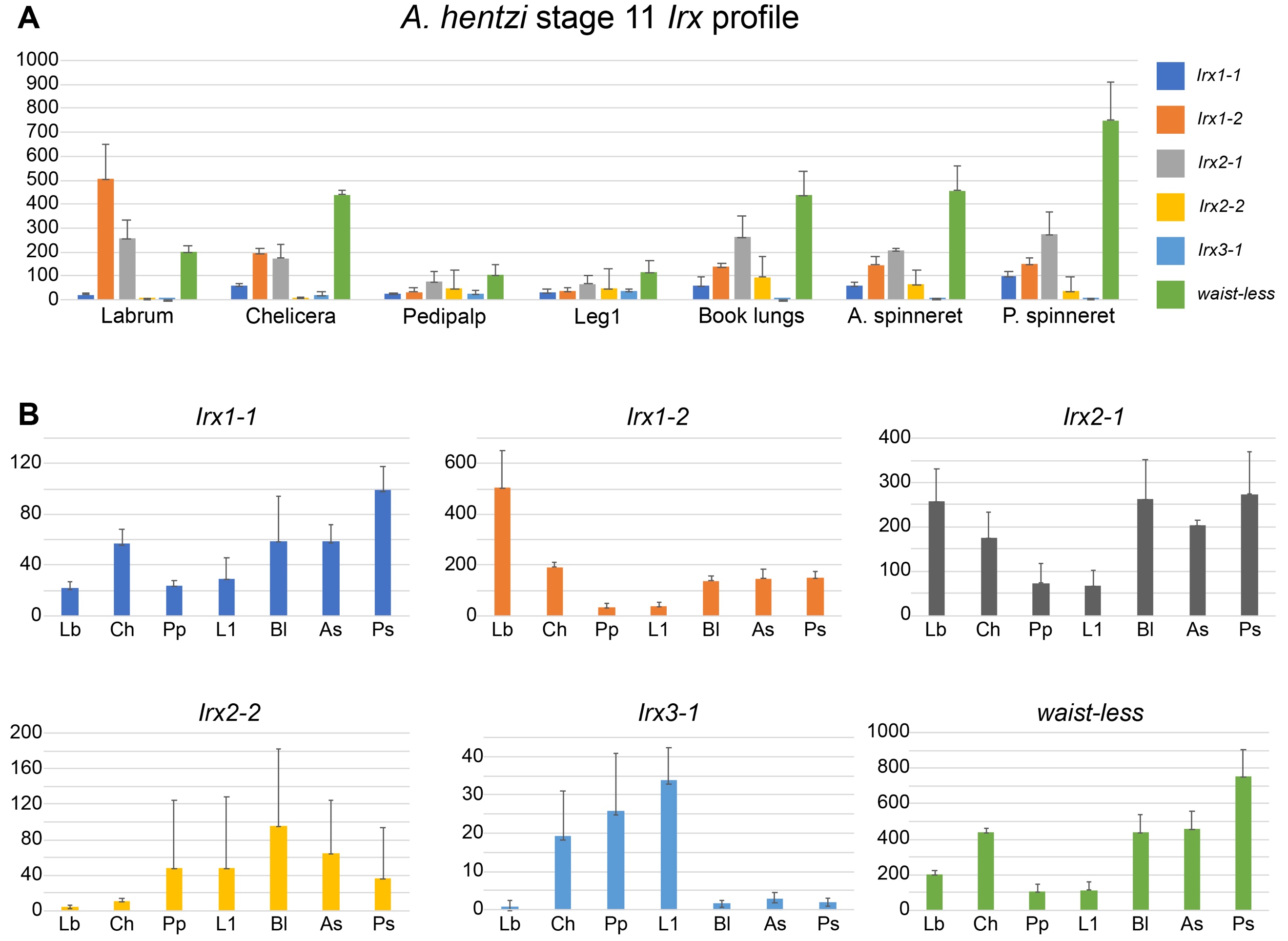

**Figure S13**. **Differential gene expression profile of *Iroquois* homologs in *A. hentzi* embryos at stage 11.** A. Expression levels for each *Iroquois* homolog by RNA-seq library (tissue type) in transcripts per million (TPM). B. Individual expression profiles of homologs by tissue type (magnified from panel A) show *Ahen-waist-less* is not comparably expressed to *Ahen-Irx3-*1 or other *Iroquois* homologs. Transcripts of *Ahen-waist-less* are enriched in RNA-seq libraries of opisthosomal tissue. Abbreviations: as, anterior spinneret; bl, book lung; ch, chelicera; lb, labrum; L1, first walking leg; pp, pedipalp; ps, posterior spinneret.
